# Engineering of *Pseudomonas putida* for accelerated co-utilization of glucose and cellobiose yields aerobic overproduction of pyruvate explained by an upgraded metabolic model

**DOI:** 10.1101/2022.07.22.501097

**Authors:** Dalimil Bujdoš, Barbora Popelářová, Daniel C. Volke, Pablo. I. Nikel, Nikolaus Sonnenschein, Pavel Dvořák

**Affiliations:** Department of Experimental Biology (Section of Microbiology), Faculty of Science, Masaryk University, Kamenice 753/5, 62500, Brno, Czech Republic; The Novo Nordisk Foundation Center for Biosustainability, Technical University of Denmark, Kemitorvet 220, 2800 Kongens Lyngby, Denmark; Department of Biotechnology and Biomedicine, Technical University of Denmark, Building 223, Søltofts Plads, 2800 Kgs. Lyngby, Denmark

**Keywords:** *Pseudomonas putida*, metabolic engineering, glucose, cellobiose, co-utilization of sugars, pyruvate, metabolic model

## Abstract

*Pseudomonas putida* KT2440 is an attractive bacterial host for biotechnological production of valuable chemicals from renewable lignocellulosic feedstocks as it can valorize lignin-derived aromatics or cellulosic glucose. *P. putida* EM42, a genome-reduced variant of *P. putida* KT2440 endowed with advantageous physiological properties, was recently engineered for growth on cellobiose, a major cellooligosaccharide product of enzymatic cellulose hydrolysis. Co-utilization of cellobiose with glucose was achieved in a mutant lacking periplasmic glucose dehydrogenase Gcd (PP_1444). However, the cause of the observed co-utilization was not understood and the Δ*gcd* strain suffered from a significant growth defect. In this study, we aimed to investigate the basis of the simultaneous uptake of the two sugars and accelerate the growth of *P. putida* EM42 Δ*gcd* mutant for the bioproduction of valuable compounds from glucose and cellobiose. We show that the *gcd* deletion abolished the inhibition of the exogenous β-glucosidase BglC from *Thermobifida fusca* by the intermediates of the periplasmic glucose oxidation pathway. The additional deletion of the *hexR* gene, which encodes a repressor of the upper glycolysis genes, failed to restore the rapid growth on glucose. The reduced growth rate of the Δ*gcd* mutant was partially compensated by the implantation of heterologous glucose (Glf from *Zymomonas mobilis*) and cellobiose (LacY from *Escherichia coli*) transporters. Remarkably, this intervention resulted in the accumulation of pyruvate in aerobic *P. putida* cultures. We demonstrated that the excess of this key metabolic intermediate can be redirected to the enhanced biosynthesis of ethanol and lactate. The overproduction of pyruvate was then unveiled by an upgraded genome-scale metabolic model constrained with proteomic and kinetic data. The model pointed to the saturation of glucose catabolism enzymes due to unregulated substrate uptake and it predicted improved bioproduction of pyruvate-derived chemicals by the engineered strain. This work sheds light on the co-metabolism of cellulosic sugars in an attractive biotechnological host and introduces a novel strategy for pyruvate overproduction in bacterial cultures under aerobic conditions.

**Highlights:**

- Co-utilization of glucose and cellobiose achieved in *P. putida* EM42 Δ*gcd* mutant.
- Growth defect of the mutant compensated by implanting exogenous sugar transporters.
- Enhanced influx of carbon caused aerobic overproduction of pyruvate and acetate.
- Carbon from excess pyruvate streamed into ethanol or L-lactate.
- Pyruvate overproduction unveiled by a mathematical model of *P. putida* metabolism.

## 1. Introduction

*Pseudomonas putida* KT2440 and its genome-reduced derivative strain EM42 with the improved expression of heterologous genes and enhanced genetic stability, viability, and thermal tolerance (Aparicio et al., 2019; Martínez-García et al., 2014b) are robust Gram-negative bacterial workhorses that attract considerable attention as new microbial platforms for lignocellulose biotechnology (Dvořák and de Lorenzo, 2018; Jayakody et al., 2018; Kohlstedt et al., 2022). *P. putida* grows well on lignin-born aromatics such as *p*-coumarate, ferulate, or benzoate, as well as on D-glucose (µ∼0.5-0.7 h^-1^), a monomeric unit of the cellulose polymer. Glucose is metabolized by this bacterium via the periplasmic oxidative route and 6-phosphogluconate or via direct phosphorylation to glucose-6-phosphate in the cytoplasm (**Figure 1**) (del Castillo et al., 2007). Both 6-phosphogluconate and glucose-6-phosphate enter the EDEMP cycle, which combines reactions from Entner-Doudoroff, Embden-Meyerhof-Parnas, and pentose phosphate pathways (Nikel et al., 2015). *P. putida*’s substrate scope was further expanded towards hemicellulosic sugars D-xylose, L-arabinose, and D-galactose by metabolic engineering (Dvořák and de Lorenzo, 2018; Elmore et al., 2020; Peabody et al., 2019) and the bacterium was demonstrated to simultaneously catabolize lignocellulosic sugars and aromatics (Dvořák and de Lorenzo, 2018; Kukurugya et al., 2019; Peabody et al., 2019). It was also empowered with surface-displayed designer protein scaffolds that can serve for the anchoring of depolymerizing enzymes (Dvořák et al., 2020a). *P. putida* also has a broad potential to valorize lignocellulosic substrates (Anna Weimer et al., 2020). The rich metabolic network of the bacterium has been tailored for the bioproduction of polyhydroxyalkanoates (Poblete-Castro et al., 2013; Salvachúa et al., 2020), rhamnolipids (Tiso et al., 2020, p. 2), *cis*,*cis*-muconic acid (Bentley et al., 2020; Kohlstedt et al., 2018), pyruvate and lactate (Johnson and Beckham, 2015), isobutanol (Ankenbauer et al., 2021), or even fluorinated building-blocks (Calero et al., 2020). These and other advantageous characteristics including HV1 safety certification (Kampers et al., 2019), resistance to oxidative stress (Elmore et al., 2020; Guarnieri et al., 2017), suitability for large-scale aerobic fermentations (Ankenbauer et al., 2020), availability of high-quality genome-scale metabolic reconstruction (Nogales et al., 2020), or amenability to genetic manipulations with an available broad palette of engineering tools (Martínez-García and de Lorenzo, 2017) predetermine *P. putida* for the biotechnological upcycling of lignocellulosic substrates.

**Figure 1.**
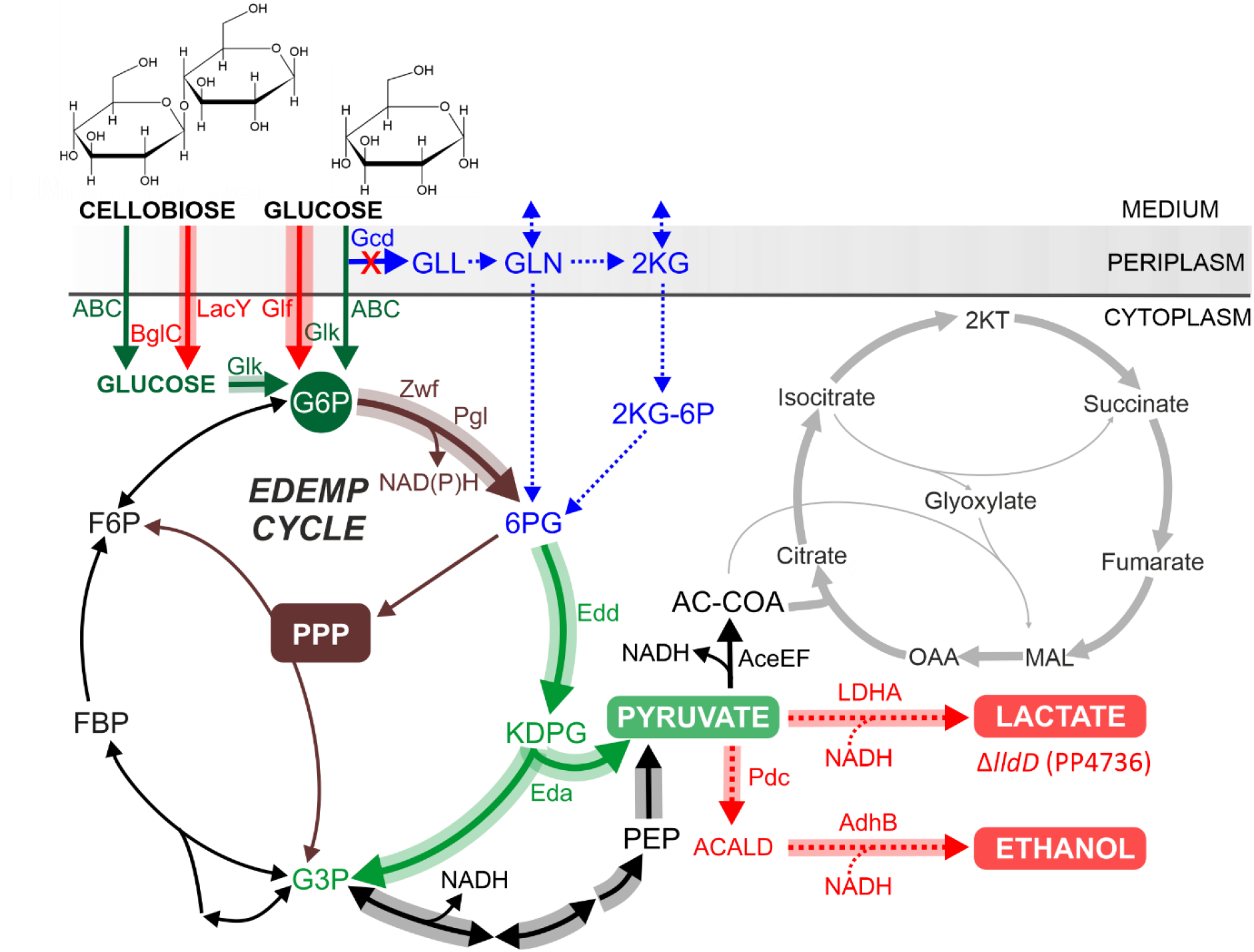
Scheme of glucose metabolism in *Pseudomonas putida* and major modifications performed in this study. *P. putida* metabolizes glucose via two major routes: (i) the periplasmic oxidative route (in blue) which starts with membrane-bound glucose dehydrogenase Gcd and (ii) the direct phosphorylation pathway (in dark green) initiated by glucokinase Glk in the cytoplasm. Products of the two pathways, 6-phosphogluconate (6PG) and glucose-6-phosphate (G6P), respectively, enter the EDEMP cycle, which combines reactions from Entner-Doudoroff (in pale green), Embden-Meyerhof-Parnas (in black), and pentose phosphate (PPP, in brown) pathways. Carbon from glucose is streamed to pyruvate and acetyl coenzyme A (AC-COA) which enters the tricarboxylic acid cycle (in grey). Cellobiose metabolism was previously established in *P. putida* by implanting cytoplasmic β-glucosidase BglC from *Thermobifida fusca* (Dvořák and de Lorenzo, 2018). Major interventions from the current study that led to the aerobic overproduction of pyruvate and derived bioproducts ethanol and lactate are shown in red. Note that the scheme shows only the reactions relevant for this work. Abbreviations: (metabolites) ACALD, acetaldehyde; FBP, fructose 1,6-biphosphate; F6P, fructose 6-phosphate; GLL, glucono-δ-lactone; GLN, gluconate; G3P glycerol 3-phosphate; KDPG, 2-keto-3-deoxy-6-phosphogluconate; 2KG, 2-ketogluconate; 2KG-6P, 2-ketogluconate-6-phosphate; 2KT, α-ketoglutarate; MAL, malate; NADH, reduced nicotinamide adenine dinucleotide; NADPH, reduced nicotinamide adenine dinucleotide phosphate; OAA, oxaloacetate; PEP, phosphoenolpyruvate; (enzymes and transporters) ABC, mannose/glucose ABC transporter; AdhB, alcohol dehydrogenase (*Zymomonas mobilis*); Eda, 2-dehydro-3-deoxy-phosphogluconate aldolase; Edd, phosphogluconate dehydratase; Glf, glucose facilitated diffusion protein (*Zymomonas mobilis*); LacY, lactose permease (*Escherichia coli*); LDHA, L-lactate dehydrogenase with N109G mutation (*Bos taurus*); lldD, L-lactate dehydrogenase; Pgl, 6-phosphogluconolactonase; Pdc, pyruvate decarboxylase (*Zymomonas mobilis*); Zwf, glucose 6-phosphate 1-dehydrogenase.

One remaining limitation was *P. putida*’s lack of growth on cellooligosaccharides (Dvořák and de Lorenzo, 2018). Cellooligosaccharides are formed as major by-products of incomplete cellulose saccharification due to the high crystallinity of the polymer and rate-limiting cellobiose hydrolysis by fungal β-glucosidases employed in enzyme cocktails (Barbosa et al., 2020; Ferreira et al., 2018; Parisutham et al., 2017). In corn stover hydrolysates, for instance, glucose can make roughly 50 % and cellooligomers form up to 16 % of all fermentable sugars (Elmore et al., 2020). In sugarcane straw hydrolysate prepared by commercial enzyme cocktails, the ratio of cellooligosaccharides to glucose ranged from 1:8 to 1:11 and cellobiose was a predominant glucose oligomer (Barbosa et al., 2020). If partial enzymatic conversion of cellulose is employed, this ratio can be modified substantially in favor of oligosaccharides (Barbosa et al., 2020). Partial cellulose hydrolysis and intracellular assimilation of resulting oligomers emerge as an alternative strategy to standard hydrolysis protocols aiming at maximized glucose yields (Oh and Jin, 2020; Parisutham et al., 2017). The adoption of microorganisms capable of efficient processing of both glucose and cellooligosaccharides is thus a rational option that can reduce the capital costs of lignocellulose biotechnologies (Parisutham et al., 2017; Taha et al., 2016; Ylinen et al., 2022; Zhang et al., 2022). In our previous work, we enriched the EM42 strain with *bglC* gene which encodes cytoplasmic β-glucosidase from *Thermobifida fusca* (Dvořák and de Lorenzo, 2018). The resulting recombinant grew well on cellobiose (µ∼0.35 h^-1^) and was also able to convert this disaccharide into valuable polyhydroxyalkanoates (Dvořák et al., 2020b). However, *P. putida bglC*^+^ preferred glucose over cellobiose in a 1:1 mixture of the two sugars and exhibited diauxic growth (Dvořák and de Lorenzo, 2018). We identified that *P. putida* mutant with deleted *gcd* gene (**Figure 1**) and closed periplasmic glucose uptake route can co-utilize glucose with cellobiose (unpublished data). The Δ*gcd* mutant is frequently used in studies that aim at utilization and valorization of glucose or xylose by *P. putida*, because it does not accumulate oxidized sugar acids gluconate or xylonate, respectively (Bentley et al., 2020; Dvořák and de Lorenzo, 2018; Elmore et al., 2020; Poblete-Castro et al., 2013) Regrettably, Δ*gcd* mutant shows a significantly reduced growth rate which narrows its utility for the bioproduction of valuable chemicals from lignocellulosic substrates.

In the present work, we aimed to elucidate the cause of glucose and cellobiose co-utilization in *P. putida* EM42 Δ*gcd* mutant and to improve its growth for enhanced bioproduction of valuable chemicals from cellulosic sugars. We hypothesized that: (i) the sequential utilization of glucose and cellobiose was caused by an inhibition of exogenous β-glucosidases rather than by a dedicated carbon catabolite repression mechanism, and (ii) the growth defect of the Δ*gcd* mutant was attributed, at least partially, to the insufficient transport of glucose and cellobiose into the cell. Kinetic analyses with BglC and metabolites from the periplasmic oxidative pathway indeed showed that the glucose preference in *P. putida gcd*^+^ stems from the inhibition of β-glucosidase. The growth rate of the Δ*gcd* mutant was then improved by introducing heterologous glucose and cellobiose transporters. Strikingly, the augmented transport of glucose and cellobiose resulted in the accumulation of pyruvate and, to a minor extent, also of acetate in aerobic *P. putida* cultures. The recombinant strain bestowed with exogenous biosynthetic modules could re-direct excess pyruvate toward the synthesis of desirable biochemicals. The accumulation of pyruvate and derived bioproducts in the mutant was then elucidated by an upgraded version of the latest genome-scale metabolic reconstruction of *P. putida* KT2440 constrained with enzymes’ abundances and kinetic parameters. This work clarifies the prerequisites of cellobiose and glucose co-metabolism in *P. putida* and introduces a novel strategy for pyruvate overproduction in aerobic bacterial cultures via augmented substrate influx and enzyme saturation. The verified metabolic model can be used as a new predictive tool in future metabolic engineering studies aimed at *P. putida*.

## 2. Materials and methods

### 2.1 Bacterial strains, plasmids, and culture conditions

All bacterial strains and plasmids used in this study are listed in **Table 1**. The workflow of the preparation of all *P. putida* mutants described in this work is outlined in **Scheme 1**. All precultures were inoculated directly from glycerol stocks stored at −80 °C. The concentrations of all antibiotics (ATB) used in the cultures of *E. coli* and *P. putida* are shown in **Table S1 in Supplementary material**. Minimal M9 medium [7 g/L Na_2_HPO_4_·7H_2_O, 3 g/L KH_2_PO_4_, 0.5 g/L NaCl, 1 g/L NH_4_Cl_2_, 2 mM MgSO_4_, 100 µM CaCl_2_, 20 µM FeSO_4_, and 3ml/L of trace elements described in (Abril et al., 1989)] was used for all *P. cultures* cultures. Unless otherwise stated, all *E. coli* strains were cultured in lysogeny broth (LB; 10 g/L tryptone, 5 g/L yeast extract, 5 g/L NaCl – in case of solid media 15g/L agar) with agitation (233 rpm, shaking incubator NB205, N-Biotek) at 37 °C. *P. putida* cultures in LB medium occurred in the same conditions except for temperature which was 30 °C.

**Table 1.**
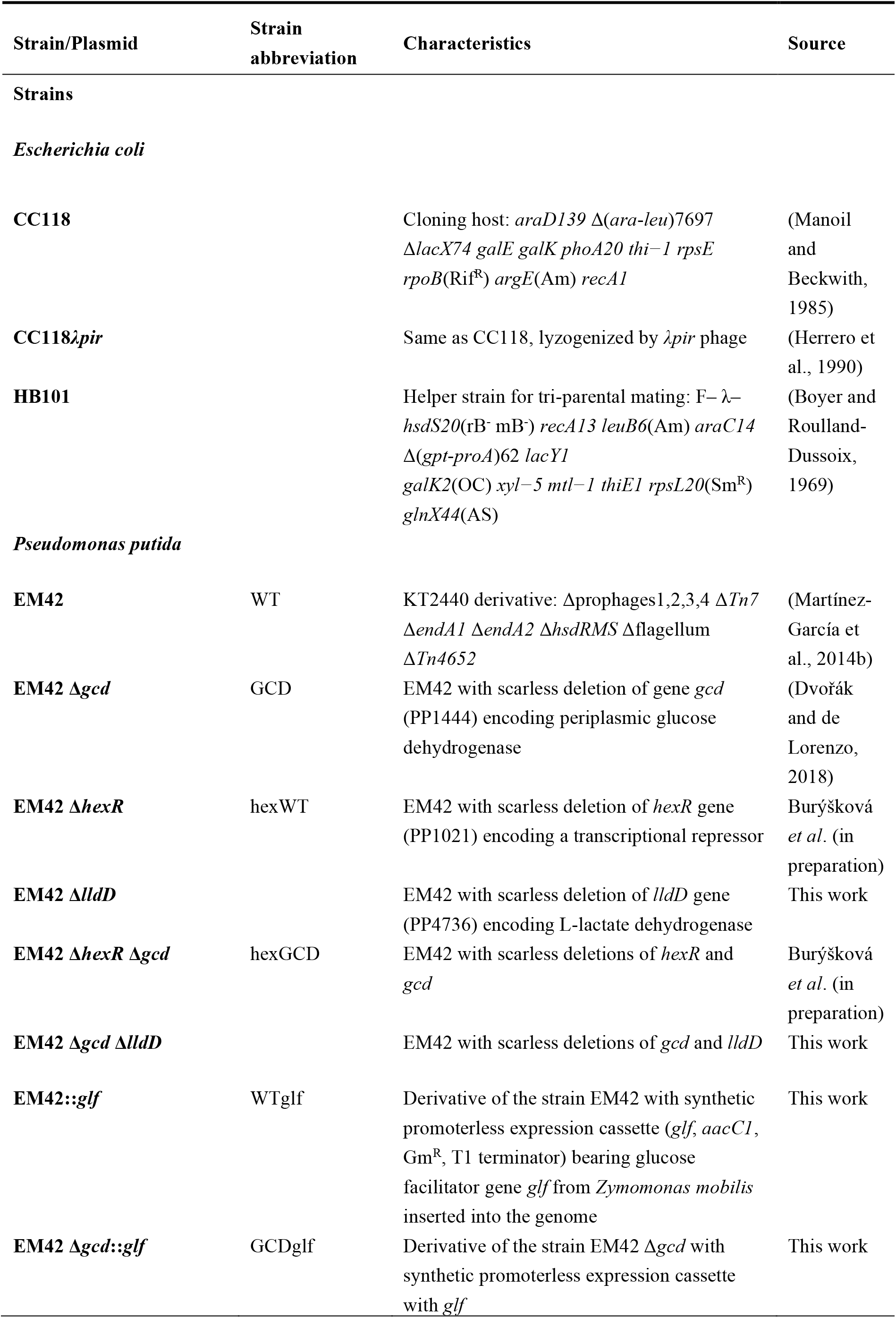
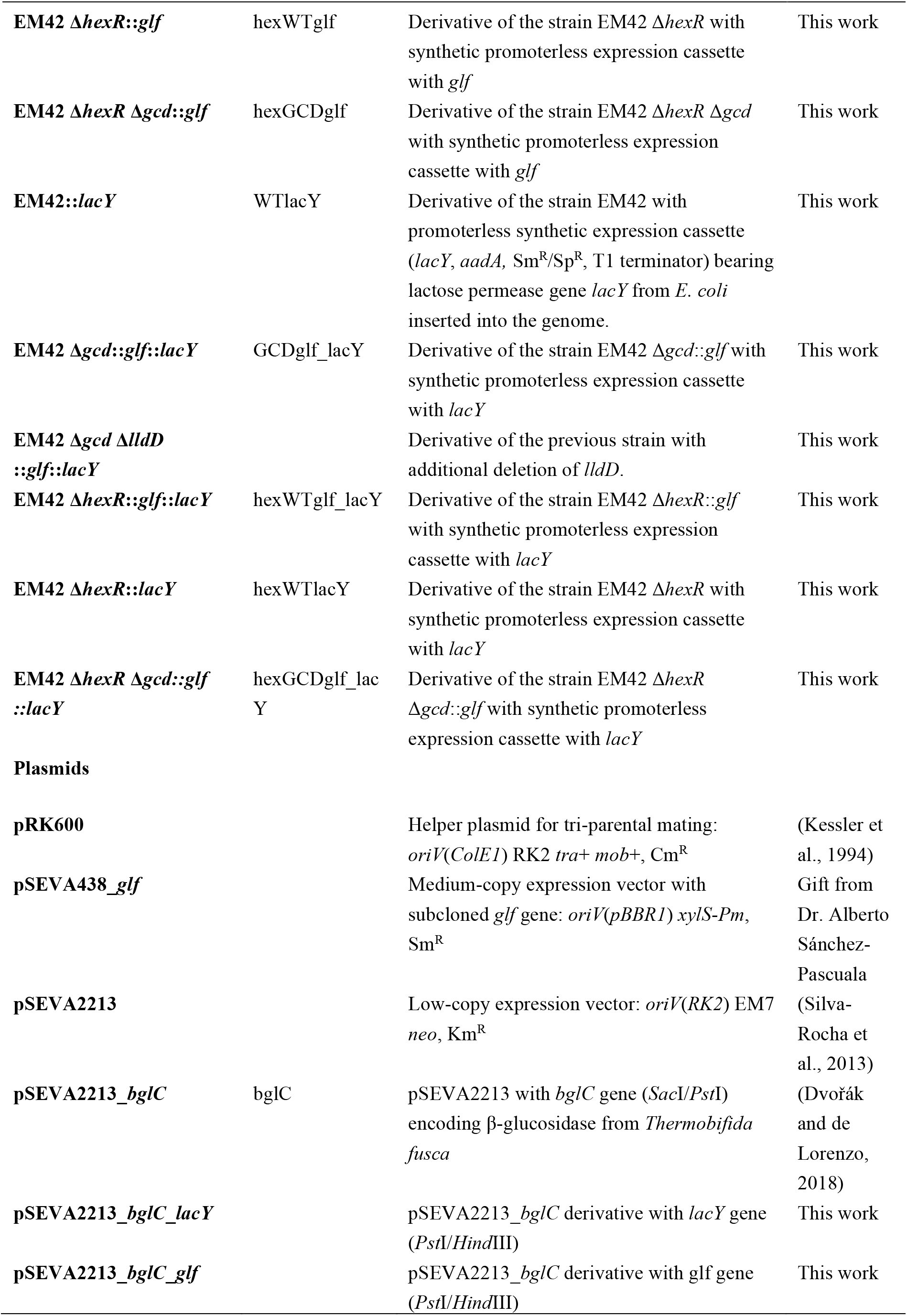
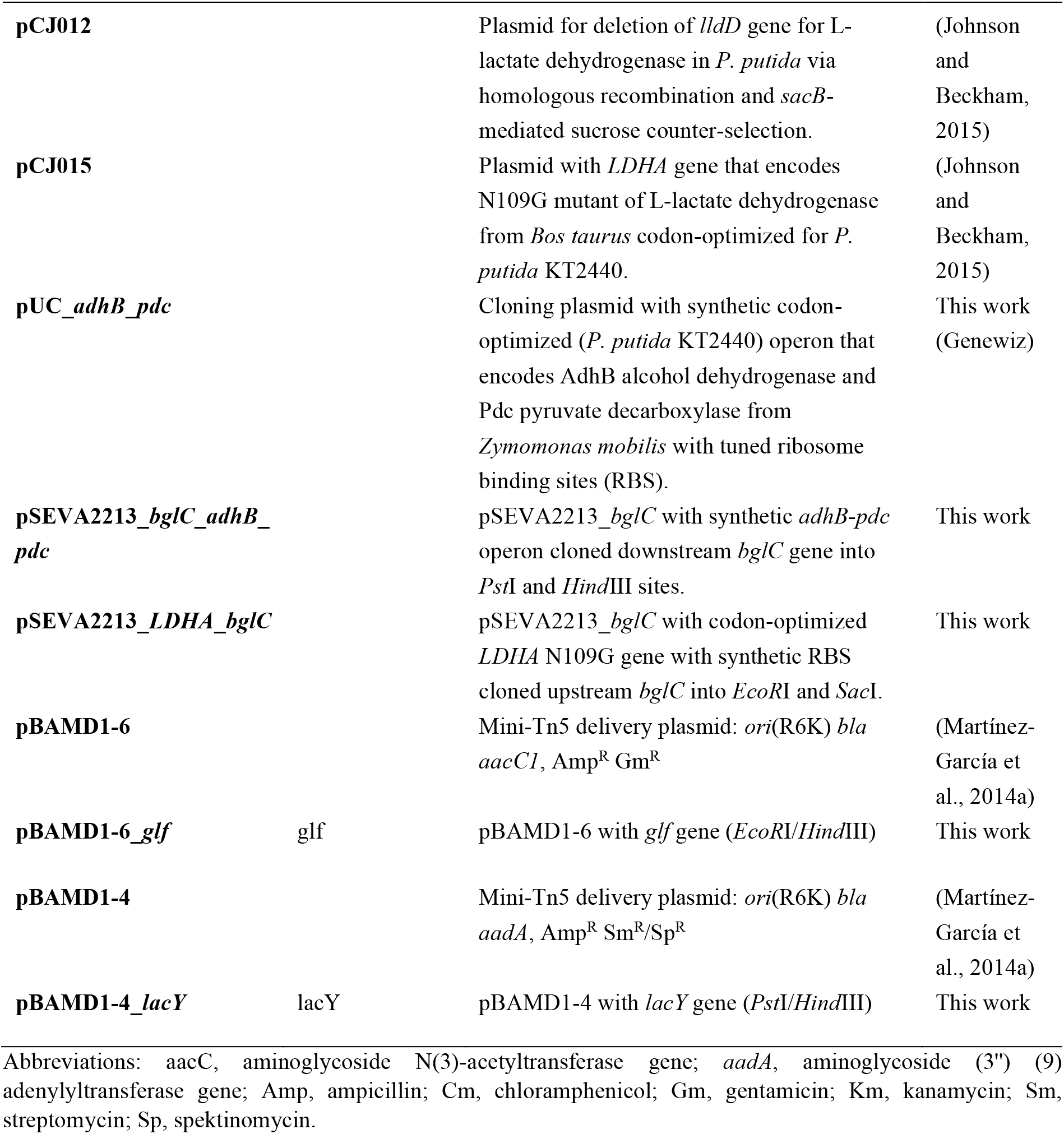
Strains and plasmids used in this work.

**Scheme 1.**
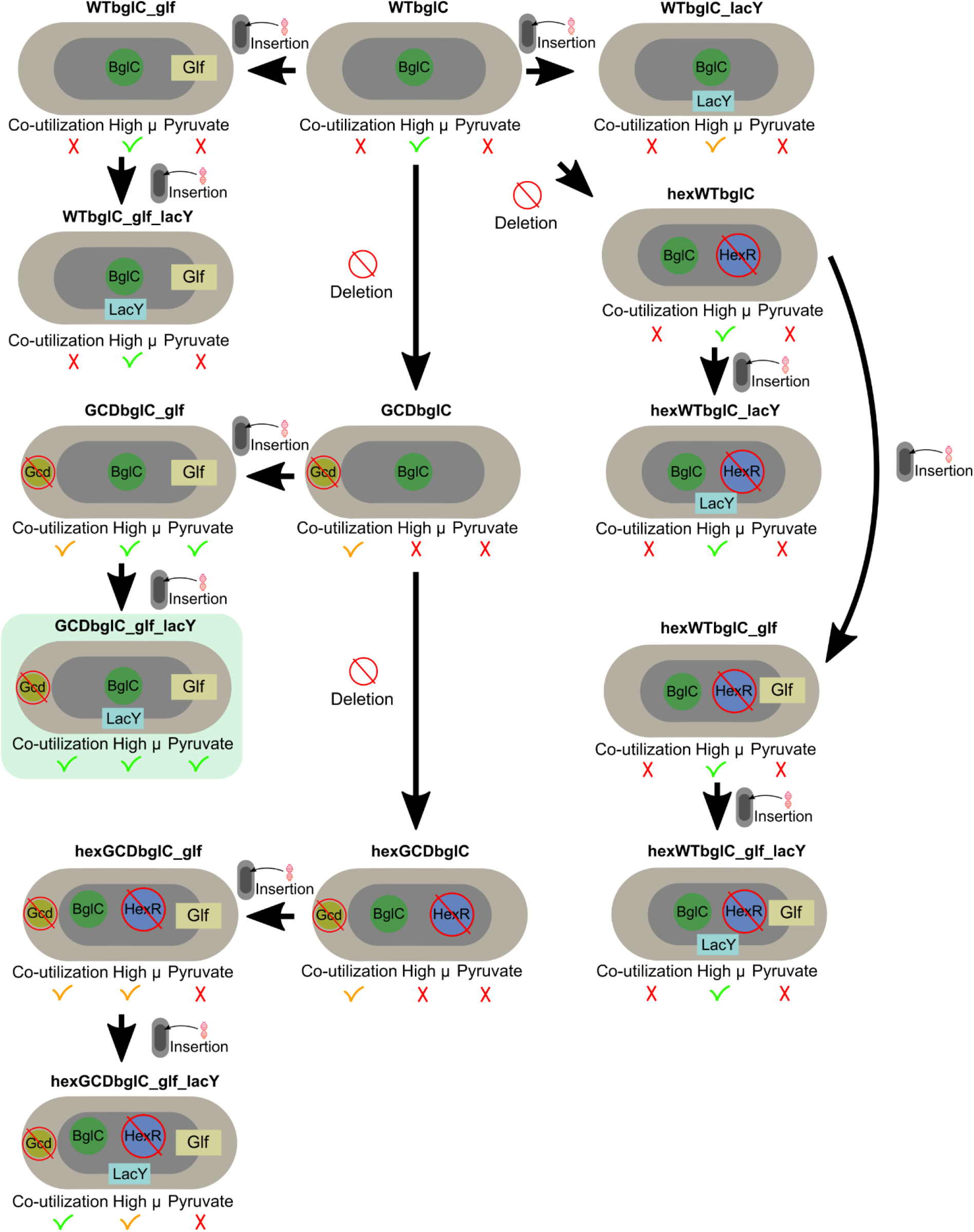
Workflow of the preparation of *Pseudomonas putida* recombinant strains described in this study. Abbreviations: BglC, β-glucosidase from *Thermobifida fusca*; Gcd, glucose dehzdrogenase (PP1444); Glf, glucose facilitator from *Zymomonas mobilis*; HexR DNA-binding transcriptional regulator (PP1021), LacY, lactose permease from *Escherichia coli*. Co-utilization signs the abbility of the given strain to utilize simultaneously glucose and cellobiose, High µ signs the specific growth rate on the mixture of glucose and cellobiose, and Pyruvate signs the abbility of the given strain to overproduce pyruvate in aerobic cell culture.

Prior to the culture in shake flasks, *P. putida* strains were precultured overnight in 2.5 ml of M9 medium with 5 g/L of glucose as a sole carbon source and an appropriate ATB. After overnight culture, cells were harvested by centrifugation (2,500 g, 10 min). Pellet was then resuspended in 1 mL of M9 medium and flasks were inoculated to the starting OD_600_ of 0.1 and incubated at 30 °C with agitation (233 RPM for cultures with substrate concentration 4 g/L and 275 RPM for cultures with substrate concentration 8 g/L) using NB205 orbital incubator (N-Biotek). The ratio of medium to flask volume was 1:5 (20 mL:100 mL) for cultures with 4g/L substrate and 1:10 (25 mL:250 mL) for cultures with 8g/L substrate. Filling of 10 mL of M9 medium in 100 mL shake flask was used for L-lactate production cultures. The exception from this protocol was the cultures performed for ethanol production in *P. putida* strains. Overnight cultures were conducted in 2.5 mL LB medium with kanamycin. Cells were then harvested by centrifugation (2,500 g, 10 min), resuspended in 1 mL of M9 medium, and 50 mL falcon tubes with 10 mL M9 medium and glucose and cellobiose (4 g/L each) were inoculated to the starting OD_600_ of 0.1 and incubated at 30 °C with agitation (275 RPM) using NB205 orbital incubator (N-Biotek). The tubes were tightly closed with a cap and sealed with parafilm to prevent evaporation of formed ethanol.

Small-scale cultures in 96- or 48-well plates were used for screening purposes. For 96-well plate cultures, strains were precultured only once. Overnight cultures were grown in 2.5 mL of LB medium with kanamycin (233 RPM, 30 °C, orbital incubator NB205). Grown cultures were harvested by centrifugation (3,500g, 10 min) and resuspended in a pure M9 medium with kanamycin and a carbon source. Starting OD_600_ was 0.05 (as measured in a cuvette with an optical path of 1 cm) in the final volume of 150 µL. The plate was placed into Infinite 200 PRO plate reader (Tecan) and incubated at 30 °C. OD was measured every 15 minutes and before the measurement, the plate was quickly shaken.

For cultures in 48-well plates, the strains were first inoculated into 200 μL of LB medium in a 2 ml Eppendorf tube and precultured for 20-22 h. Then, 100 μL of grown cells in LB medium were transferred to 10 mL of M9 medium (flask volume 50 mL) with glucose (5 g/L), and the cells were cultured for 14-17 h at 30 °C with agitation (200 RPM) using orbital incubator IS-971 (Jeio Tech). The culture was then started by adding the cells to the initial OD_600_ of 0.05 (as measured in a cuvette with an optical path of 1 cm) in 600 μL of medium with ATB and a substrate. The plate was placed in Infinite 200 PRO plate reader (Tecan) and incubated at 30°C. The OD_600_ was measured every 15 minutes and orbital shaking with 2.5 μm amplitude was applied in between measurements, linear shaking with 2.5 μm amplitude was performed 10 s before each OD measurement. To avoid precipitation of water on the plate lid, triton solution in ethanol (0.05% v/v Triton X-100 in 20% v/v EtOH) was poured into the lid so that it covered the whole surface (Brewster, 2003). The lid was dried and then sterilized by UV light.

### 2.2 General cloning procedures, construction of plasmids and mutant strains

Plasmid DNA was routinely isolated using GeneJET Plasmid Miniprep Kit (Thermo Fisher Scientific) or QIAprep Spin Miniprep Kit (Qiagen). Genomic DNA was isolated using GenElute bacterial genomic DNA kit (Sigma-Aldrich, USA). DNA was purified from PCR mixtures or agarose gels using NucleoSpin Gel and PCR Clean-up (Macherey-Nagel). The purity and concentration of DNA were checked by NanoDrop 2000 (Thermo Fisher Scientific). If needed, DNA was concentrated using DNA 120 SpeedVac Concentrator (Thermo Fisher Scientific). Plasmid constructs were sequenced by Eurofins Scientific or SEQme Czech Republic. The oligonucleotide primers used in this study (**Table S2 in Supplementary material**) were purchased from Merck. Chemocompetent *E. coli* CC118 cells were transformed with ligation mixtures or plasmid constructs and individual clones selected on LB agar plates with an antibiotic were used for the preparation of glycerol (20% w/v) stocks.

Constructed plasmids were transferred from *E. coli* CC118 donor to *P. putida* EM42 by triparental mating, using *E. coli* HB101 helper strain with pRK600 plasmid (**Table 1**). Alternatively, electroporation was used for the transformation of *P. putida* cells with selected plasmids using 2 mm gap cuvette (Thermo Fisher Scientific) and GenePulser XcellTM (Bio-Rad). Parameters of the electroporation were: voltage of 2500 V, the capacitance of 25 µF, and resistance of 200 Ω. Preparation of *P. putida* electrocompetent cells and electroporation procedure were performed as described elsewhere (Aparicio et al., 2015). After the electric pulse, 0.9 mL of LB medium was added in the cuvette and the mixture was transferred to a 2 mL plastic tube. Cells were incubated for 2 h in case of electroporating pSEVA2213_*bglC* and 5 h in case of pBAMD plasmids (170 RPM, 30 °C, NB205 incubator). After the initial cultivation, 10 µL of the culture and 90 µL of fresh LB medium were mixed and plated on a selective LB plate with ATB. The plate was left at 30 °C overnight. Single colonies were then re-streaked twice on fresh LB plates with ATB and checked for the presence of plasmid by restriction analysis.

All gene amplifications were performed using polymerase chain reaction (PCR) in a 50 µL mix with Q5 High-Fidelity DNA Polymerase (New England Biolabs) according to the manufactureŕs protocol. Colony PCR with NZYTaq II 2x Green Master Mix (Nzytech) or DreamTaq 2x Green PCR Master Mix (Thermo Fisher Scientific) was routinely run in a 10 µL mix for strain verification and confirmation of the presence of insert in a plasmid according to the manufacturer’s instructions. Forward and reverse primers were added to the final concentration of 0.5 µM and all PCR reactions were carried out in Labcycler Gradient (SensoQuest). The PCR products were analyzed by agarose gel electrophoresis with 0.8 % gels. PCR products were visualized using G:Box XT4 Digital Imaging System (Syngene) and bands were compared to the EZ Load 500 bp Molecular Ruler (Bio-Rad). All ligation reactions were performed in a 10 µL mix with T4 DNA Ligase (New England Biolabs) at 16°C overnight according to the manufactureŕs instructions.

#### Construction of plasmids pSEVA2213_bglC_lacY and pBAMD1-4_lacY

The gene *lacY* which encodes lactose permease was cloned from the genome of *E. coli* BL21(DE3). The gene was amplified from chromosomal DNA by PCR. DNA was obtained by disrupting *E. coli* cells from a couple of colonies dissolved in 20 µL of water with heating for five min. at 85 °C. The cell lysate was spun for two min. at 3,000 g and 1 µL of the supernatant was used in PCR mix with Q5 High-Fidelity DNA Polymerase using primers lacY for PstI RBS and lacY rev HindIII (**Table S2**). Annealing temperature T_a_ was 65 °C. PCR product with consensus ribosome binding site (RBS) AAGGAG introduced upstream the gene was purified and digested with *Pst*I and *Hind*III (New England Biolabs). Digested fragment was ligated into pSEVA2213_*bglC* plasmid cut with the same restriction endonucleases. Chemocompetent cells of *E. coli* CC118 were transformed with the ligation mixture and cells were plated on LB agar with kanamycin and incubated at 37 °C overnight. Transformants were checked for the presence of the plasmid by colony PCR and restriction analysis of the isolated plasmid. The sequence of the cloned *lacY* gene was eventually verified by sequencing using primers Glf seq fw (anneals to the *bglC* gene) and PS2new (**Table S2**). Gene *lacY* with synthetic RBS was then subcloned from pSEVA2213_*blgC*_*lacY* into the suicide plasmid pBAMD1-4 using *Pst*I and *Hind*IIII restriction enzymes. *E. coli* CC118λpir was transformed with the resulting plasmid. The pBAMD1-4 carries resistance markers for streptomycin and ampicillin. Thus, *E. coli* CC118λpir pBAMD1-4_*lacY* was at all times cultured with both antibiotics. Transformants were checked by colony PCR and restriction analysis.

#### Construction of plasmids with glf

The gene *glf* that encodes glucose facilitator from *Zymomonas mobilis* was subcloned with consensus RBS AGGAGG was from the plasmid pSEVA438_*glf* kindly provided by Dr. Alberto Sánchez-Pascuala. PCR with Q5 High-Fidelity DNA Polymerase was used for gene amplification and the addition of restriction sites *Pst*I and *Hind*III with primers glf for PstI and glf rev HindIII (**Table S2**). Annealing temperature T_a_ of 57.5 °C was applied and GC enhancer was used in the reaction. The PCR product was purified and digested by *Pst*I and *Hind*III enzymes (New England Biolabs). The plasmid pSEVA2213_*bglC* was isolated and then digested using the same restriction enzymes. Remaining cloning steps were identical to those described above for *lacY* gene. The subcloned *glf* gene was sequenced using the primer Glf seq fw (**Table S2**). Subcloning of *glf* from the newly constructed pSEVA2213_*bglC*_*glf* into pBAMD1-6 was carried out using the same workflow, except for the combination of restriction enzymes which was *Eco*RI and *Hind*III in this case.

#### Random minitransposon-driven insertions in chromosome of P. putida

The insertions of expression cassettes bearing *glf* and *lacY* into the chromosome of *P. putida* EM42 were carried out using mini-Tn5 transposon vectors as described previously (Martínez-García et al., 2014a) with some modifications. The *P. putida* strains were electroporated with plasmids pBAMD1-6_*glf* or pBAMD1-4_*lacY* and left to recover in LB medium for 5 h. The cells were harvested and resuspended in M9 minimal medium. Then 20 mL of selection M9 medium with gentamycin and 4g/L glucose in the case of pBAMD1-6_*glf* or with streptomycin, kanamycin (selection for pSEVA2213_*bglC* plasmid), and 4g/L cellobiose in the case of pBAMD1-4_*lacY* were inoculated with the cells. The culture was left overnight and then the same M9 medium was inoculated from this culture to the final OD 0.1 and the cells were incubated for another 12 h at 30 °C. From this culture, cells were restreaked onto a 20 cm diameter Petri dish with the respective solid M9 agar medium. For further screening, the largest colonies of each strain were picked and their growth was tested in plate cultures. To verify that the entire pBAMD plasmid was not incorporated into the chromosome, the clones were plated also on solid M9 agar with citrate and ampicillin. Only colonies that did not grow on this plate were selected for further experiments. Insertions in *P. putida* chromosome were localized by arbitrary PCR according to (Martínez-García et al., 2014a) (**Tables S2 and S3 in Supplementary material**).

#### Deletion of lldD gene and construction of plasmids pSEVA2213_LDHA_bglC and pSEVA2213_bglC_adhB_pdc

The *lldD* gene that encodes innate L-lactate dehydrogenase (PP4736) was deleted in WT, GCD, and GCDglf_lacY strains using the pCJ012 plasmid and kanamycin/sacB system of selection and counter-selection described in detail previously (Johnson and Beckham, 2015). The *lldD* deletion was verified in selected clones using colony PCR with primers oCJ123 and oCJ124 (**Table S2**). Codon-optimized *LDHA* gene that encodes mutant variant (N109G) of L-lactate dehydrogenase from *Bos taurus* was amplified from pCJ015 plasmid (Johnson and Beckham, 2015) with its synthetic RBS (AGGAGGA) by PCR. The PCR was performed with primers LDHA EcoRI fw and LDHA SacI rv (**Table S2**). Annealing temperature T_a_ was 63 °C. PCR product was purified and digested with *EcoR*I and *Sac*I (New England Biolabs). The *adhB*_*pdc* operon that encodes AdhB alcohol dehydrogenase and Pdc pyruvate decarboxylase from *Zymomonas mobilis* with synthetic ribosome binding sites was synthesized by Genewiz. The delivered plasmid pUC_*adhB*_*pdc* was digested with *Pst*I and *Hind*III. Digested fragments with *LDHA* or *adhB*_*pdc* were ligated into pSEVA2213_*bglC* plasmid cut with the same restriction endonucleases. Chemocompetent cells of *E. coli* CC118 were transformed with the ligation mixtures and cells were plated on LB agar plates with kanamycin and incubated at 37 °C overnight. Transformants were checked for the presence of the plasmid by colony PCR and by restriction analysis of the isolated plasmids. The sequence of cloned *LDHA* gene or *adhB*_*pdc* operon was eventually verified by sequencing using oligos PS1new and LDHA seq rv or adhB seq fw, pdc seq fw, and PS2new, respectively (**Table S2**). Sequences of synthetic operons *LDHA_bglC* and *bglC_adhB_pdc* with respective regulation sequences are provided in **Supplementary material**.

### 2.3 Analytical methods

The optical density in cell cultures was recorded at 600 nm using Spectronic GENESYS 5 (TermoFisher Scientific). Analytes from cultures were collected by withdrawing 0.5 ml of culture medium. The sample was then centrifugated (16,300 g, 10 min). The supernatant was filtered through 4 mm syringe filters 0.45 µm LUT Syringe Filters (Labstore) and immediately frozen at −20 °C. Prior to the HPLC analysis, 50 mM H_2_SO_4_ was added to the samples to stop any hydrolytic activity.

HPLC was used to quantify organic acids and cellobiose. In cases where strains did not produce gluconate and 2-ketogluconate (Δ*gcd* strains), HPLC was also used to measure the concentration of glucose. HPLC analysis was carried out using Agilent 1100 Series HPLC Value System (Agilent Technologies) or Agilent Infinite II 1260 (Agilent Technologies) equipped with a refractive index detector and Hi-Plex H, 7.7 x 300 mm, 8 µm HPLC column (Agilent Technologies). Samples of culture supernatants were diluted twice with milliQ water with 50 mM H_2_SO_4_. The parameters were: mobile phase: 5 mM H_2_SO_4_, injection volume: 20 µL, column temperature: 65 °C, detector temperature: 55 °C, flow of mobile phase: 0.5 mL/min. Glucose, cellobiose, acetate, pyruvate, and lactate standards (Sigma-Aldrich) were used for the preparation of calibration curves.

Glucose in cell cultures was also quantified by a commercial Glucose (GO) Assay Kit (Sigma-Aldrich) according to the manufacturer’s instructions. Product concentration was measured spectrophotometrically at 540 nm using Infinite 200 PRO plate reader (Tecan). Both gluconate and glucono-δ-lactone were quantified using D-Gluconic Acid/D-Glucono-δ-lactone Assay Kit (Megazyme) according to the manufacturer’s instructions.

For the measurement of 2-ketogluconate, an assay described in Lanning and Cohen (1951) was used. First, 150 µL of *o*-phenylenediamine hydrochloride 2.5% (w/v, in water) was added to 150 µL of sample and 100 µL of milliQ water. Then the solution was heated in a thermoblock to 60 °C for 60 min. Samples were quickly spun and 250 µL were pipetted into a 96-well plate. Absorbance was measured at 330 nm using Infinite 200 PRO plate reader (Tecan).

Ethanol in cultures of *P. putida* recombinants was quantified using Trace 1310 Gas Chromatograph (Thermo Fisher Scientific) equipped with ZB-WAX plus column (30 m, 0.25 I.D., 0.25 µm, Phenomenex) and combined with ISQ 7000 Single Quadrupole Mass Spectrometer (Thermo Fisher Scientific). Falcon tubes with *P. putida* cultures were centrifuged (2,500 g, 15 min), the cap was penetrated, and 1.9 mL of supernatant was withdrawn for the GC-MS analysis in 2 mL glass crimp top vials. The parameters of the GC method were as follows: 1 µL sample volume, inlet temperature 170 °C, front inlet flow 1.0 mL/min, split flow 5.0 mL/min, split ratio 5.0, temperature ramp: 40 – 75 °C at the rate of 6 °C/min, 75 – 220 °C at the rate of 30 °C/min, hold time 3 min at 220 °C. ISQ settings were: MS transfer line temp. 250 °C, ion source temp. 300 °C, ionization mode EI, mass range 20 – 200, dwell or scan times 0.2 s. Ethanol absolute (Sigma-Aldrich) was used for the preparation of the calibration curve in M9 medium.

#### 13C-based metabolic flux analysis

A 250-mL-Erlenmeyer flask containing 50 mL M9 medium was inoculated with an overnight culture to an OD_600_ of 0.05. The media contain either 20 mM 50 % U-^13^C glucose, 100 % 1-^13^C glucose (Cambridge Isotope Laboratories), or unlabelled glucose. Cultures were grown agitated to an OD_600_ of 0.5, at which point 5 mL of culture was withdrawn and vacuum filtered (Durapore membrane filter, 0.45 µm). Upon filtration, the filter was extracted with 3 mL acetonitrile/methanol/water (40/40/20 %, v/v), acidified with 0.1 M formic acid at −20°C (Rabinowitz and Kimball, 2007)). Full extraction was facilitated by three freeze/thaw cycles in dry ice. Subsequently, cell debris was removed by centrifugation at 17,000 × *g*, 2 min, 4°C. The supernatant was transferred to a new tube and solvents were removed by evaporation at 30°C for 90 min at reduced pressure (Concentrator Plus, Eppendorf). The samples were then fully dried in a freeze dryer and stored at −80°C. Prior to analysis, the samples were reconstituted in 100 µL H_2_O. Label incorporation into the metabolites was measured according to (McCloskey et al., 2016). Measuring was performed in triplicates and corrected for natural isotope abundance through the data obtained from growth on unlabelled glucose. Data of label incorporation was used to calculat fluxes in INCA (Young, 2014).

### 2.4 Kinetic analyses with β-glucosidase BglC

Two assay methods were applied in this study and we followed the recommendations presented in Assay Guidance Manual (Markossian et al. 2021). One method was based on the collection of samples at different time points throughout the assay. This method was used for screening possible inhibitors of BglC as well as for the determination of basic kinetic properties of the enzyme. The purified enzyme was mixed with the tempered 100 mM sodium phosphate buffer of pH 7.0 and tested potential inhibitors of BglC (glucose, gluconate, ketogluconate, gluconolactone, xylose, arabinose) to reach a final concentration of 2.63 nM of the enzyme (total reaction volume 570 µL). The solution was incubated for 5 minutes in the dry heating block at 37 °C. The reaction was started by adding 30 µL of 100 mM (final concentration 5 mM) substrate 4-nitrophenyl β-D-glucopyranoside (Sigma-Aldrich). At regular five-minute intervals, 120 µL of the mixture was periodically withdrawn and mixed in a 96-well plate with 80 µL of 1M Na_2_CO_3_ to stop the reaction. The concentration of the released 4-nitrophenol was measured at 405 nm and the activity of the enzyme was calculated from the linear part of the product formation curve.

The second method, a continuous measurement in a 96-well plate, was used only for the determination of the modality of inhibition of BglC. Each measurement of activity was repeated at least three times. As in the first method, the enzyme was mixed with sodium phosphate buffer (100 mM) and the measured potential inhibitor to reach the final concentration of 2.63 nM of the enzyme. 96-well plate with this solution was incubated at 37 °C for 5 min. The reaction was started by the addition of tempered substrate 4-nitrophenyl-β-D-glucopyranoside (total reaction volume 250 µL). Product formation was recorded photometrically at 405 nm for 40 min at 37 °C in Infinite 200 PRO reader (Tecan). Enzyme activity was calculated from the linear part of the measured curve. A separate calibration curve was constructed for measurement by this method. Due to its instability, glucono-δ-lactone solution was prepared freshly before every measurement. Non-linear regression models in software Prism (GraphPad 8) were used to determine *K_m_*, *V_max_*, *k_cat_* and K_i_ values.

### 2.5 Data analyses, model construction, and simulations

All analyses were done using either Microsoft Excel, Python 3, or MATLAB 2020a (Mathworks Inc.). The growth rate (μ) was calculated from at least three data points of the growth curve in an early exponential growth phase. Biomass yield on glucose and cellobiose (Y_S/X_) was calculated as a lumped value. Specific uptake rates of glucose and cellobiose or lumped uptake were calculated as the growth rate (μ) / biomass yield (Y_S/X_). If intervals are given for data points in figures with culture time courses, these are standard deviations. In each culture experiment, OD_600_ was measured from four biological replicates and concentrations of substrates and metabolites from two biological replicates. Data are presented as means from all biological replicates ± standard deviation calculated in Microsoft Office Excel. The number of independent experiments conducted for each flask culture is indicated in the figure captions.

#### GECKO model construction

We modified the genome-scale metabolic model (GEM) of the bacterium *P. putida* so that in addition to stoichiometry, it also contains information about the capacity of enzymes. There are many methods for integrating a proteome into GEM (Domenzain et al., 2021; Sánchez et al., 2017; Sánchez and Nielsen, 2015). We used proteomic data collected for *P. putida* KT2440 (Kozaeva et al., 2021). The dataset was used for the verification and parametrization of the model. When measuring a large number of proteins, the standard deviations are usually high (Zhang et al., 2013) and the data were therefore adjusted so that 1.96 SD was added to the mean of the measurements.

The GECKO pipeline version 2.0.2 with herein specified modifications was used for the construction of the model and for the integration of proteomic data. GEM iJN1463 of *Pseudomonas putida* KT2440 (Nogales et al., 2020) was obtained from https://github.com/DD-DeCaF/pputida-gem/. This version of the model has corrected gene-protein-reaction rule for oxaloacetate decarboxylase. Two models were constructed: the FBAwMC model referred here as protein constrained or simply pc model (Beg et al., 2007), and an enzyme constrained model here referred as the ec model. Input parameters are listed in **Table S4 in Supplementary Material**. The first few versions of the pc model could not reach the desired growth rate without having an enzyme saturation factor σ > 0.8. The problem was mainly in low turnover numbers for ATP synthase and enzymes in central carbon metabolism. This stems from the fact that the BRENDA database contains only a limited number of entries for some reactions in which *P. putida* has high flux, but in model, organisms are of minor importance, e.g., reverse flux through fructose bisphosphate aldolase. Some enzyme entries do not contain *k_cat_* high enough to provide experimentally measured fluxes. These were turnover numbers for glucose-6-phosphate isomerase, fructose-bisphosphatase, or KDPG aldolase. Extensive manual curation together with the use of proteomic (Kozaeva et al., 2021) and fluxomic (collected for this study) data were then needed to make the simulations correspond more closely to measured fluxes. The list of all manually curated kinetic data is in **Table S5.**

Previously measured turnover numbers and cofactor specificities were set for all three isoforms of enzyme glucose-6-phosphate dehydrogenase (Volke et al., 2021). Therefore, there are two arm pseudo reactions for this reaction, one with NAD^+^ and the other with NADP^+^ as cofactors. Cofactor specificity was also changed in quinate and shikimate dehydrogenases (Singh et al., 2008). There are now two separate reactions for quinate dehydrogenase QUIDHy_NADPNo1 (PP_3002) and QUIDHy_NADNo1 (PP_2406). The same was done for shikimate dehydrogenases: SHK3Dr_NADPNo1 (PP_0074) and SHK3Dr_NADNo1/No2 (PP_3002 or PP_2406). Amino acid racemase reactions were also modified to account for measured data and substrate preferences (Radkov and Moe, 2013).

Initial simulations with ec model failed to achieve the correct distribution of fluxes into the peripheral and direct phosphorylation glucose pathways. This is due to the high *k_cat_* of the initial reaction of glucose dehydrogenase (Gcd) in the peripheral pathway ∼3,600 s^-1^ (Lynch et al., 1975) while the concentration of this enzyme is also not limiting. However, pyrroloquinoline quinone (PQQ) is an essential cofactor of Gcd and it has previously been shown that the presence or absence of PQQ governs flux through this reaction (An and Moe, 2016). COBRA methods are limited by steady-state assumptions (O’Brien and Palsson, 2015). Thus, it was necessary to introduce a partitioning reaction into the model. All three major reactions that are involved in upper glucose metabolism, glucose ABC transporter GtsABCD (GLCabcppNo1), glucose dehydrogenase (Gcd, GLCDppNo1), and gluconate dehydrogenase (Gad, GAD2ktppNo1), produce a pseudometabolite in the ec model. These three pseudometabolites are then eliminated from the network in one (unbalanced) sink reaction. The stoichiometric coefficients of the pseudometabolites in this sink reaction then represent the distribution ratio into the individual pathways. The ratio was calculated from fluxomic data obtained for *P. putida* KT2440 grown on glucose: 18.1:89.9:4.9 for GLCabcppNo1 : GLCDppNo1 : GAD2ktppNo1.

Some proteins and reactions were removed from the model. Proteins P0A103 and Q88RM4 were excluded from the ATP synthase and cytochrome *c* oxidase aa3 reactions respectively. These are assembly proteins, which are therefore not a direct part of the structure of these proteins. Two lumped pyruvate dehydrogenase reactions were removed as there is already a set of three reactions representing each step. One copy of the reaction for α-ketoglutarate dehydrogenase was removed and the remaining copy was renamed. Reactions for quinate and shikimate dehydrogenases (QUIDHy, SHK3Dr) were removed and new ones were introduced (Singh et al., 2008). Reaction L-tryptophan decarboxylase (LTDCLNo1) was removed from the model because it was shown to be non-functional (Koyanagi et al., 2012). Protein pseudometabolites were removed from all ABC transport reactions. Proteins were not assigned to any other transporter reactions. PP_3722 was removed from the reaction representing cysteine racemase (DLCYSR) (Radkov and Moe, 2013). All manually integrated *k_cat_* values were obtained either from the database BRENDA (Chang et al., 2021), SABIO RK (Wittig et al., 2012) or from primary literature sources. Except for ATP synthase (ATPS4rppNo1), during the parameterization of the model by the function modifyKcats.m, the manually integrated *k_cat_* values cannot be modified. All entries in BRENDA that list *P. putida* KT2440 as a source are included in the set of manually curated data (**Table S5**). Following steps in model construction, parametrization, and verification and modifications of the GECKO pipeline are described in **Supplementary material - Supplementary methods.**

#### Simulations

Simulations with all model variants were performed using the solveLP functions for models in RAVEN format (Wang et al., 2018) or optimizeCbModel for models in COBRA format (Heirendt et al., 2019). If flux values were demanded, the loopless solution variant of the optimizeCbModel function was used. Simulations and model construction were performed using MATLAB 2020a (Mathworks).

## 3. Results and discussion

### 3.1 Deletion of gcd promotes co-utilization of glucose and cellobiose in P. putida EM42

Although *P. putida* EM42 bears a gene encoding periplasmic β-glucosidase (PP_1403), the strain was shown to be unable to grow in minimal medium with cellobiose (Dvořák and de Lorenzo, 2018). This implies that the bacterium has not evolved any carbon catabolite repression (CCR) mechanisms favoring either glucose or cellobiose. However, engineered *P. putida* EM42 pSEVA2213_*bglC* empowered with cytoplasmic β-glucosidase from *Thermobifida fusca* (BglC) on low copy plasmid with constitutive EM7 promoter utilized glucose and cellobiose sequentially (Dvořák and de Lorenzo, 2018). Glucose was consumed first, and only when it was completely depleted, cellobiose was utilized. Numerous studies report inhibition of β-glucosidases by glucose or by intermediates of its metabolism (Deshpande et al., 1978; Kawai et al., 2003; Spiridonov and Wilson, 2001). We argued that the sequential utilization of glucose and cellobiose was caused by such an inhibition rather than by a dedicated CCR mechanism. In *P. putida*, glucose is predominantly (up to 90 % of carbon flux from this substrate) metabolized via the periplasmic oxidative pathway initiated by membrane-associated glucose dehydrogenase Gcd (PP_1444, **Figure 1**) (Nikel et al., 2015). This route forms several potential β-glucosidase inhibitors (Kawai et al., 2003; Spiridonov and Wilson, 2001). Hence, we aimed to investigate the utilization of glucose and cellobiose in the *gcd* deficient strain.

For the purpose of the current study, a previously constructed plasmid pSEVA2213_*bglC* was freshly introduced into *P. putida* EM42 and into available *P. putida* EM42 Δ*gcd* mutant (Dvořák and de Lorenzo, 2018). The resulting strains were designated WTbglC and GCDbglC, respectively. These strains were subsequently cultured in M9 medium with 2 g/L glucose and 2 g/L cellobiose (**Figures 2A and 2B**). In the culture with WTblgC, the concentration profiles of the individual substrates coincided with the previously reported data (Dvořák and de Lorenzo, 2018). The second lag phase that marked the transition from glucose metabolism to cellobiose was evident (**Figure 2A**). The growth rates on glucose and cellobiose were not equivalent: 0.52 ± 0.02 h^-1^ and 0.09 ± 0.00 h^-1^, respectively (**Table 2**). This highlighted the need to engineer efficient co-utilization of the two substrates. Gluconate and ketogluconate were detected in the medium which indicated that the peripheral oxidative glucose pathway was active. These organic acids are secreted by the *P. putida* cells during the growth on glucose and then utilized (**Figure 2A**).

**Figure 2.**
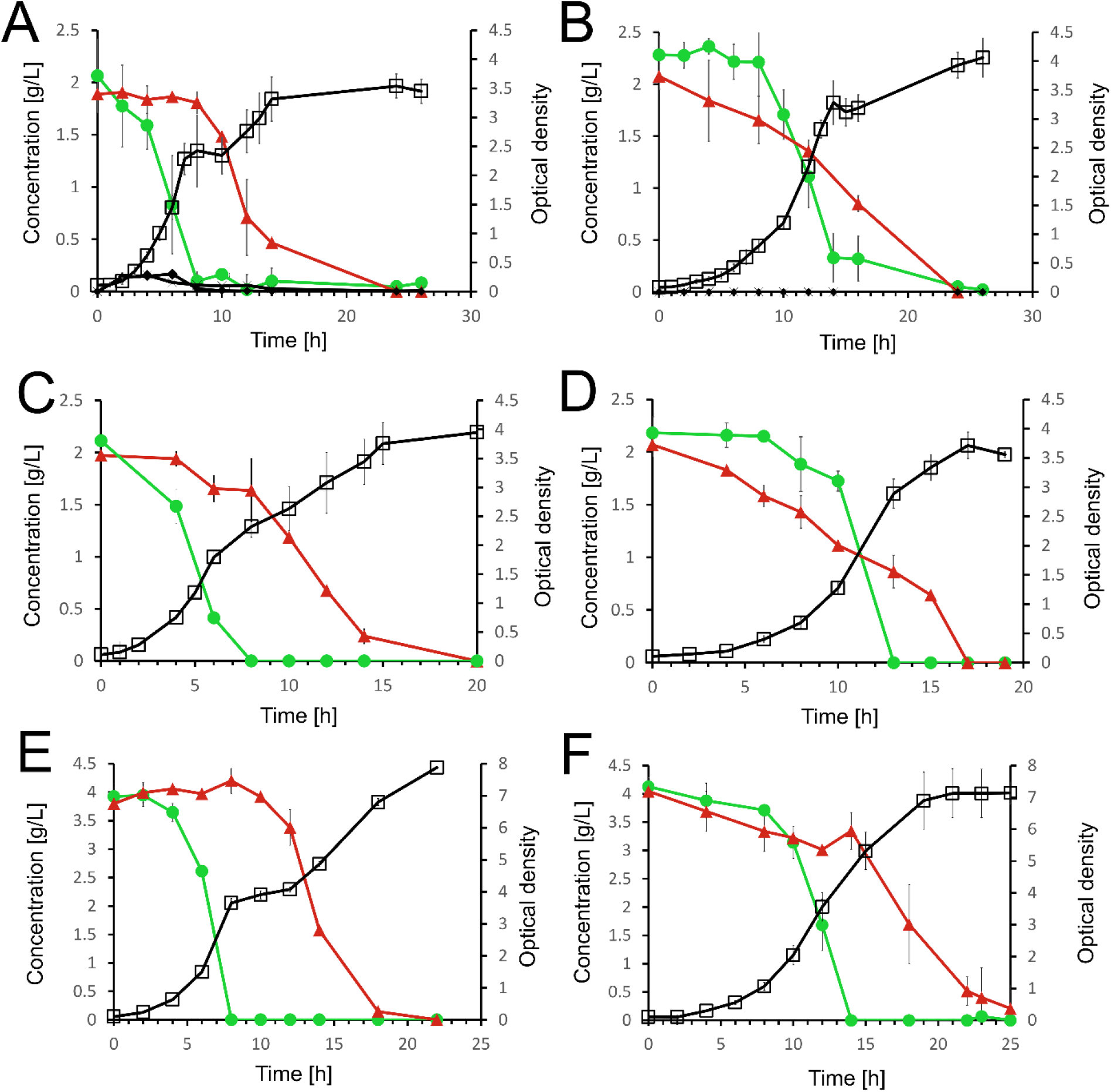
Shake flask cultures of *P. putida* EM42 recombinants in M9 minimal medium with glucose and cellobiose. (A) WTbglC, (B) GCDbglC, (C) hexWTbglC, (D) hexGCDbglC in minimal medium with glucose and cellobiose (2 g/L each). (E) WTbglC, (F) GCDbglC in minimal medium with glucose and cellobiose (4 g/L each). Cell growth, open squares (□); glucose, green line with filled circles; cellobiose, red line with filled triangles; gluconate, filled diamonds (♦); ketogluconate, stars (☓). Data points represent means ± standard deviations of quadruplicate (OD600) or duplicate (concentrations) measurements from two independent experiments.

**Table 2.** Growth parameters of *P. putida* strains in M9 medium supplemented with 2 g/L of glucose and 2 g/L of cellobiose.

| Strain | $\mu \text{ 1}^{\text{st}}$ [$\text{h}^{-1}$] | $\mu \text{ 2}^{\text{nd}}$ [$\text{h}^{-1}$] | Max. D-gluconate conc. [g/L] | Max. 2-ketogluconate conc. [g/L] |
| --- | --- | --- | --- | --- |
| WTbglC | $0.52 \pm 0.02$ | $0.09 \pm 0.00$ | $0.165 \pm 0.025$ | $0.145 \pm 0.035$ |
| GCDbglC | $0.32 \pm 0.01$ | ND | 0 | 0 |
| hexWTbglC | $0.49 \pm 0.02$ | $0.08 \pm 0.00$ | NM | NM |
| hexGCDbglC | $0.30 \pm 0.01$ | ND | NM | NM |

$\mu \text{ 1}^{\text{st}}$ is the growth rate during the first growth phase on glucose; $\mu \text{ 2}^{\text{nd}}$ is the growth rate during the second growth phase on cellobiose; ND, not detected; NM, not measured

On contrary, the growth curve of the GCDbglC strain lacked any apparent second lag phase (**Figure 2B**). However, the growth rate dropped to 0.32 ± 0.01 h^-1^. A similar reduction in growth rate was reported previously for *P. putida* Δ*gcd* mutant cultured on glucose (del Castillo et al., 2007; Dvořák and de Lorenzo, 2018; Elmore et al., 2020; Poblete-Castro et al., 2013). The growth defect can be attributed to the closure of one of the two major glucose uptake routes (**Figure 1**) and the fact that Gcd activity is directly coupled to electron transfer and generation of proton motive force (van Schie et al., 1985). Interestingly, substrate concentration curves did not indicate co-utilization of both substrates during the initial phase of growth. During the first eight hours, cellobiose was utilized exclusively and the glucose concentration remained constant. We attributed this phenomenon to BglC activity in the growth medium. The BglC activity in the medium increased with the time of the culture and it peaked in the transition from logarithmic to stationary phase. For WTblgC the maximum measured BglC activity was 18.3 ± 8.2 U/L. In the case of GCDbglC, a slightly higher activity of 23 ± 6.3 U/L was detected. These values were higher than the trace extracellular BglC activity recorded in the exponential growth phase in our previous work (2.76 ± 0.24 U/L) (Dvořák and de Lorenzo, 2018). Such a result acknowledges the former assumption that a small portion of well-overexpressed enzyme is released out of *P. putida* cells either due to the lysis of dead bacteria or by virtue of a secretion mechanism that remains to be deciphered. No Sec or TAT signal sequence was identified in BglC and none of the non-specific secretion routes has been described for this enzyme either. In any case, the detected BglC activity in the culture media may, to some extent, skew all substrate consumption curves. Cellobiose utilization can be overestimated and glucose utilization underestimated.

### 3.2 Enzyme BglC is inhibited by metabolites in the peripheral glucose pathway

Gluconolactone, gluconate, 2-ketogluconate, and 2-ketogluconate-6-phosphate were not produced in the GCDbglC strain (**Figure 1**). We tested whether one or multiple of the latter three metabolites (2-ketogluconate-6-phosphate was not commercially available) could inhibit BglC activity. In addition, we tested inhibition by other lignocellulosic sugars xylose and arabinose, and also by glucose to check for product inhibition. The enzyme BglC with N-terminal His-tag was overproduced in cells *P. putida* pSEVA238_*bglC* and purified to homogeneity by affinity chromatography (**Supplementary methods, Figure S1**). Determined basic kinetic parameters (*K*_m_ = 0.56 ± 0.01 mM, *k_cat_* = 48.68 s^-1^, and *V_max_* = 53.70 ± 0.11 U/mg, **Figure S2**) were in good agreement with the values measured by other authors for BglC and synthetic *p*-nitrophenyl-β-D-glucopyranoside or cellobiose used as substrates (Pei et al., 2011; Spiridonov and Wilson, 2001).

In the study of Spiridonov and colleagues (2001), xylose and gluconolactone were shown to be inhibitors of BglC. In our work, xylose had no inhibitory effect on BglC while D-gluconate and D-glucono-δ-lactone were confirmed as inhibitors (**Figure S3**) These two chemicals were used for the determination of the modality of inhibition and the measurement of dissociation constants (**see Methods section**). Results obtained for gluconate could be described by neither competitive, non-competitive nor mixed-type inhibition (see **Supplementary results and discussion** for more information). The results obtained for gluconolactone were best described by competitive inhibition (**Figure S4**). The estimated *K_i_* was 6.63 ± 1.3 µM and in good agreement with similar studies (Deshpande et al., 1978; Kawai et al., 2003). Inhibition of hydrolases by their respective lactones is very common. Yet, we cannot state with certainty that the diauxic growth of WTbglC on the glucose-cellobiose mix was caused only by gluconolactone. It is not clear whether gluconolactone is transported from periplasm to cytoplasm where it can inhibit BglC. In addition, there is another gluconolactone produced in *P. putida* EM42 - 6-phosphoglucono-δ-lactone. It is formed in the cytoplasm by glucose-6-phosphate dehydrogenase (Zwf) and converted to 6-phosphogluconate by 6-phosphogluconolactonase (Pgl) (**Figure 1**). The compound was not commercially available and direct measurement of inhibition was thus not feasible. Phosphogluconolactone is formed in all *P. putida* strains prepared in this study, but the highest production was expected in GCDbglC where all carbon is processed through Zwf and Gnl reactions. This strain nonetheless displayed substrate co-utilization, and D-glucono-δ-lactone thus remains a prime candidate for the BglC inhibition in engineered *P. putida*. Next, we redirected our attention towards increasing the growth rate of the GCDbglC mutant.

### 3.3 Deletion of hexR has no significant effect on co-utilization of glucose and cellobiose and growth rate of *Δ*gcd *mutant*

The glucose-cellobiose co-utilization was a promising feature of the GCDbglC strain. But the growth defect narrowed this benefit. The deletion of *hexR* (PP_1021) was reported to accelerate the growth on glucose of *P. putida* KT2440 Δ*gcd* mutant (Bentley et al., 2020). HexR repressor controls the expression of enzymes in the direct phosphorylation pathway (Glk, Zwf, Edd, Eda, and GapA, **Figure 1**) which was the only remaining route for glucose utilization in GCDbglC (Bentley et al., 2020; Udaondo et al., 2018). We, therefore, aimed to probe this strategy in our study. First, the gene *hexR* was deleted in strain WTbglC resulting in strain hexWTbglC. We hypothesized that the deletion could at least partially redirect cellular flux from the peripheral oxidative pathway towards the direct phosphorylation route and thus the upper glycolytic fluxes in hexWTbglC could resemble those in GCDbglC. We hoped to obtain a strain with the ability to co-utilize glucose and cellobiose but at a growth rate approaching WTbglC.

Strain hexWTbglC was cultured on the mixture of glucose and cellobiose (2 g/L each) (**Figure 2C**). Unlike WTblgC, the growth curve measured for the hexWTbglC strain did not have a sharp transition between the two growth phases. However, the preference for glucose was evident. In the first growth phase, the growth rate of this strain was comparable to WTbglC: 0.49 ± 0.02 h^-1^ (**Table 2**). In the second growth phase (cellobiose utilization) the growth rate dropped to 0.08 ± 0.00 h^-1^, again, this value was very close to that observed for WTbglC. It was concluded that *hexR* deletion, leading to the de-repression of genes of enzymes in the direct phosphorylation pathway (del Castillo et al., 2008; Udaondo et al., 2018), was not enough to establish the co-utilization of glucose and cellobiose.

Next, strain hexGCDbglC was constructed to increase the capacity of the direct phosphorylation pathway on the Δ*gcd* mutant background. The strain cultured on the mixture of glucose and cellobiose (**Figure 2D**) showed a growth profile and growth rate very similar to those of GCDbglC (**Figure 2B**, **Table 2**). Thus, our data did not confirm the results obtained by Bentley and co-workers (2020). It is nonetheless necessary to consider the differences between their *P. putida* KT2440 strain and the EM42 recombinant used in this study. The parental engineered strain of Bentley and colleagues possessed other modifications besides the Δ*gcd* deletion and had a growth rate of ∼0.1 h^-1^ (Bentley et al., 2020). This growth rate is much lower than the one observed herein (0.32 ± 0.01 h^-1^) and the strains are therefore not directly comparable. We thus demonstrated that the deletion of the repressor gene *hexR* has no significant effect on glucose-cellobiose co-utilization. We hypothesized that a bottleneck in the metabolism of glucose and cellobiose in Δ*gcd* mutants may lie in the transport of these substrates, which is not influenced by HexR (Udaondo et al., 2018). It was observed in our previous study that the glucose ABC transporter GtsABCD (**Figure 1**) is probably partially responsible for the uptake of cellobiose as the mutant strain *P. putida* Δ*gts* showed reduced growth on this disaccharide (Dvořák and de Lorenzo, 2018). In the Δ*gcd* mutant, the ABC transporter is the only remaining entry point also for glucose. This fact made us speculate that cellobiose and glucose transport may become limiting upon *gcd* deletion.

### 3.4 Expression of additional glucose transporter improves the growth of Δgcd mutants

Due to the deletion of *gcd*, all carbon flux from glucose and probably part from cellobiose flows through the ATP-dependent transporter GtsABCD. The energy status of the cell might be disturbed because of additional ATP expense. This situation can be further exacerbated by the reduced electron supply from periplasmic glucose oxidation into the respiration chain (van Schie et al., 1985). Hence, glucose facilitator *glf* from the bacterium *Zymomonas mobilis* was chosen as an additional transport route with no ATP demand. Glf is a high-capacity/low-affinity transporter (Parker et al., 1995) which could well complement the remaining high-affinity glucose transporter GtsABCD in *P. putida* Δ*gcd* strain (del Castillo et al., 2007). In previous studies that took advantage of the *glf* gene in heterologous bacterial hosts, satisfactory result was obtained only after optimization of its expression (Kyselova et al., 2018; Tang et al., 2013). We first subcloned *glf* from available plasmid pSEVA438_*glf* to low-copy pSEVA2213_*bglC*_*glf* with a constitutive EM7 promoter and inserted the plasmid construct into the GCD strain. The resulting recombinant *GCDbglC_glf* with *glf* expressed from plasmid did not show significantly faster growth when compared with the GCDbglC strain (**Figure S5**). This could be caused by the suboptimal expression of *glf* from the plasmid. Hence, we aimed to optimize the expression of *gcd* by its random insertion into *P. putida* chromosome using a suicide pBAMD1-6 plasmid with mini-Tn5 transposon sequences (**Table 1**) (Martínez-García et al., 2014a). In this way, a library of mutants with different expression levels of *glf*, which largely depend on the context of chromosomal insertion, was created.

Four parental strains (WT, GCD, hexWT, and hexGCD) were electroporated with pBAMD1-6_*glf*. Mutants were selected from the constructed libraries based on their growth on glucose. The 23 fastest-growing colonies for each template strain were selected and tested in a 96-well plate growth assay (**Figure S6**). The five fastest-growing strains were selected and re-tested (**Figure S7**). The fastest-growing mutants obtained from the screening were named WTglf, GCDglf, hexWTglf, and hexGCDglf. The mutant strains WTglf and hexWTglf did not grow faster than the parent strains, indicating that the enhancement of the direct phosphorylation pathway on this background is unlikely to provide a growth advantage. In contrast, new mutants with Glf transporter and *gcd* deletion showed faster growth than their parental strains (**Figure S7**). The sites of the random insertions of *glf* gene are listed in **Table S3**.

### 3.5 Effects of expression of additional cellobiose transporter

The next step was to find a suitable transporter for cellobiose. We looked for a simple bacterial permease for this purpose. Searches in the Transporter Classification Database (Saier et al., 2021) and in Carbohydrate Active enZYmes (CAZy database) (Levasseur et al., 2013) did not show any suitable candidates. Sekar et al., (2012) demonstrated that the lactose transporter LacY from *E. coli* is also able to transport cellobiose (Sekar et al., 2012). The gene *lacY* was cloned from the chromosome of *E. coli* BL21(DE3) into pSEVA2213_*bglC* downstream *bglC* and then subcloned into the mini-Tn5 delivery plasmid pBAMD1-4 (**Table 1**, Martínez-García 2014). Six parental strains (WT_bglC, WTbglC_glf, hexWTbglC, hexWTbglC_glf, GCDbglC_glf, and hexGCDbglC_glf) were electroporated with the latter plasmid. Growth in 96-well plate on cellobiose did not show any improvement over parental strains (data not shown). Hence, we cultured five fast-growing colonies from cellobiose agar plates for each of the template strains in a 48-well plate, and here the effect of the additional cellobiose transporter was more evident (**Figure S8**). Interestingly, the strains that benefited most from the insertion of *lacY* were those without *gcd* deletion – WTbglC_glf_lacY, hexWTbglC_lacY, hexWTbglC_glf_lacY. The only exception was WTbglC_lacY which did not grow better on cellobiose than its parent WTbglC. The insertion of mini-Tn5 transposons into the genome is random, so it is possible that we coincidentally missed the WTbglC_lacY strain with improved performance during the screening. Strains with *gcd* deletions (GCDbglC_glf_lacY and hexGCDbglC_glf_lacY) showed only a minor improvement in growth when compared with controls (**Figure S8**) but they were used for further work due to their potential to co-utilize cellobiose with glucose.

The improvement of the growth of strains WTbglC_glf_lacY, hexWTbglC_lacY, and hexWTbglC_glf_lacY on cellobiose (**Figure S8**) begged the question of whether the preference for glucose in WT strain was not caused solely by higher affinity and transportation capacity of the common transporter GtsABCD for glucose. In such a case, additional cellobiose transporter should enable monoauxic growth on the sugar mix. However, diauxic growth was observed for three WT derivatives cultured on the mixture of glucose and cellobiose (**Figure S9**) which confirmed that the deletion of *gcd* was necessary to establish co-utilization.

### 3.6 The high growth rate of Δgcd mutants with Glf and LacY transporters is associated with the overproduction of pyruvate and acetate

Next, we focused on the characterization of new mutants GCDbglC_glf_lacY and hexGCDbglC_glf_lacY. For all subsequent co-utilization experiments glucose and cellobiose concentrations were fixed at 4 g/L for each sugar to make diauxia and monoauxia more distinguishable. Other culture conditions were adapted to the higher substrate concentrations (see **Methods**). Growth parameters of template strains WTbglC and GCDbglC remained almost unchanged in modified conditions (**Table 3**, **Figures 2E and 2F**). New culture time courses confirmed that WTbglC grows in two separated growth phases while the GCDblgC strain has only one growth phase and co-utilizes glucose and cellobiose at the beginning of the culture. Subsequently, however, only glucose was utilized and the concentration of cellobiose remained constant. Towards the end of cultivation, only cellobiose was utilized and the growth rate decreased, which suggested that the utilization of this substrate in *P. putida* GCDblgC recombinant was not as effective as the utilization of glucose.

**Table 3.** Growth parameters of engineered *P. putida* strains determined in shake flask cultures. Strains were cultured in M9 medium with 4 g/L glucose and 4 g/L cellobiose.

| Strain | $\mu$ 1<sup>st</sup> [h<sup>-1</sup>] | $\mu$ 2<sup>nd</sup> [h<sup>-1</sup>] | Y<sub>S/X</sub> [g<sub>CDW</sub>/g<sub>subst</sub>] |
| --- | --- | --- | --- |
| WTbglC | 0.52 ± 0.02 | 0.08 ± 0.00 | 0.31 ± 0.01 |
| GCDbglC | 0.31 ± 0.02 | ND | 0.28 ± 0.02 |
| WTbglC_glf_lacY | 0.49 ± 0.01 | 0.04 ± 0.00 | NM |
| hexWTbglC_glf_lacY | 0.47 ± 0.02 | 0.07 ± 0.00 | NM |
| GCDbglC_glf_lacY | 0.43 ± 0.01 | ND | 0.54 ± 0.02 |
| hexGCDbglC_glf_lacY | 0.36 ± 0.01 | ND | 0.30 ± 0.01 |

$\mu$ 1<sup>st</sup> is the growth rate during the first growth phase on glucose; $\mu$ 2<sup>nd</sup> is the growth rate during the second growth phase on cellobiose
Y<sub>S/X</sub> is the biomass yield on both glucose and cellobiose calculated at the time when a culture reached maximum OD<sub>600</sub>
ND, not detected; NM, not measured

The newly engineered strains GCDbglC_glf_lacY and hexGCDbglC_glf_lacY were then cultured in the same conditions (**Figures 3A and 3B**). The optical density during the culture of the strain GCDbglC_glf_lacY reached a value of 7.03 ± 0.13 which is comparable to GCDbglC (OD_600_ = 7.38 ± 0.29) and WTblgC (OD_600_ = 8.0 ± 0.10). The growth rate increased by 39 % when compared with GCDbglC (**Table 3**). But this high growth rate was maintained for only 6 hours and then the culture was not growing exponentially anymore. The slowdown was likely caused by the accumulation of pyruvate and acetate in cells and in the medium (**Figure 3A**). Due to the overproduction and secretion of acids, the pH in the culture of GCDbglC_glf_lacY dropped to ∼5 after the first 12 hours, and then the growth was almost halted because *P. putida* is an organism with a relatively narrow pH optimum (Reva et al., 2006). The growth arrest resulted in the presence of ∼2 g/L of residual glucose in the medium. Under the same conditions, the pH of the M9 medium in the culture of strain WTbglC declined to 6.61, and in the culture of GCDbglC to 6.71 from the original pH of ∼6.9. It is worth noting that the strain GCDbglC_glf_lacY showed greatly increased biomass yield (0.54 ± 0.02 g_biomass_/g_substrate_) compared to WTbglC (0.31 ± 0.01 g_biomass_/g_substrate_) and GCDbglC (0.28 ± 0.02 g_biomass_/g_substrate_). The final yield of pyruvate in the culture of the mutant strain was 0.21 ± 0.01 g/g_substrate_ and the final yield of acetate was 0.09 ± 0.00 g/g_substrate_. This data indicates that the strain GCDbglC_glf_lacY forms less CO_2_ as a final product of aerobic metabolism and that it converts the substrate to biomass more efficiently than WT.

**Figure 3.**
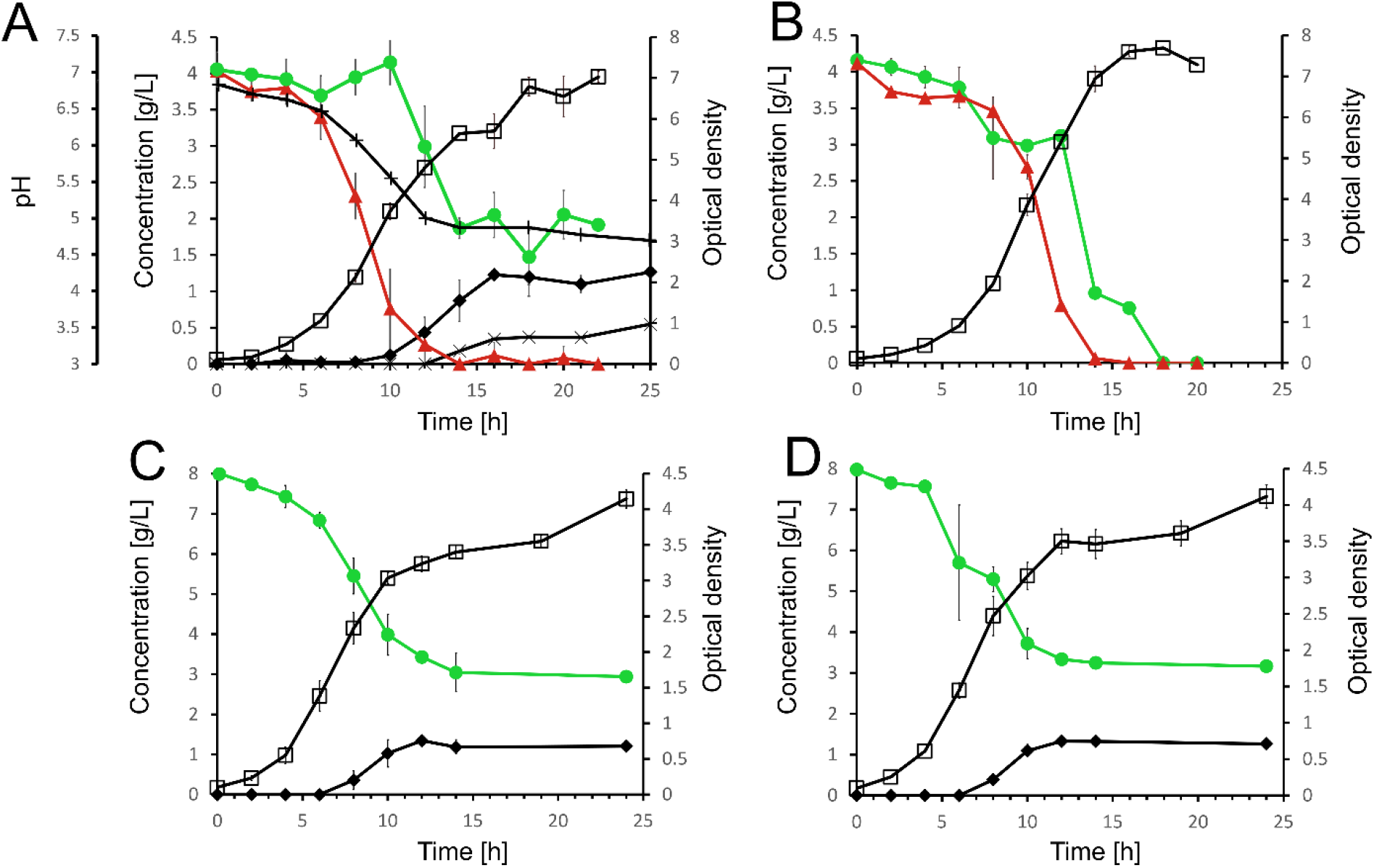
Shake flask cultures of *P. putida* mutants in M9 minimal medium with the mixture of glucose and cellobiose or with glucose only. (**A**) and (**B**) GCDbglC_glf_lacY and hexGCDbglC_glf_lacY, respectively, on glucose and cellobiose (4 g/L each) (**C**) and (**D**) GCDglf and GCDbglC_glf_lacY, respectively, on 8 g/L glucose. Cell growth, open squares (□); glucose, green line with filled circles; cellobiose, red line with filled triangles; pyruvate, filled diamonds (♦); acetate, stars (☓); pH, crosses (+). Data points represent means ± standard deviations of quadruplicate (OD600) or duplicate (concentrations) measurements from two independent experiments in case of (A) or from a single experiment in case of (B), (C), and (D).

Interestingly, the substrate utilization curves suggested a reverse diauxia – cellobiose concentration declined rapidly while glucose was initially utilized slowly and its concentration even increased between the sixth and tenth hour of the culture (**Figure 3A**). We revealed that the apparent preference for cellobiose could be a result of a high cellobiohydrolytic activity in the medium. The β-glucosidase activity reached 1.54 ± 0.54 g_cellobiose_/L/h at the eighth hour of the culture (**Figure S10**) which was sufficient to cause a rapid decrease in cellobiose concentration and accumulation of glucose. It could be caused by increased cell death rate, lysis, and following BglC release due to the acidification of the intracellular environment. It is, therefore, probable that both substrates were co-utilized as in GCDbglC cultures, but the co-utilization was masked in the reaction courses due to the intensified release of BglC.

In the cultivation with the hexGCDbglC_glf_lacY strain (**Figure 3B**), a maximum optical density of 7.69 ± 0.13 was reached. The growth rate (0.36 ± 0.01) was ∼16% lower than that of GCDbglC_glf_lacY, but still ∼16% higher than of GCDbglC and hexGCDbglC. Surprisingly, no pyruvate and only traces of acetate were detected. The cause of the lower growth rate and absence of acids in the cultures of the mutant with *hexR* deletion was ambiguous. The only difference, when compared with GCDbglC_glf_lacY, was the expected upregulation of genes in upper glucose metabolism (del Castillo et al., 2008; Udaondo et al., 2018). After investigating the insertion sites of *glf* and *lacY* expression cassettes in this strain, we argued that the strain grew slower because *glf* was inserted into gene PP_1318 which encodes cytochrome *b*. Cytochrome *b* is an essential part of the most abundant electron transfer complex in bacteria. However, when we cultured another promising hexGCDbglC_glf_lacY clone obtained from the screening (hexGCDglf colony 2 with *glf* insertion in locus PP_0909 encoding a hypothetical protein, **Table S3**) and compared it to the first candidate, the two strains showed very similar growth profiles and no accumulation of acids (**Figure S11**). An alternative explanation for the different behavior of *hexR* mutant is that due to the randomness of Tn5 minitransposon-driven insertions, the expression of transporters in strains GCDbglC_glf_lacY and hexGCDbglC_glf_lacY is unlike. In any case, glucose and cellobiose (4 g/L each) were fully utilized by hexGCDbglC_glf_lacY mutant and maximum OD_600_ in its cultures was reached within 18 h interval. This was achieved neither with WTbglC nor with other GCDbglC strains. Thus, this mutant can be an attractive alternative to GCDbglC_glf_lacY in applications where the overproduction of pyruvate is not a desirable feature.

### 3.6 Pyruvate overproduced in recombinant P. putida can be converted into desirable chemicals

In the next step, we explored whether the excess carbon in the metabolism of the GCDbglC_glf_lacY strain can be redirected toward higher-value products. Pyruvate is a key metabolic precursor for the biosynthesis of numerous substances of industrial and pharmaceutical interest - alcohols, terpenoids, phenazines, propionate, or lactate (Johnson and Beckham, 2015; Maleki and Eiteman, 2017). Its overproduction in aerobic conditions without the need for blocking the pyruvate utilization pathways is an attractive addition to *P. putida*’s biosynthesis repertoire (Maleki and Eiteman, 2017). We selected two chemicals whose biosynthesis from pyruvate is most straightforward – ethanol and lactate. Production of both these compounds was already reported in engineered *P. putida* KT2440 with exogenous biosynthetic routes. However, ethanol biosynthesis from glucose was described only in anoxic conditions (Nikel and de Lorenzo, 2014, p. 201), and aerobic lactate production was achieved from benzoate and *p*-coumarate but not from glucose or cellooligosaccharides (Johnson and Beckham, 2015). We equipped the pyruvate overproducing mutant GCDbglC_glf_lacY and two control strains WTbglC and GCDbglC with ethanol biosynthesis route from *Z. mobilis* or with mutant variant (N109G) of bovine L-lactate dehydrogenase LDHA (**Figure 1**). This evolutionary distant enzyme was previously adopted by Johnson and Beckham (2015) for aerobic conversion of accumulated pyruvate to L-lactate in *P. putida* cultures grown on lignin-derived aromatics (Johnson and Beckham, 2015).

Codon-optimized genes encoding the ethanol pathway (*adhB* and *pdc*) or L-lactate dehydrogenase (*LDHA*) were subcloned into pSEVA2213_*bglC* construct and distributed into all tested strains. To prevent the reverse conversion of lactate to pyruvate, the gene of innate lactate dehydrogenase LldD (PP4736) was deleted in GCDbglC_glf_lacY, WTbglC, and GCDbglC strains before their transformation with pSEVA2213_*LDHA*_*bglC* plasmid (Escapa et al., 2012; Johnson and Beckham, 2015). The resulting recombinants were grown for 24 h in M9 medium with glucose and cellobiose (4 g/L each) in shake flasks (lactate production) or in falcon tubes (ethanol production, see **Materials and methods** for details). Product concentrations, as well as residual quantities of glucose and cellobiose, were determined at the end of the cultures and used for the calculation of product titer (g/L) and product yield (g_product_/gs_ubstrate_). Results are summarized in **Figure 4**. GCDbglC_glf_lacY mutant surpassed WTbglC and template strain GCDbglC in the production of both ethanol and L-lactate. GCDbglC_glf_lacY produced 139 % more ethanol (0.52 ± 0.14 g/L) than WTbglC (0.22 ± 0.13 g/L). Ethanol yield from sugar substrates was increased by 148 % in the former strain (0.13 ± 0.03 g/g *vs*. 0.05 ± 0.03 g/g for WTbglC). Only traces of ethanol were detected in GCDbglC cultures and no alcohol was measured in GCDbglC_glf_lacY control without *adhB* and *pdc* genes. Acetate (up to 0.69 ± 0.17 g/L in GCDbglC_glf_lacY cultures) and lactate (up to 0.31 ± 0.05 g/L for GCDbglC_glf_lacY control without ethanol pathway) but no pyruvate were detected in cultures of all compared strains. This indicates that pyruvate was indeed streamed to ethanol but part of the carbon was secreted in the form of organic acids whose production contributes to NAD^+^ regeneration (lactate, **Figure 1**) during cultivation in falcon tubes with suboptimal aeration.

**Figure 4.**
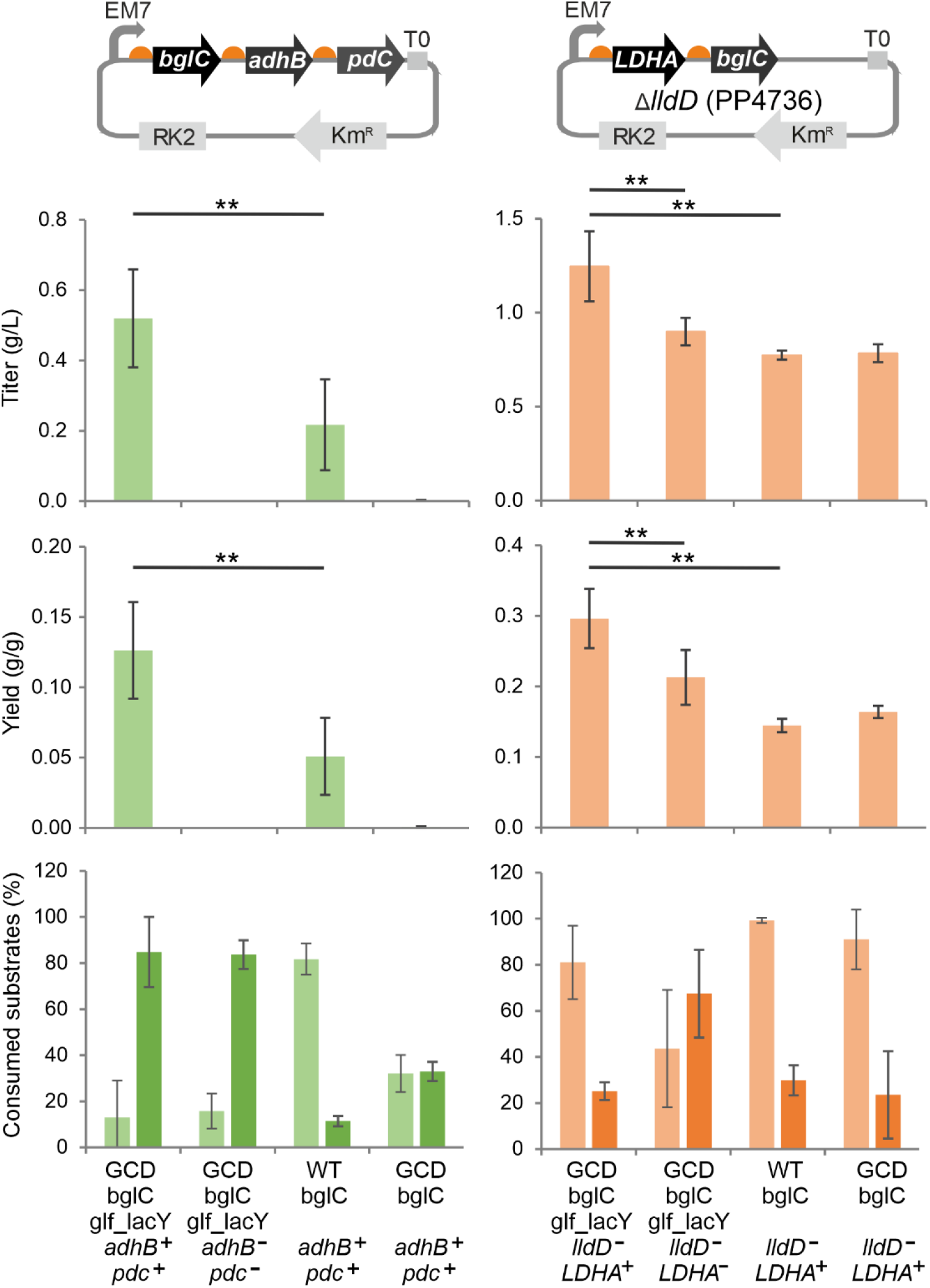
Comparison product titer (g/L), yield (gproduct/gsubstrate), and substrate consumption (%) in cultures of engineered *P. putida* strains endowed with exogenous ethanol (graphs with green columns) and L-lactate (graphs with orange columns) biosynthetic pathways from *Zymomonas mobilis (adhb* and *pdc* genes) and *Bos taurus* (*LDHA* gene). The shown values were calculated from substrate and product concentrations determined at the end of 24 h cultures. The bottom graphs show % of glucose (pale green or orange) and cellobiose (dark green or orange) consumed at the end of cultures. Pictures above graphs depict pSEVA2213 plasmids with subcloned *bglC* gene of β-glucosidase, biosynthetic genes from *Z. mobilis* (*adhB* gene of alcohol dehydrogenase and pyruvate decarboxylase gene *pdc*) or *B. taurus* (*LDHA* gene of L-lactate dehydrogenase), and other relevant sequences (constitutive EM7 promoter, ribosome-binding sites as orange hemispheres, T0 transcriptional terminator, RK2 origin of replication, kanamycin resistance gene). Note that the elements in these schemes are not to scale. All strains tested for the production of L-lactate were deprived of the *lldD* gene that encodes innate lactate dehydrogenase. Bars represent mean value ± standard deviation calculated from at least six cultures conducted in two independent experiments. Asterisks denote the significance of the difference between two means at P < 0.01 as calculated using the two-tailed Student’s t test in Microsoft Office Excel.

Pyruvate was absent also in cultures of L-lactate-producing strains. GCDbglC_glf_lacY with deleted *lldD* and implanted *LDHA* accumulated 1.25 ± 0.19 g/L of L-lactate, ∼60 % more than WTbglC (0.77 ± 0.02 g/L) and GCDbglC (0.78 ± 0.05 g/L) derivatives (**Figure 4**). Lactate yield from glucose and cellobiose was doubled in GCDbglC_glf_lacY (0.30 ± 0.04 g/g) when compared with both controls. Interestingly, a considerable quantity of lactate (0.90 ± 0.07 g/L) was discerned also in cultures of GCDbglC_glf_lacY control without *LDHA* gene. This signs that *lldD* deletion on its own promotes the generation of lactate from pyruvate in aerobic *P. putida* cultures.

It should be noted here that after 24 h interval, strain GCDbglC_glf_lacY consumed a lower total amount of glucose and cellobiose (∼47 % in ethanol-producing cultures and ∼55 % in lactate-producing cultures) than WTbglC control (∼49 % in ethanol-producing cultures and ∼67 % in lactate-producing cultures) in all compared cultures (**Figure 4**). Hence, its enhanced ethanol and lactate production capacity cannot be attributed to faster consumption of substrates but rather to its ability to overproduce pyruvate. Reduced consumption of cellobiose and increased utilization of glucose when compared with ethanol-producing recombinants were observed in the cultures of GCD strains with *LDHA* gene (**Figure 4**). We argue that the increased distance between *bglC* and EM7 promoter after *LDHA* insertion upstream *bglC* in the pSEVA2213 construct could affect the expression of the β-glucosidase gene (Lim et al., 2011). This effect can be prevented, *e.g.*, by implanting *LDHA* into the chromosome of *P. putida* host (Johnson and Beckham, 2015).

### 3.7 Additional cell cultures and computational analyses of P. putida metabolism elucidate aerobic overproduction of pyruvate and acetate

We demonstrated that GCDbglC_glf_lacY can stream pyruvate into desirable chemicals more efficiently than WT and GCD strains but it was still not clear why GCDbglC_glf_lacY overproduced acids. Under given conditions (minimal medium with glucose, sufficient aeration), the accumulation of acids other than gluconate and ketogluconate in *P. putida* KT2440 culture is unprecedented. Pyruvate and acetate were detected in the late stationary phase of aerobic cultures of *P. aeruginosa* P4 and *P. fluorescens* 13525 grown in M9 minimal medium with excess (at least 12 g/L) of glucose (Buch et al., 2008). But in *P. putida* KT2440, acetate secretion was reported only at limiting oxygen concentrations (Kampers et al., 2021; Nikel and de Lorenzo, 2014; Sohn et al., 2010). The leakage of pyruvate and acetate was detected in aerobic LB medium cultures with KT2440 mutant that lacked global regulator Crc (Molina et al., 2019). Pyruvate was also overproduced from lignin-born aromatic molecules in KT2440 strain endowed with the exogenous meta-cleavage pathway from *P. putida* mt-2 (Johnson and Beckham, 2015). Among these different scenarios, it was difficult to find a single link that would point out what exactly led to the overproduction and secretion of pyruvate and acetate in our recombinant. We argued that this phenomenon could be caused by the fact that there were multiple overexpressed proteins in the strain. The total protein concentration in the cell is almost constant (Sánchez et al., 2017), and the overproduction of particular proteins can limit the synthesis of the others. In such a case, enzymes can be rapidly saturated, and the excess metabolites secreted into the medium (Bolognesi and Lehner, 2018). The expression limit can be variable and depends a lot on the organism and the type of protein. Our strains expressed several exogenous genes – *bglC*, *glf*, *lacY*, and antibiotic resistance genes from pSEVA2213 plasmid and synthetic expression cassettes in the chromosome. Therefore, we decided to carry out a simple experiment: the cultivation of strains GCDglf and GCDbglC_glf_lacY on glucose (8 g/L, **Figures 3C and 3D**). Though the former strain overexpressed four genes less, the two recombinants showed almost identical growth curves. The growth rate was 0.45 ± 0.00 h^-1^ for GCDglf and 0.44 ± 0.00 h^-1^ for GCDbglC_glf_lacY. Both strains also accumulated ∼1.4 g/L of pyruvate after 12 h and pH in the medium dropped to ∼4.5. Hence, it was concluded that the accumulation of acids was not caused by the overexpression of multiple exogenous genes.

The observed acid overproduction could also possibly stem from *glf* insertion into gene PP_5410 which encodes the DeoR family transcriptional regulator. To the best of our knowledge, this putative regulator was not yet investigated. We wanted to make sure that the site of the insertion played no role in the acid secretion. Hence, we cultured another GCDglf strain with a different insertion site in gene PP_1204 that encodes a hypothetical protein (GCDglf colony 2, **Table S3**) on 8 g/L glucose. Two specific features of GCDbglC_glf_lacY – fast initial growth and secretion of organic acids - were both confirmed in cultures of GCDglf colony 2 (**Figure S12**). During the initial five hours of the culture, the growth rate reached 0.53 ± 0.04 h^-1^, which is an unprecedented value for *P. putida* Δ*gcd* mutant. Then, the growth decelerated and due to the limited consumption of glucose, the maximum OD_600_ of only ∼1.5 was reached after 24 h. Pyruvate (1.73 g/L ± 0.17 g/L) and to a minor extent also acetate (0.39 ± 0.01 g/L) were both accumulated in the culture medium after this time interval. Consequently, it could be concluded that the overproduction of acids on glucose was a common feature of GCDglf hits from screening regardless of *glf* insertion site.

Last but not least, we cultured WTbglC on 8 g/L of glucose in conditions identical to the cultivations with GCDglf recombinants to probe pyruvate and acetate accumulation by the parent *P. putida* strain (**Figure S13**). Only traces of pyruvate and acetate were detected in WTbglC cultures. Thus, the overproduction of acids under given conditions was neither a result of a random gene interruption nor the pristine property of template *P. putida* strain, and other factors were at play.

Our attention turned to the two implanted transporters and especially to the high-capacity/low-affinity Glf from *Z. mobilis*, a bacterium that, unlike *P. putida*, is adapted to live in conditions that are very rich in carbohydrates (Parker et al., 1995). Internal fluxes in *Z. mobilis* are very high and the glucose uptake rate can reach more than 60 mmol/g_cdw_/h (Fuhrer et al., 2005). Our hypothesis was that the integration of *glf* in *P. putida* caused unregulated uptake of the substrate and led to the saturation of catabolic enzymes and subsequent metabolic overflow. Metabolic overflow occurs during microbial fermentation when enzymatic complexes in oxidative phosphorylation and tricarboxylic acid (TCA) cycle are saturated and excess carbon is secreted in the form of fermentation products such as ethanol, acetate, lactate, formate, or succinate (Sánchez et al., 2017). In such a case, the onset of fermentation is a rational step, as NAD^+^ is regenerated during it. In our measurements, however, the main secreted metabolite was pyruvate – a substance that is still oxidized. As there were no straightforward options on how to further verify our hypothesis experimentally, we decided to employ mathematical modeling of *P. putida* metabolism. We assembled a GECKO model from the most actual *P. putida* genome-scale metabolic model (GEM) iJN1463 (Nogales et al., 2020) (**Materials and methods, Supplementary methods**). The upgraded model integrated kinetic data and protein concentrations (Domenzain et al., 2021; Sánchez et al., 2017). As *P. putida* metabolic network is difficult to simulate (Kohlstedt and Wittmann, 2019), first, a set of different simulation objectives was investigated to test the new model. To that end, we used a difference between normalized Euclidean distances of experimentally measured fluxes and simulated fluxes. In total, 18 simulation objectives were tested. Five appeared to be the best: (i) maximizing the production rate of CO_2_, (ii) maximizing the uptake of glucose, (iii) maximizing the growth rate, and (iv)+(v) maximizing the NADH and NADPH production rates (**Table S6**). Because the distance between maximizing NADH and NADPH production rate was found to be as low as 0.0022, just the NADPH production objective was chosen for further analysis.

Subsequently, a model representing the Δ*gcd* mutant was constructed by setting the upper bound of the exchange reaction for the Gcd protein (Uniprot ID Q88MX4 for glucose dehydrogenase) to 0. Since we did not detect any gluconate or ketogluconate in the cultures of Δ*gcd* mutants, these exchange reactions were constrained to 0. In contrast, exchange reactions for pyruvate and acetate were left unconstrained. The growth rate of the Δ*gcd* mutant model, calculated with the use of Flux Balance Analysis (Orth et al., 2010), was reduced compared to the growth rate of the wild-type model (0.45 h^-1^ *vs*. 0.68 h^-1^) which corresponded well with the experimentally measured values (**Table 2)** (del Castillo et al., 2007; Elmore et al., 2020, p. 202; Poblete-Castro et al., 2013). Interestingly, *in silico* overexpression of only one gene PP_1011 encoding Glk glucokinase – the first enzyme in the direct glucose phosphorylation pathway - almost completely restored growth rate to the value observed in the wild-type model (0.67 h^-1^ *vs*. 0.68 h^-1^). This result was surprising because our mutant hexGCDbglC_glf_lacY, with a de-repressed glucokinase gene, grew slower than the strain GCDbglC_glf_lacY (**Table 3**). Our presumption that glucokinase was not a limiting enzyme in our strains was nonetheless confirmed by the fact that the Δ*gcd* model was not able to produce pyruvate and acetate. To create the Δ*gcd* Δ*hexR* model, data from Δ*hexR* mutant were integrated into the Δ*gcd* model (del Castillo et al., 2008). The resulting model had growth rate almost identical to that of the wild-type model (0.67 h^-1^). The model thus showed that the additional deletion of Δ*hexR* regulator should fully restore the growth rate of the Δ*gcd* mutant on glucose. However, neither the original GEM nor our GECKO model included kinetic and proteomic data for transporters. This may explain the discrepancy between the *in-silico* simulations and our experimental observations. Also, all our strains in experiments were precultured on glucose. This step might result in the adaptation of Δ*gcd* mutant and derepression of HexR-controlled genes.

To study the effect of uncontrolled glucose uptake in the GECKO model, we gradually increased the glucose uptake rate in the Δ*gcd* Δ*hexR* model. Then, we simulated the growth rate and used it as a lower bound in the next simulation where one of four cellular objectives (growth, glucose uptake rate, CO_2_ production rate, or NADPH production rate) was maximized. The results of the simulations are shown in **Figure 5**. All simulations showed that with increasing glucose uptake rate, the growth rate increases linearly, but, after reaching the value of ∼ 0.67 h^-1^, the metabolism is saturated with glucose. Subsequently, due to the excess of carbon in the model, the growth rate decreases and organic acids pyruvate and acetate begin to accumulate. The fact that the same scenario was obtained for all four selected simulation objectives highlights the robustness of the analysis.

**Figure 5.**
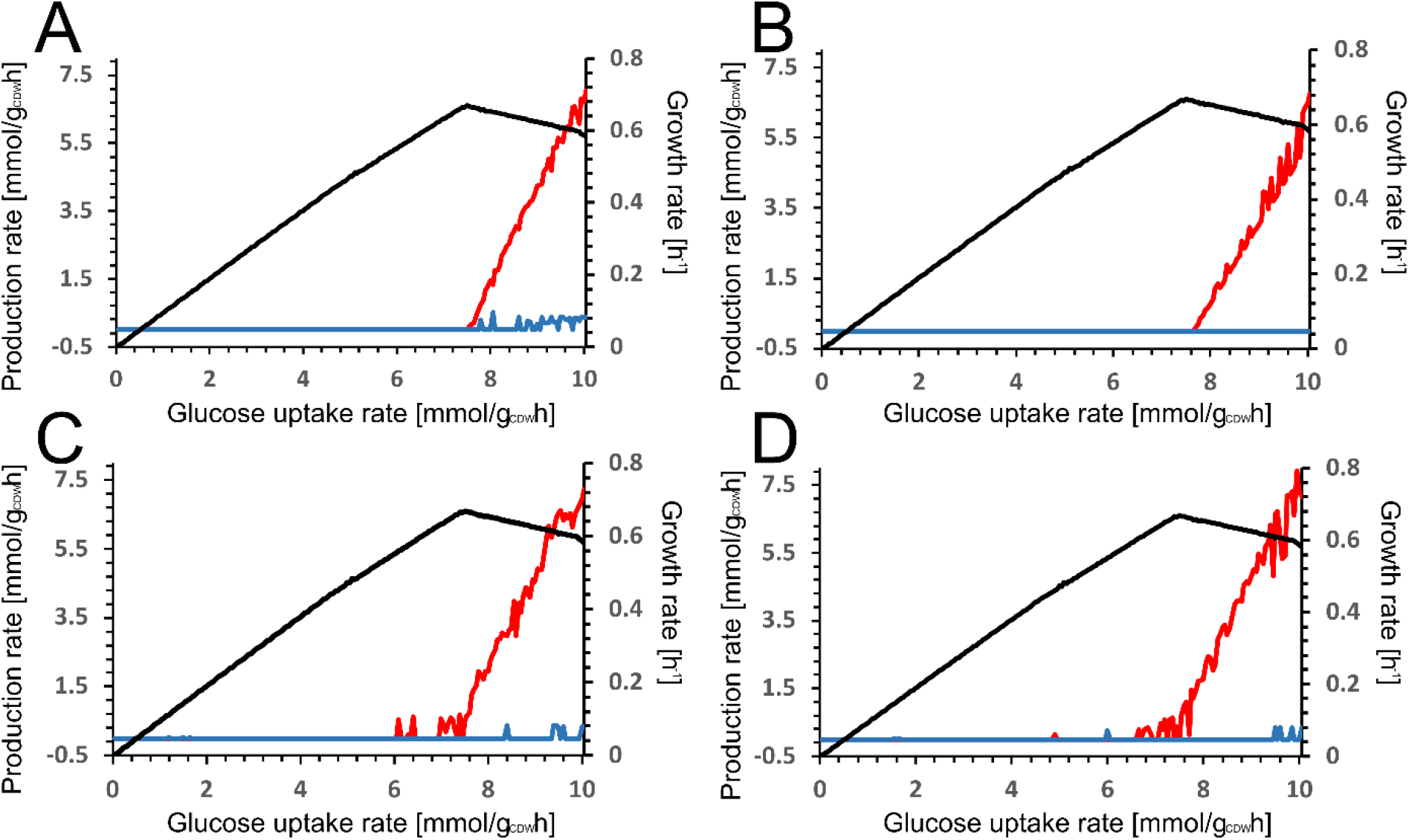
Simulations with a model adapted for *hexR* and *gcd* deletions. First, a defined glucose uptake rate was set (x-axis). The maximal growth rate was simulated and this value was used as a lower bound for subsequent simulation of (**A**) growth rate, (**B**) maximal CO2 production rate, (**C**) maximal glucose uptake rate, (**D**) maximal NADPH production rate. Growth rate, black line; pyruvate production rate, red line; acetate production rate, blue line.

These simulations correctly predicted that the major overflow metabolite is pyruvate, and acetate is also produced to a lesser extent. As can also be seen in **Figure 5**, pyruvate production was dependent on the growth rate of the bacterium, which agreed with our data. The GCDbglC_glf_lacY strain cultured on glucose reached a growth rate of 0.43 h^-1^ and produced pyruvate to a maximum concentration of 1.4 g/L (**Figure 3A**). In contrast, when the same strain was cultured on cellobiose with a growth rate of 0.31 ± 0.01 h^-1^, no accumulation of pyruvate was observed (**Figure S14)**. Also, slower-growing hexGCDbglC_glf_lacY mutant (µ = 0.36) accumulated no acids.

There was a partial discrepancy between the data from the Δ*gcd* Δ*hexR* model and the data from the experiments. The glucose uptake rate necessary to saturate the metabolism of the Δ*gcd* Δ*hexR* model was ∼7.8 mmol/g_cdw_/h, but our experiments on glucose substrate (Figure 3C and 3D) showed that pyruvate is accumulated even at 6.14 mmol/g_cdw_/h. This can be explained by the fact that samples for proteomic analysis were collected at a growth rate of 0.68 h^-1^ – higher than the maximum growth rate of WT strain on glucose (0.52 h^-1^) measured here. Deletions of *gcd* and hexR may also have an effect on the proteome composition (del Castillo et al., 2008, 2007). Taken together, the computational simulations using the GECKO model of *P. putida* metabolism showed that the observed production of pyruvate and acetate in aerobic cultures of mutants with exogenous transporters is most probably caused by the deregulation of substrate uptake and saturation of catabolic enzymes. One of the saturated enzymes was pyruvate dehydrogenase which indicates a potential bottleneck in this step. Insufficient activity of this multienzyme complex in *P. putida* (Ebert et al., 2011) might be exacerbated by other factors such as product inhibition with NADH (Bisswanger, 1974; Moxley and Eiteman, 2021). A massive flux of carbon through the direct phosphorylation route in GCDbglC_glf_lacY mutant may lead to the enhanced production of NADH in glucose 6-phosphate 1-dehydrogenase (Zwf) step (**Figure 1)** (Volke et al., 2021) and lack of NAD^-^ required for the pyruvate dehydrogenase reaction. Excess acetate (˃ 60 % of it, as shown by the model) likely comes from L-glutamate formed from α-ketoglutarate in the tricarboxylic acid cycle (**Figure 1**) which is converted by the activities of NAD(P)-specific glutamate dehydrogenase (PP_2080, PP_0675), acetylornithine aminotransferase (PP_4481, PP_0372), and acetylornithine deacetylase (PP_5186, PP_3571). The first reaction contributes to NAD(P)^+^ regeneration.

### 3.8 New GECKO model highlights the potential of engineered P. putida for the biosynthesis of attractive chemicals

The GECKO model was eventually used to evaluate the potential of the GCDbglC_glf_lacY strain to produce valuable chemicals from excess pyruvate. Simulations were carried out to test the formation of six compounds whose bioproduction in *P. putida* was experimentally verified in this and other studies: ethanol, lactate, medium chain length polyhydroxyalkanoates (mcl-PHA), rhamnolipids, β-carotene, and *cis*, *cis*-muconate (Bentley et al., 2020; Johnson and Beckham, 2015; Nikel and de Lorenzo, 2014, p. 201; Poblete-Castro et al., 2013; Sánchez-Pascuala et al., 2019; Tiso et al., 2020). A sink reaction for each of these substances was set as an objective function in simulations. In the case of β-carotene, ethanol, and lactate, missing reaction(s) had to be added to the model (see **Supplementary methods** in **Supplementary material** for model details). Simulation of mcl-PHA production was more complicated as it is not a single defined substance but a polymer of different lengths (Kavitha et al., 2018). The synthesis takes place in cycles with acetyl-coenzyme A as the main substrate. Thus, acetyl-coenzyme A level was used as a proxy for the production of mcl-PHA.

The analyses were performed with three models – a wild-type model (growth rate: 0.678 h^-1^ and glucose uptake rate 8.71 mmol/g_CDW_/h), a Δ*gcd* Δ*hexR* mutant model which is not stressed by a high glucose uptake rate (growth rate: 0.665 h^-1^ and glucose uptake rate 7.48 mmol/g_CDW_/h) and the same strain but with a glucose uptake rate set at 10.05 mmol/g_CDW_/h and growth rate at 0.582 h^-1^. The results of all calculations are summarized in **Table 4**. The calculations were in agreement with our previous experimental observations when they correctly predicted significantly increased productivity of ethanol and lactate in cells stressed with high glucose uptake when compared with WT. For both L-lactate and ethanol the trend was the same. WT model produced more chemical than Δ*gcd*/Δ*hexR* model but once this model was stressed with high glucose uptake it produced more L-lactate and ethanol than WT model. The fact that WT model produces more ethanol or lactate than Δ*gcd*/Δ*hexR* model (note that the same trend for ethanol production was observed also in our experiments, **Figure 4**), points to the slower growth of Δ*gcd* mutant and to the important role of the peripheral glucose oxidation pathway in the regulation of NAD(P)H/ NAD(P)^+^ ratio (Nikel et al., 2021). Since the synthesis of both fermentation products also involves the regeneration of NAD^+^, it is necessary to set the optimal ratio of fluxes through the peripheral oxidative pathway consuming NADPH in the step of 2-ketogluconate-6-phosphate conversion to 6-phosphogluconate and through the direct oxidation route which in turn produces NAD(P)H (**Figure 1**).

**Table 4.** Production rates of selected bioproducts in wild-type and engineered *P. putida* calculated with the use of the prepared metabolic models.

| Product | Model WT | Model $\Delta gcd/\Delta hexR$ | Model $\Delta gcd/\Delta hexR$ stressed<sup>a</sup> |
| --- | --- | --- | --- |
| **Ethanol** | 1.877 mmol/g<sub>CDW</sub>/h | 0.067 mmol/g<sub>CDW</sub>/h | 5.743 mmol/g<sub>CDW</sub>/h |
| **L-lactate** | 0.815 mmol/g<sub>CDW</sub>/h | 0.513 mmol/g<sub>CDW</sub>/h | 4.376 mmol/g<sub>CDW</sub>/h |
| **Rhamnolipids** | 0.0499 μmol/g<sub>CDW</sub>/h | 0.0536 μmol/g<sub>CDW</sub>/h | 0.0763 μmol/g<sub>CDW</sub>/h |
| **Cis, cis-muconic acid** | 0.074 mmol/g<sub>CDW</sub>/h | 0.074 mmol/g<sub>CDW</sub>/h | 0.074 mmol/g<sub>CDW</sub>/h |
| **Acetyl-coenzyme A<sup>b</sup>** | 0.489 μmol/g<sub>CDW</sub>/h | 0.489 μmol/g<sub>CDW</sub>/h | 0.489 μmol/g<sub>CDW</sub>/h |
| **β-carotene** | 0.066 μmol/g<sub>CDW</sub>/h | 0.078 μmol/g<sub>CDW</sub>/h | 0.103 μmol/g<sub>CDW</sub>/h |

<sup>a</sup>High glucose uptake rate stress this model
<sup>b</sup>Serves as a proxy value for mcl-PHA synthesis

Concerning the remaining tested bioproducts, it was concluded that our interventions in *P. putida* metabolism should theoretically have no effect on the production of acetyl-coenzyme A and *cis*, *cis*-muconate. On the other hand, the production of rhamnolipids was improved by 53%. As can be seen from the data (**Table 4**), *in silico* production was increased both after deletion of *gcd* and after the increase in glucose uptake rate. β-carotene is produced from pyruvate and glyceraldehyde-3-phosphate. Both are generated in the upper glucose metabolism and thus carbon saturation should lead to higher productivities. Just as for rhamnolipid synthesis, the WT model produced the least β-carotene and the Δ*gcd*/Δ*hexR* model stressed with high glucose uptake formed the most (56% more than the WT model).

Taken together, the model showed correlation with our experimental data and identified ethanol and L-lactate as attractive target products that can be formed more efficiently by our engineered strain GCDbglC_glf_lacY than by wild-type *P. putida*. Based on the model, the recombinant strain also holds promise as a potent producer of other pyruvate-derived chemicals such as rhamnolipids and β-carotene. The model is freely available at GitHub and can be used as a valuable predictive tool in future metabolic engineering studies.

## 4. Conclusions

In this work, we intended to install efficient co-metabolism of glucose and cellobiose in *P. putida* EM42 for the bioproduction of valuable compounds. Cultivation experiments corroborated that the blockage of the periplasmic glucose oxidation pathway via the deletion of glucose dehydrogenase *gcd* gene enables co-utilization of the two sugars. Kinetic measurements indicated that the deletion relieved the inhibition of β−glucosidase by some of the metabolites from the peripheral glucose pathway – most probably glucono-δ-lactone and gluconate. Our presumption that a null mutation of HexR local transcriptional regulator could correct the growth defect of Δ*gcd* mutant on glucose was not confirmed. Instead, the reduced growth rate was partially compensated by the implantation of exogenous glucose (Glf) and cellobiose (LacY) transporters. To the best of our knowledge, this is the first report showing that the growth of *P. putida* Δ*gcd* mutant on glucose is, at least partially, limited by sugar transport. This finding is instrumental for any studies that will use Δ*gcd* mutant to enhance the bioproduction capacity of *P. putida* and some other pseudomonads on glucose or other sugars that can be oxidized by Gcd (Bentley et al., 2020; Blombach et al., 2022; Dvořák and de Lorenzo, 2018; Elmore et al., 2020; Poblete-Castro et al., 2013). Our experimental and computational results also show that simple overexpression of the transporters will not result in a higher growth rate. Instead, a balancing of expression is necessary – here achieved via random insertions of transporter genes into *P. putida* chromosome. We observed an accumulation of pyruvate and acetate in aerobic cultures of *P. putida* mutant GCDbglC_glf_lacY selected after introducing the sugar transporters and we demonstrated that the excess pyruvate formed from glucose and cellobiose can be streamed into ethanol or L-lactate after harnessing the strain with exogenous biosynthetic pathways. Another selected *P. putida* mutant, hexGCDbglC_glf_lacY with transporters inserted into Δ*gcd* Δ*hexR* background, did not accumulate organic acids but displayed improved co-utilization of glucose and cellobiose when compared with the parental strain.

To check our hypothesis that the overproduction of acids was caused by the unregulated inflow of carbon from sugar substrates, we prepared a GECKO metabolic model representing the Δ*gcd* mutant constrained with proteomic and kinetic data. The model showed that the glucose uptake rate increased above a certain threshold saturates *P. putida*’s metabolism and it correctly predicted pyruvate as a major overflow metabolite. The concrete cause of pyruvate overflow remains unclear and further investigations are needed to specify the mechanism underlying aerobic accumulation of acids in GCDbglC_glf_lacY strain and its absence in hexGCDbglC_glf_lacY mutant. In any case, experimental data and theoretical calculations performed in this study suggest that the GCDbglC_glf_lacY strain will find its use in the biosynthesis of pyruvate-derived products from cellulosic sugars. In conclusion, this work unveils the prerequisites for the efficient glucose and cellobiose co-utilization in *P. putida* and introduces a new strategy for pyruvate overproduction in aerobic bacterial cultures as well as an upgraded metabolic model of *P. putida* that will be instrumental for further engineering of this attractive microbial host.

