## Supplementary materials for "Engineering of *Pseudomonas putida* for accelerated co-utilization of glucose and cellobiose yields aerobic overproduction of pyruvate explained by an upgraded metabolic model"

by

Dalimil Bujdoš[1], Barbora Popelářová[1], Daniel Volke[2], Pablo. I. Nikel[2], Nikolaus Sonnenschein[3], Pavel Dvořák[1*]

*[1]Department of Experimental Biology (Section of Microbiology), Faculty of Science, Masaryk University, Kamenice 753/5, 62500, Brno, Czech Republic.*

*[2] The Novo Nordisk Foundation Center for Biosustainability, Technical University of Denmark, Kemitorvet 220, 2800 Kongens Lyngby, Denmark*

*[3]Department of Biotechnology and Biomedicine, Technical University of Denmark, Building 223, Søltofts Plads, 2800 Kgs. Lyngby, Denmark*

* Corresponding author:

Dr. Pavel Dvořák

Department of Experimental Biology (Section of Microbiology), Faculty of Science

Masaryk University, Kamenice 735/5, Brno 62500, Czech Republic

### Content

***Supplementary methods***

***Supplementary results and discussion***

***Supplementary tables***

**Table S1.** Final concentrations of antibiotics used in *Escherichia coli* and *Pseudomonas putida* cultures in this study.

**Table S2.** Oligonucleotide primers used in this study.

**Table S3.** Tn5-driven gene insertions in *P. putida* strains mapped by arbitrary PCR.

**Table S4.** Input parameters for the GECKO pipeline.

**Table S5.** The list of manually curated kinetic data for protein constrained (pc) and enzyme constrained (ec) model of *P. putida* metabolism.

**Table S6.** Analysis of different cellular objectives with enzyme constrained and protein constrained models.

***Supplementary figures***

**Figure S1.** SDS-polyacrylamide gel with samples from purification of BglC.

**Figure S2.** Michaelis & Menten plot for enzyme BglC with substrate 4-nitrophenyl β-D-glucopyranoside.

**Figure S3.** Screening of possible inhibitors of enzyme BglC.

**Figure S4.** Measurement of BglC activity with substrate 4-nitrophenyl β-D-glucopyranoside in presence of various concentrations of D-glucono-δ-lactone.

**Figure S5** Comparison of the growth of engineered *P. putida* strains in M9 minimal medium with glucose (4 g/L) in 48 well-plate format.

**Figure S6.** The first round of screening of *glf* mutants in 96-well plate format.

**Figure S7.** The second round of screening of *glf* mutants in 96-well plate format.

**Figure S8.** Screening of the library of *lacY* insertion mutants in 48-well plate format.

**Figure S9.** Comparison of the growth of engineered *P. putida* strains in M9 minimal medium with the mixture of glucose (4 g/L) and cellobiose (4 g/L) in shake flasks.

**Figure S10.** Cellobiohydrolytic activity in culture supernatants collected at multiple time points during shake flask culture of strain GCDbglC_glf_lacY on the mixture of glucose and cellobiose (4 g/L each).

**Figure S11**. Comparison of the growth of HexGCDglf and hexGCDglf_colony2 in M9 minimal medium with glucose (4 g/L) in shake flasks.

**Figure S12**. Shake flask culture of GCDglf colony 2 in M9 minimal medium with 8 g/L glucose.

**Figure S13.** Shake flask culture of *P. putida* WTbglC in M9 minimal medium with 8 g/L glucose.

**Figure S14**. Shake flask culture of GCDbglC_glf_lacY in M9 minimal medium with 8 g/L cellobiose.

***Nucleotide sequences of genes used in this study***

***References***

#### Supplementary methods

##### ***Purification of β-glucosidase BglC***

Strain *P. putida* EM42 pSEVA238_*bglC* was used for the BglC protein production. The plasmid pSEVA238 has an inducible expression system *xylS/Pm* (Gawin et al., 2017). BglC was tagged on its N-terminus with a 6x histidine anchor and Ni-NTA metal affinity chromatography was used for the purification of the protein.

The overnight culture was prepared by inoculating strain *P. putida* EM42 pSEVA238_*bglC* to 10 mL of LB medium with kanamycin. From this pre-culture, 200 mL of LB medium with kanamycin in a 1 L flask was inoculated to the starting OD of 0.05. The cells were cultured for 3 h with agitation (170 RPM) in Lab Companion IS-971 incubated orbital shaker (Jeio Tech) until the culture reached $OD_{600}$ of ~ 0.5. The protein production was then induced by the addition of 3-methyl benzoate to the final concentration of 1 mM. After the induction, the cells were grown for another 6 h. The cells were then collected (3,500 g, 10 min) and washed twice with purification buffer A (25 mM Tris-HCl, 250 mM NaCl, 5 mM imidazole, pH 7.5). To lyse cells and extract proteins the following components were added to the pellet: 3.5 mL of B-PER Bacterial Protein Extraction Reagent (ThermoFisher Scientific), 3.5 mL of buffer A, 3.5 µL of lysozyme (50 mg/mL), 3.5 µL of Dnase I (2500 U/mL) and $\frac{1}{4}$ of tablet cOmplete, Mini, EDTA-free Protease Inhibitor Cocktail (Roche). Cells were left to lyse at RT for 30 min with constant slow tilting. The lysate was centrifuged (19,720 g, 30 min, 4 °C). Supernatant was collected and mixed with an equal volume of pre-cooled buffer A. Subsequently the cell-free extract was filtered through 0.45 µm, 25 mm LUT Syringe Filters MCE (Labicom). The concentration of protein was measured by the Bradford method (Bradford, 1976) using Bradford Reagent (Sigma-Aldrich). The BglC was purified from the cell-free extract using fast protein liquid chromatography (NGC chromatography Scout system, Bio-Rad). The HisTrap High Performance 5 mL column (GE Healthcare) was connected to the system and washed with buffer A to equilibrate. A linear gradient of imidazole prepared by mixing buffer B (same composition as buffer A but with 50 mM imidazole) with buffer C (same composition as buffer A but with 500 mM imidazole) was used to elute the protein. A 30 kDa cut-off spin concentrator (Agilent Technologies) was used to concentrate the protein and change buffer to sodium phosphate buffer (5.38 g/L $NaH_2PO_4.H_2O$ and 16.35 g/L $Na_2HPO_4.7H_2O$ filled up to 1 L with milliQ $H_2O$). BglC was purified to homogeneity and the enzyme was divided into several aliquots and stored at -20 °C.

***GECKO model construction***

Using average measured enzyme concentration ($c_i$ g/$g_{prot}$) and concentration of all proteins ($P_{tot}$, $g_{prot}/g_{CDW}$), turnover numbers were calculated from specific activities ($SA_{CFEi}$) measured in the cell-free extract of *P. fluorescens* E-20 (Equation 1) (Lynch et al., 1975).

$$k_{cat\,i}\,[s^{-1}] = \frac{SA_{CFEi}[\mu mol/min/mg] \cdot \frac{1}{c_i\,[g/g_{prot}]} \cdot MW_i[mg/\mu mol]}{60} \quad \textbf{(1)}$$

Here $k_{cat\,i}$ is the turnover number if enzyme *i*, $c_i$ is a concentration of enzyme *i* with a unit gram of measured enzyme per gram of total protein, $MW_i$ is the molar weight of protein *i*.

During the initial simulations of pc models, ATP synthase appeared to be the most growth-limiting reaction. The measured $k_{cat}$ from ATP synthase in *E. coli* is low (16.6 $s^{-1}$) (Iino et al., 2009) compared to, for example, that used in the enzyme constrained model of *S. cerevisiae* (120 $s^{-1}$) (Sánchez et al., 2017). In the pc model, flux is forced through other less efficient reactions for ATP synthesis and proper growth rate and glucose uptake rate could not be achieved. The measured concentrations of enzymes in the ATP synthase reaction are not satisfactory either and proteins Q88BW8 (ATP synthase subunit a) and Q88BW9 (ATP synthase subunit c) are missing entirely. For ec model we decided to fit the proper $k_{cat}$ using a classical GEM. The original model iJN1463 was constrained by the measured glucose uptake rate (7.7457 mmol/$g_{CDW}$/h) and gluconate and ketogluconate production rates ($q_{exp}$). These two rates were not measured but taken from the study of Kohlstedt and Whittmann (Kohlstedt and Wittmann, 2019). The growth rate reported therein ($\mu_x$) was only 0.55 $h^{-1}$ and the production rates were thus rescaled using Equation 2.

$$q_{exp}[mmol/g_{CDW} \cdot h] = \frac{\mu_{exp}[h^{-1}]}{\mu_X\ [h^{-1}]} q_X[mmol/g_{CDW} \cdot h] \quad \textbf{(2)}$$

Where $q_X$ are the production rates reported (Kohlstedt and Wittmann, 2019) and $\mu_{exp}$ is the experimentally measured growth rate (0.6781 $h^{-1}$).

The growth rate simulated by the constrained GEM closely corresponded to the experimentally measured growth rate and thus flux through ATP synthase in this simulation was used to

$$k_{cat}[h^{-1}] = \frac{f_{ATPS}[mmol/g_{CDW} \cdot h]}{min(AvgC_i[mmol/g_{CDW}] + (1.96 \cdot SD)) \cdot \frac{1}{S_i}} \quad \textbf{(3)}$$

calculate the fitted $k_{cat}$ using Equation 3.

Where $f_{ATPS}$ is the flux through ATP synthase reaction in the GEM, $AvgC_i$ is an average molar concentration of subunit $i$, $S_i$ is the number of these subunits in ATP synthase. The expression 1.96SD (SD = standard deviation of the protein measurement) then ensures that low measurements do not overconstrain the model. The fitted $k_{cat}$ for ATP synthase was 223.19 $s^{-1}$.

Fluxomic data for *P. putida* KT2440 was collected for the purpose of this study. This was used for additional parametrization of the model. Two reactions were omitted: (i) glucose-6-

phosphate dehydrogenase, because the turnover numbers were recently measured and are reliable (Volke et al., 2021) and (ii) malate dehydrogenase, because this is a lumped flux from two different reactions: $NAD^+$ dependent malate dehydrogenase (MDH) and ubiquinone dependent malate dehydrogenase (MDH2). Both enzymes were present in data sets and thus either reaction could not be left considered. The fluxomic data were used to ensure that the turnover numbers in the model, combined with the measured enzyme concentrations, could carry the measured flux. This was governed by two functions fitToFluxes which fits $k_{cat}$ into most reactions and fixArm which fits $k_{cat}$ into arm reactions *i.e.*, reactions with isoenzymes. Both are embedded and called from GECKO\geckomat\change_model\manualModifications.m. The fitting process is run before modifyKcats.m and is applied to only those reactions where all proteins were measured. The function first checks if this inequality exists:

| | |
|---|---|
| $k_{cat\ i}[h^{-1}] \cdot min(AvgC_{ij}[mmol/g_{CDW}] + (1.96 \cdot SD)) < f_{exp\ i}$ | **(4)** |

Where $k_{cat\ i}$ is the turnover number of enzyme *i*, $AvgC_{ij}$ is the average concentration of subunit *j* of an enzyme *i* and $f_{exp\ i}$ is the experimentally measured flux of a reaction that enzyme *i* catalyzes.

The function loops through every subunit in proteins in all reactions and identifies the lowest value that satisfies the condition above. If such value exists, then new $k_{cat}$ is fitted using Equation 5. which is a general version of Equation 3.

| | |
|---|---|
| $k_{cat}[h^{-1}] = \dfrac{f_{exp\ i}[mmol/g_{CDW} \cdot h]}{\min\limits_{j}\ (AvgC_{ij}[mmol/g_{CDW}] + (1.96 \cdot SD))}$ | **(5)** |

The new $k_{cat}$ is set to every protein participating in the reaction.

Reactions that are catalyzed by multiple isoenzymes must be treated differently. The burden of the measured flux can be distributed onto multiple isoenzymes and each individual isoenzyme may not be able to provide the entire flux. If the combined maximal calculated flux is not sufficient to carry the measured flux the burden is distributed based on the measured concentration. Therefore, if the parameters (concentration and $k_{cat}$) satisfy the condition given by the inequality in Equation 6 then a new $k_{cat}$ is calculated to match the measured flux.

$$\sum_i \left( k_{cat\ i}[h^{-1}] \cdot min(AvgC_{ij} \left[\frac{mmol}{g_{CDW}}\right] + (1.96 \cdot SD)) \right) < f_{exp\ i} \quad \textbf{(6)}$$

Where $k_{cat\ i}$ is the turnover number of enzyme $i$, $AvgC_{ij}$ is the average concentration of subunit $j$ in an enzyme $i$ and $f_{exp\ i}$ is the experimentally measured flux of a reaction that enzyme $i$ catalyzes. $\Sigma_i$ is then the sum of all fluxes that the isoenzymes are able to provide.

New $k_{cat}$ is calculated based on the molar fraction of the individual enzymes. It is therefore assumed that the flux $f_{exp\ i}$ will be distributed into the isoenzymes based on their measured concentrations. By modifying Equation 6 we get Equation 7 for the calculation of the new $k_{cat}$.

$$k_{cat}[h^{-1}] = \frac{f_{exp\ i}[mmol/g_{CDW} \cdot h] \cdot \frac{k_{cat\ i}[h^{-1}]\ \cdot\ min(AvgC_{ij} \left[\frac{mmol}{g_{CDW}}\right] + (1.96 \cdot SD))}{\sum_i \left( k_{cat\ i}[h^{-1}] \cdot min(AvgC_{ij} \left[\frac{mmol}{g_{CDW}}\right] + (1.96 \cdot SD)) \right)}}{min(AvgC_{ij}[mmol/g_{CDW}] + (1.96 \cdot SD))} \quad \textbf{(7)}$$

Where the variables are the same as in Equation 6.

The fraction multiplying $f_{exp\ i}$ can be interpreted as the flux that the given isoenzyme can ensure divided by the flux that all isoenzymes can ensure. Since $k_{cat}$ assigned by the pipeline to individual isoenzymes is the same. In most cases, the newly fitted $k_{cat}$ will be likewise the same for all isoenzymes as the only variables determining the fraction are concentrations of the enzymes.

The list of all $k_{cat}$ values that were obtained this way is listed in GECKO\databases\fittedKcats.csv. Unlike turnover numbers in **Table S5**. These fitted $k_{cat}$ values can be increased in subsequent parametrization by modifyKcats.m as they ensure only minimal flux.

***Modifications of the GECKO pipeline.*** We have made minor changes to how the pipeline works. Absolute protein concentration ($P_{tot}$ [$g_{protein}/g_{CDW}$]) was calculated from the biomass reaction. Stoichiometric coefficients in this reaction represent the composition of the biomass of cells and have a unit mmol/$g_{CDW}$. Using the Equation 8. and MW of each amino acid we determined $P_{tot}$ to be equal to 0.58666 $g_{prot}/g_{CDW}$.

$$P_{tot}\left[g_{prot}/g_{CDW}\right] = \frac{|X_i|[mmol/g_{CDW}]}{1000} \cdot \left(MW_i\left[\frac{g}{mol}\right] - 18\right) \quad \textbf{(8)}$$

Where $X_i$ is a stoichiometric coefficient of amino acid *i* in biomass reaction, $MW_i$ is the molar weight of amino acid *i* and 18 is the molar weight of water molecule which is lost upon the formation of the peptide bond. This calculation is done by the function GECKO\geckomat\limit_proteins\sumProtein.m. For our models, it was not necessary to rescale the biomass reaction during proteome integration since the sum of all measured proteins (including those missing in the model) was set to be equal to $P_{tot}$. However, in simulations, the model also uses other enzymes that have not been measured. This leads to a formal problem where the model produces fewer amino acids than it uses in the form of proteins.

The function modifyKcats.m was also changed. Previously it could happen that $k_{cat}$ of a protein was changed and if the protein belonged to an enzyme complex the rest of the proteins would maintain the old $k_{cat}$. The function was changed so that the newly found $k_{cat}$ is assigned to every protein in a reaction. The second problem was that the function could get stuck in an endless loop if the new $k_{cat}$ is the same as the old value. This issue was also resolved.

***Modifications of constructed metabolic model for the calculation of production rates of selected bioproducts***

Except for mcl-PHA, all the simulated products are produced during active cell growth, and the growth rate of these models was thus set. Mcl-PHA are produced during nitrogen or sulfur limitation (Prieto et al., 2016). Hence, model growth and glucose uptake rate were left unconstrained. Also, proteomic data for *P. putida* KT2440 from the study of Mozejko-Ciesielska and Sefarim (2019) was used in this particular case because the composition of the proteome in these and in log phase conditions fundamentally differs (Dabrowska et al., 2020; Możejko-Ciesielska and Serafim, 2019).

The synthesis of L-lactate from pyruvate is carried out by only one enzyme L-lactate dehydrogenase. Such conversion was previously established in *P. putida* KT2440, by implanting mutant L-lactate dehydrogenase from *Bos taurus* (Johnson and Beckham, 2015). The synthesis of ethanol from pyruvate in *P. putida* requires the presence of enzyme pyruvate decarboxylase which converts pyruvate to acetaldehyde. Acetaldehyde is then converted by alcohol dehydrogenate to ethanol. The genome of *P. putida* KT2440 does not contain a functional pyruvate decarboxylase and thus this reaction was added to the model. *P. putida* has two putative alcohol dehydrogenases Q88G86 and Q88MF5. Nikel and de Lorenzo (2014), however, expressed exogenous alcohol dehydrogenase from *Zymomonas mobilis* to establish ethanol production (Nikel and de Lorenzo, 2014). In our simulations, the two alcohol dehydrogenase reactions were left unconstrained and the proteins were removed from the reactions. *P. putida* is able to produce rhamnolipids without any additional modifications (Tiso et al., 2020). In our simulations, the production of rhamnosyl-alpha-1-3-N-acetyl undecaprenyl diphosphate was used as an evaluation of the production of rhamnolipids. *Cis, cis* muconate is an intermediate in *P. putida* metabolism (Bentley et al., 2020). Thus, for the production of this compound, no additional reactions had to be added. The last tested product was β-carotene. *P. putida* is unable to synthesize this compound but has all the enzymes to produce two main precursors isopentenyl pyrophosphate and farnesyl pyrophosphate. The rest of the pathway is linear (Shi et al., 2014) and was thus added to the models.

#### Supplementary results and discussion

***Enzyme BglC is inhibited by metabolites in the peripheral glucose pathway***

D-gluconate and D-glucono-δ-lactone were used for the determination of the modality of inhibition and the measurement of dissociation constants. Results obtained from gluconate inhibition could be described by neither competitive, non-competitive nor mixed-type inhibition. Gluconate may inhibit BglC only because it is in dynamic equilibrium with the gluconolactone and its respective sodium salt (Sawyer and Bagger, 1959). Another contributing factor is that inhibitions were measured at very high concentrations of gluconate with sodium gluconate being used for the preparation of the assay. Thus, high sodium concentrations could also affect the activity of BglC.

#### Supplementary tables

**Table S1.** Final concentrations of antibiotics used in *Escherichia coli and Pseudomonas putida* cultures in this study.

| Antibiotic | *E. coli* (µg/mL) | *P. putida* (µg/mL) |
|---|---|---|
| **Kanamycin** | 50 | 50 |
| **Gentamicin** | 10 | 10 |
| **Streptomycin** | 50 | 60 |
| **Ampicillin** | 150 | 500 |
| **Chloramphenicol** | 30 | 30 |

**Table S2.** Oligonucleotide primers used in this study.

| Name | Sequence (5'→3')[a] | Source |
|---|---|---|
| lacY for PstI RBS | AAACTGCAGCTTTAAG**AAGGAG**ATATACAT ATGTACTATTTAAAAAACACAAACTTTTGG | This study |
| lacY rev HindIII | AAAAAGCTTTTAAGCGACTTCATTCACCTG | This study |
| glf for PstI | ATAACTGCAGAATTCAGGAGGAAAAACATA TG | This study |
| glf rev HindIII | ATTAATAAGCTTTTACTTCTGGGAGCG | This study |
| Glf seq fw | CACGGTGAAGGACAGCG | This study |
| ARB6 | GGCACGCGTCGACTAGTACNNNNNNNNNN ACGCC | (Martínez-García et al., 2014) |
| ARB2 | GGCACGCGTCGACTAGTAC | (Martínez-García et al., 2014) |
| ME-I-Sm-Ext-R | ATGACGCCAACTACCTCTGATA | (Martínez-García et al., 2014) |
| ME-I-Sm-Int-R | TCACCGCTTCCCTCATGATGTT | (Martínez-García et al., 2014) |
| ME-O-Sm-Ext-F | CTTGGCCTCGCGCGCAGATCAG | (Martínez-García et al., 2014) |
| ME-O-Sm-Int-F | CACCAAGGTAGTCGGCAAAT | (Martínez-García et al., 2014) |
| ME-O-Gm-Ext-F | GCACTTTGATATCGACCCAAGT | (Martínez-García et al., 2014) |
| ME-O-Gm-Int-F | TCCCGGCCGCGGAGTTGTTCGG | (Martínez-García et al., 2014) |

| | | |
|---|---|---|
| ME-I-Gm-Ext-R | GTTCTGGACCAGTTGCGTGAG | (Martínez-García et al., 2014) |
| ME-I-Gm-Int-R | GAACCGAACAGGCTTATGTCA | (Martínez-García et al., 2014) |
| oCJ123 | TTTGCCGTCACCCAGTAC | (Johnson and Beckham, 2015) |
| oCJ124 | TTTACCCACAGCGGCATG | (Johnson and Beckham, 2015) |
| LDHA EcoRI fw | AATGAATTCAGGAGGACAGCTATGGC | This study |
| LDHA SacI rv | AATGAGCTCTCAGAACTGCAGTTCCTTC | This study |
| PS1new | AGGGCGGCGGATTTGTC | This study |
| LDHA seq rv | GAAGCGGATATCCGGTTTG | This study |
| adhB seq fw | GACTACGAGAGCCAGACG | This study |
| pdc seq fw | CCAAGAAAGAGGACGTCC | This study |
| PS2new | CGGCAACCGAGCGTTC | This study |

[a] Restriction sites are underlined.

**Table S3.** Tn5-driven gene insertions in *P. putida* strains mapped by arbitrary PCR.

| Strain | Strain abbreviation | Locus tag | Gene name |
|---|---|---|---|
| | | ***glf* insertion sites** | |
| *P. putida* EM42::*glf* | WTglf | PP_5084 | Penicilin binding protein 1A |
| *P. putida* EM42 Δ*gcd*::*glf* colony 1 | GCDglf | PP_5410 | DeoR family transcription regulator |
| *P. putida* EM42 Δ*gcd*::*glf* colony 2 | – | PP_1204 | Hypothetical protein |
| *P. putida* EM42 Δ*hexR*::*glf* | hexWTglf | PP_0983 | LPS export ABC transporter permease LptG |
| *P. putida* EM42 Δ*hexR* Δ*gcd*::*glf* colony 1 | hexGCDglf | PP_1318 | Cytochrome *bc* complex cytochrome subunit |
| *P. putida* EM42 Δ*hexR* Δ*gcd*::*glf* colony 2 | – | PP_0909 | Hypothetical protein |
| | | ***lacY* insertion sites** | |
| *P. putida* EM42::*lacY* pSEVA2213_*bglC* | WTbglC_lacY | Not tested | |
| *P. putida* EM42::*glf*::*lacY* pSEVA2213_*bglC* | WTbglC_glf_lacY | Not tested | |
| *P. putida* EM42 Δ*gcd*::*glf*::*lacY* pSEVA2213_*bglC* | GCDbglC_glf_lacY | PP_0033 | Undecaprenyl-glycosyl transferase |
| *P. putida* EM42 Δ*hexR*::*lacY* pSEVA2213_*bglC* | hexWTbglC_lacY | PP_0961 | Toluene tolerance protein |
| *P. putida* EM42 Δ*hexR*::*glf*::*lacY* pSEVA2213_*bglC* | hexWTbglC_glf_lacY | PP_0619/PP_0620 | Branched chain amino acid transporter/GntR family transcription regulator |
| *P. putida* EM42 Δ*gcd* Δ*hexR*::*glf*::*lacY* pSEVA2213_*bglC* | hexGCDbglC_glf_lacY | PP_1802 | Glycosyltransferase |

**Table S4.** Input parameters for the GECKO pipeline.

| | |
|---|---|
| Expected enzyme saturation constant (σ) | 0.5 |
| Objective function | BIOMASS_KT2440_WT3 |
| The total concentration of protein | 0.58666 $g_{prot}/g_{CDW}$[a] |
| Ratio of enzymes/proteins | 0.5 |
| Indexes of reactions in oxidative phosphorylation and ATP synthase | NADH16pp, SUCDi, CYTBO3_4pp, CYO1_KT, ATPS4rpp |
| Index of non-growth associated reaction | ATPM |
| Substrate, Exchange reaction | Glucose, EX_glc__D_e |
| Target glucose uptake rate | 7.7457 mmol/$g_{CDW}$/h |
| Target growth rate | 0.6785 $h^{-1}$ |
| Gluconate production rate | 0.7398 mmol/$g_{CDW}$/h |
| Ketogluconate production rate | 0.1233 mmol/$g_{CDW}$/h |

[a] Calculated from biomass reactions

**Table S5.** The list of manually curated kinetic data for protein constrained (pc) and enzyme constrained (ec) model of *P. putida* metabolism.

| **Reaction name** | **Reaction ID** | **Proteins (Uniprot ID)** | **$k_{cat}$ ($s^{-1}$)** | **Source** |
|---|---|---|---|---|
| 3-Hydroxy-L-tyrosine carboxy-lyase (No1) | 3HLYTCLNo1 | Q88JU5 | 1.9 | (Koyanagi et al., 2012) |
| 4 hydroxyproline epimerase (reversible) (No1) | 4HPROE_REVNo1 | Q88NF3 | 7.74 | (Radkov and Moe, 2013) |
| 4 hydroxyproline epimerase (No1) | 4HPROENo1 | Q88NF3 | 78.3 | (Radkov and Moe, 2013) |
| Oxoglutarate dehydrogenase (lipoamide) (No1) | AKGDaNo1 | Q88FA9 | 39.77 | (Bunik et al., 2000) |
| Oxoglutarate dehydrogenase (dihydrolipoamide S-succinyltransferase) (No1) | AKGDbNo1 | Q88FB0 | 39.77 | (Bunik et al., 2000) |
| Alanine racemase (reversible) (No1) | ALAR_REVNo1 | Q88CB2 | 40.14 | (Radkov and Moe, 2013) |
| Alanine racemase (reversible) (No2) | ALAR_REVNo2 | Q88GJ9 | 8.83 | (Radkov and Moe, 2013) |

| Reaction name | Reaction ID | Proteins (Uniprot ID) | $k_{cat}$ ($s^{-1}$) | Source |
|---|---|---|---|---|
| Alanine racemase (No1) | ALARNo1 | Q88CB2 | 103.6 | (Radkov and Moe, 2013) |
| Alanine racemase (No2) | ALARNo2 | Q88GJ9 | 7.33 | (Radkov and Moe, 2013) |
| Alcohol dehydrogenase | ALCDppNo1 | Q88JH5 | 6882 | (Wehrmann and Klebensberger, 2018) |
| Arginine deiminase (No1) | ARGDINo1 | Q88P52 | 74.65 | (Patil et al., 2018) |
| Arginine deiminase (No1) | ARGDrNo1 | Q88P52 | 74.65 | (Patil et al., 2018) |
| Arginine racemase (reversible) (No1) | ARGR_REVNo1 | Q88GJ9 | 250.63 | (Radkov and Moe, 2013) |
| Arginine racemase (No1) | ARGRNo1 | Q88GJ9 | 213.47 | (Radkov and Moe, 2013) |
| Aspartate transaminase (reversible) (No1) | ASPTA_REVNo1 | Q88GK0 | 31.95 | (Yagi et al., 1976) |
| Aspartate transaminase (reversible) (No2) | ASPTA_REVNo2 | Q88LG1 | 32.58 | (Yagi et al., 1976) |
| ATP synthase (four protons for one ATP) (periplasm) (No1) | ATPS4rppNo1 | Q88BX0, Q88BX1, Q88BX2, Q88BX3, Q88BX4, Q88BX5, Q88BW8, Q88BW9 | 21.2 | (Iino et al., 2009) |
| Ubiquinol cytochrome c reductase (No1) | CYO1_KTNo1 | Q88N93, Q88N94, Q88N95 | 919 | (Dadak and Holik, 2008) |
| Cytochrome oxidase bd (ubiquinol-8: 2 protons) (periplasm) (No1) | CYTBDppNo1 | Q88E17, Q88E18 | 279.77 | (Jünemann and Wrigglesworth, 1995) |
| Cytochrome oxidase bo3 (ubiquinol-8: 4 protons) (periplasm) (No1) | CYTBO3_4ppNo1 | Q88PN4, Q88PN5, Q88PN6, Q88PN7 | 1052 | (Ding et al., 2019) |
| Cytochrome c oxidase aa3 2 protons1 electron periplasm (No1) | CYTCAA3ppNo1 | Q88RM3, Q88RM5, Q88RM6 | 91 | (Fukumori et al., 1985) |
| Cytochrome c oxidase bb3 05 protons1 electron periplasm (No1) | CYTCBB3ppNo1 | Q88F46, Q88F47, Q88F48, Q88F49 | 230 | (Urbani et al., 2001) |

| Reaction name | Reaction ID | Proteins (Uniprot ID) | $k_{cat}$ ($s^{-1}$) | Source |
|---|---|---|---|---|
| Cytochrome c oxidase bb3 05 protons1 electron periplasm (No2) | CYTCBB3ppNo2 | Q88F41, Q88F42, Q88F43, Q88F44 | 230 | (Urbani et al., 2001) |
| Asparagine racemase (reversible) (No1) | DLASNR_REVNo1 | Q88GJ9 | 2.9 | (Radkov and Moe, 2013) |
| Asparagine racemase (No1) | DLASNRNo1 | Q88GJ9 | 13.46 | (Radkov and Moe, 2013) |
| Cystein racemase unidirectional (No1) | DLCYSRNo1 | Q88CB2 | 3.43 | (Radkov and Moe, 2013) |
| Methionine racemase (reversible) (No1) | DLMETR_REVNo1 | Q88GJ9 | 33.07 | (Radkov and Moe, 2013) |
| Methionine racemase (No1) | DLMETRNo1 | Q88GJ9 | 121.85 | (Radkov and Moe, 2013) |
| 2-dehydro-3-deoxy-phosphogluconate aldolase (No1) | EDANo1 | Q88P29 | 260.46 | (Taha and Deits, 1994) |
| 6-phosphogluconate dehydratase (No1) | EDDNo1 | Q88P43 | 266.89 | (Scopes and Griffiths-Smith, 1984) |
| Fructose-bisphosphate aldolase (reversible) (No1) | FBA_REVNo1 | Q88D67 | 12.6 | (Stolzenberger et al., 2013) |
| Fructose-bisphosphate aldolase (No1) | FBANo1 | Q88D67 | 1.53 | (Banerjee et al., 1985) |
| Fructose-bisphosphatase (No1) | FBPNo1 | Q88CY9 | 39.51 | (Lynch et al., 1975) |
| Glucose 6-phosphate dehydrogenase (No1) | G6PDH2r_NADNo1 | Q88P31 | 279 | (Olavarria et al., 2015) |
| Glucose 6-phosphate dehydrogenase (No2) | G6PDH2r_NADNo2 | Q88C32 | 121 | (Volke et al., 2021) |
| Glucose 6-phosphate dehydrogenase (No3) | G6PDH2r_NADNo3 | Q88FP7 | 0.81 | (Volke et al., 2021) |
| Glucose 6-phosphate dehydrogenase (No1) | G6PDH2r_NADPNo1 | Q88P31 | 103 | (Olavarria et al., 2015) |
| Glucose 6-phosphate dehydrogenase (No2) | G6PDH2r_NADPNo2 | Q88C32 | 114 | (Volke et al., 2021) |
| Glucose 6-phosphate dehydrogenase (No3) | G6PDH2r_NADPNo3 | Q88FP7 | 0.56 | (Volke et al., 2021) |
| Gluconate 2 dehydrogenase periplasm (No1) | GAD2ktppNo1 | Q88HH4, Q88HH5, Q88HH6 | 300.39 | (Lynch et al., 1975) |
| galactouronate dehydrogenase | GALURDHNo1 | Q88NN6 | 33 | (Yoon et al., 2009) |

| Reaction name | Reaction ID | Proteins (Uniprot ID) | $k_{cat}$ ($s^{-1}$) | Source |
|---|---|---|---|---|
| Glucose dehydrogenase (ubiquinone-8 as acceptor) (periplasm) (No1) | GLCDppNo1 | Q88MX4 | 3193.51 | (Lynch et al., 1975) |
| glucuronate dehydrogenase | GLCURDHNo1 | Q88NN6 | 55 | (Yoon et al., 2009) |
| Glutamine racemase (reversible) (No1) | GLNR_REVNo1 | Q88GJ9 | 19.77 | (Radkov and Moe, 2013) |
| Glutamine racemase (No1) | GLNRNo1 | Q88GJ9 | 57.88 | (Radkov and Moe, 2013) |
| Glutarate hydroxylase (No1) | GLUTARHNo1 | Q88IU0 | 1380.73 | (Zhang et al., 2018) |
| Glycol dehydrogenase forward rxn ethanol acetaldehyde (No1) | GLYCOLDHppNo1 | Q88JH5 | 0.46 | (Wehrmann et al., 2020) |
| Glycol dehydrogenase forward rxn ethanol acetaldehyde (No2) | GLYCOLDHppNo2 | Q88JH0 | 1.08 | (Wehrmann et al., 2020) |
| Phosphogluconate dehydrogenase (No1) | GNDNo1 | Q88FP6 | 71.32 | (Stournaras et al., 1983) |
| Gluconokinase (No1) | GNKNo1 | Q88HE4 | 54.36 | (Lynch et al., 1975) |
| Hexokinase (D-glucose:ATP) (No1) | HEX1No1 | Q88P42 | 640.46 | (Lynch et al., 1975) |
| Histidine racemase (reversible) (No1) | HISR_REVNo1 | Q88GJ9 | 2.92 | (Radkov and Moe, 2013) |
| Histidine racemase (No1) | HISRNo1 | Q88GJ9 | 10.69 | (Radkov and Moe, 2013) |
| Isocitrate lyase (No1) | ICLNo1 | Q88FI0 | 9.81 | (Campos-Garcia et al., 2013) |
| Leucine racemase (reversible) (No1) | LEUR_REVNo1 | Q88GJ9 | 4.08 | (Radkov and Moe, 2013) |
| Leucine racemase (No1) | LEURNo1 | Q88GJ9 | 10.88 | (Radkov and Moe, 2013) |
| Lysine racemase (reversible) (No1) | LYSRC_REVNo1 | Q88GJ9 | 281.2 | (Radkov and Moe, 2013) |
| Lysine racemase (No1) | LYSRCNo1 | Q88GJ9 | 1843.4 | (Radkov and Moe, 2013) |
| Malate synthase (No1) | MALSNo1 | Q88QX8 | 21.55 | (Roucourt et al., 2009) |
| Malate dehydrogenase (ubiquinone 8 as acceptor) (No1) | MDH2No1 | Q88PU7 | 7.35 | (Oh et al., 2020) |

| Reaction name | Reaction ID | Proteins (Uniprot ID) | $k_{cat}$ ($s^{-1}$) | Source |
|---|---|---|---|---|
| NADH dehydrogenase (ubiquinone-8 & 3 protons) (periplasm) (No1) | NADH16ppNo1 | Q88FG5, Q88FG6, Q88FG7, Q88FG8, Q88FG9, Q88FH0, Q88FH1, Q88FH2, Q88FH3, Q88FH4, Q88FH5, Q88FH6, Q88FH7 | 2032.23 | (Narayanan et al., 2013) |
| NADH dehydrogenase (ubiquinone-8) (No1) | NADH5No1 | Q88Q70 | 872 | (Lencina et al., 2018) |
| NADH dehydrogenase (ubiquinone-8) (No2) | NADH5No2 | Q88GK1 | 872 | (Lencina et al., 2018) |
| Oxaloacetate decarboxylase (No1) | OAADCNo1 | Q88P29 | 7500 | (Narayanan et al., 2008) |
| Pyruvate carboxylase (No1) | PCNo1 | Q88C36, Q88C37 | 16.82 | (Choi et al., 2016) |
| Pyruvate dehydrogenase (lipoamide) (No1) | PDHaNo1 | Q88QZ5 | 30.2 | (Nemeria et al., 2001) |
| Pyruvate dehydrogenase (dihydrolipoamide) reversible (No1) | PDHbrNo1 | Q88QZ6 | 78 | (Arjunan et al., 2014) |
| Pyruvate dehydrogenase (dihydrolipoamide dehydrogenase) reversible (No1) | PDHcrNo1 | Q88C17 | 420 | (Sahlman and Williams, 1989) |
| Pyruvate dehydrogenase (dihydrolipoamide dehydrogenase) reversible (No2) | PDHcrNo2 | Q88FB1 | 420 | (Sahlman and Williams, 1989) |
| PhenylethylalcoholPQQ oxidoreductase (No1) | PEAOppNo1 | Q88JH5 | 9.68 | (Wehrmann et al., 2020) |
| PhenylethylalcoholPQQ oxidoreductase (No2) | PEAOppNo2 | Q88JH0 | 7.14 | (Wehrmann et al., 2020) |

| Reaction name | Reaction ID | Proteins (Uniprot ID) | $k_{cat}$ ($s^{-1}$) | Source |
|---|---|---|---|---|
| Glucose-6-phosphate isomerase (No1) | PGINo1 | Q88LW9 | 682.15 | (Lynch et al., 1975) |
| Glucose-6-phosphate isomerase (No2) | PGINo2 | Q88DW7 | 682.15 | (Lynch et al., 1975) |
| Phosphoglycerate kinase (No1) | PGKNo1 | Q88D64 | 537 | (Bentahir et al., 2000) |
| Phosphoribosyl-ATP pyrophosphatase (No1) | PRATPPNo1 | Q88D14 | 68.38 | (Keesey et al., 1979) |

**Table S6.** Analysis of different cellular objectives with enzyme constrained and protein constrained models. The lower the normalized Euclidean distance is the better the objective reflects the real distribution of fluxes.

| Condition | Normalized Euclidean distance ec model[a] | Normalized Euclidean distance pc model[b] |
|---|---|---|
| Maximized flux through biomass reaction[c] | 0.0966 | 0.2293 |
| Maximized ATP production through ATP synthase | 0.1187 | 0.2291 |
| Minimized ATP production through ATP synthase | 0.1394 | 0.2293 |
| Maximized ATP production rate | 0.1060 | 0.2293 |
| Minimized ATP production rate | 0.1151 | 0.2293 |
| Maximized $CO_2$ production | 0.0961 | 0.2293 |
| Minimized $CO_2$ production | 0.1030 | 0.2291 |
| Maximized $O_2$ production | 0.1108 | 0.4385 |
| Minimized $O_2$ production | 0.1211 | 0.2292 |
| Maximized glucose uptake rate | 0.0968 | 0.2291 |
| Minimized glucose uptake rate | 0.1033 | 0.2292 |
| Minimized sum of all fluxes | 0.1055 | 0.2293 |
| Maximized flux through NADH dehydrogenase | 0.1483 | 0.2293 |
| Minimized flux through NADH dehydrogenase | 0.1074 | 0.2293 |
| Maximized NADH production rate | 0.0960 | 0.2293 |
| Minimized NADH production rate | 0.1088 | 0.2293 |
| Maximized NADPH production rate | 0.0948 | 0.2293 |
| Minimized NADPH production rate | 0.1235 | 0.2292 |
| Maximize proteome pool investment | ND | 0.2293 |
| Minimize proteome pool investment | ND | 0.2293 |

[a]Wherever applicable the growth rate was constrained to 0.678 $h^{-1}$ which is the maximal growth rate that can be achieved in ec model.

[b]Wherever applicable the growth rate was constrained to 0.4605 $h^{-1}$ which is the maximal growth rate when oxygen uptake is constrained with upper bound 30 mmol/$g_{CDW}$/h.
[c]Growth rate was left unconstrained.

#### Supplementary figures

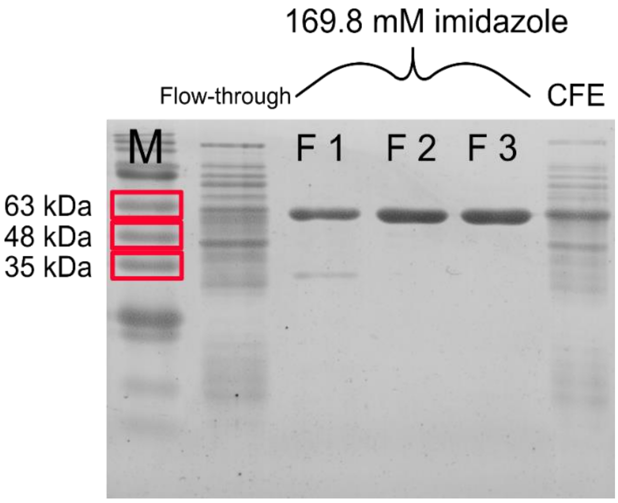


**Figure S1.** SDS-polyacrylamide gel with samples from the purification of BglC. Letter M designates marker (three bands with their corresponding molecular weight are highlighted) and Fx stands for collected elution fraction number x. Lines designated 169.8 mM imidazole contain 2.5 µg of protein. The remaining two lines (Flow-through and CFE – cell-free extract) contain 5 µg of protein. Major bands visible in the lines designated 169.8 mM of imidazole (concentration of imidazole in which BglC was eluted) correspond well with the theoretical molecular weight of BglC (53.4 kDa).

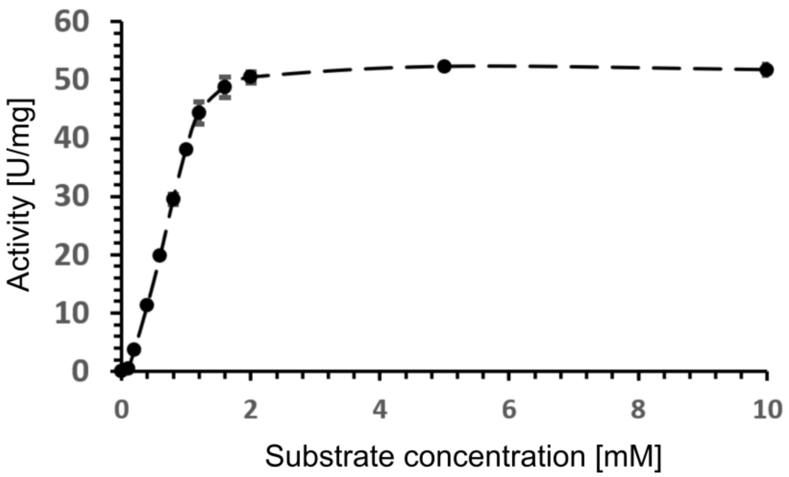


**Figure S2.** Michaelis & Menten plot for enzyme BglC with substrate 4-nitrophenyl β-D-glucopyranoside. The data points are means ± SD calculated from three biological replicates.

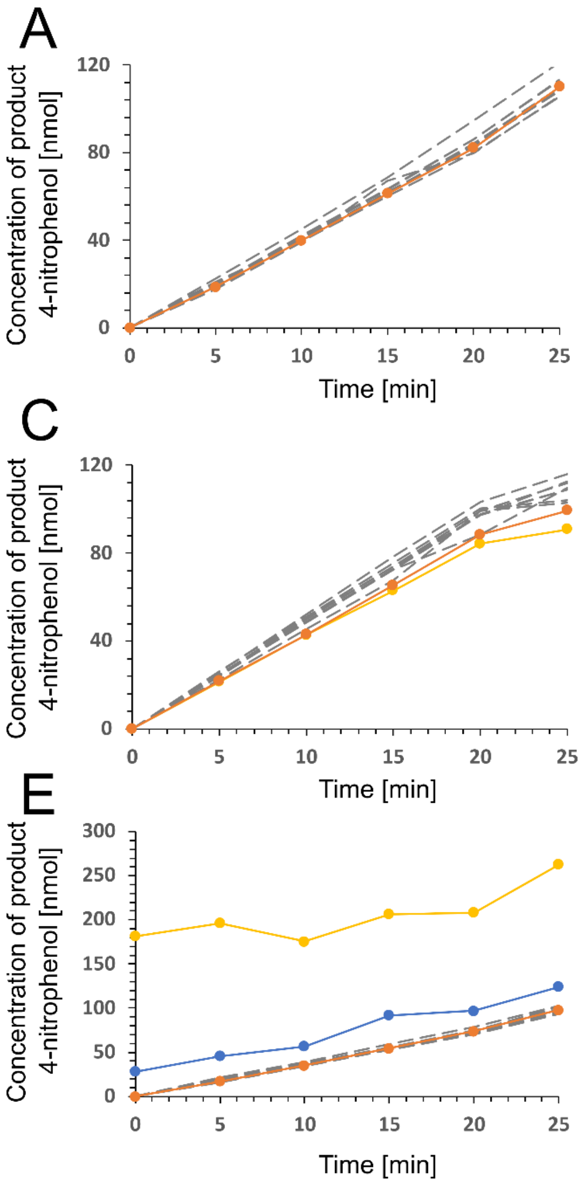


**Figure S3.** Screening of possible inhibitors of enzyme BglC. The curve for D-glucono-δ-lactone is not shown because the enzyme was completely inhibited at all concentrations tested. In total, six inhibitor concentrations were tested. (**A**) D-xylose, (**B**) D-gluconate, (**C**) D-glucose, (**D**) L-arabinose, (**E**) 2-ketogluconate.

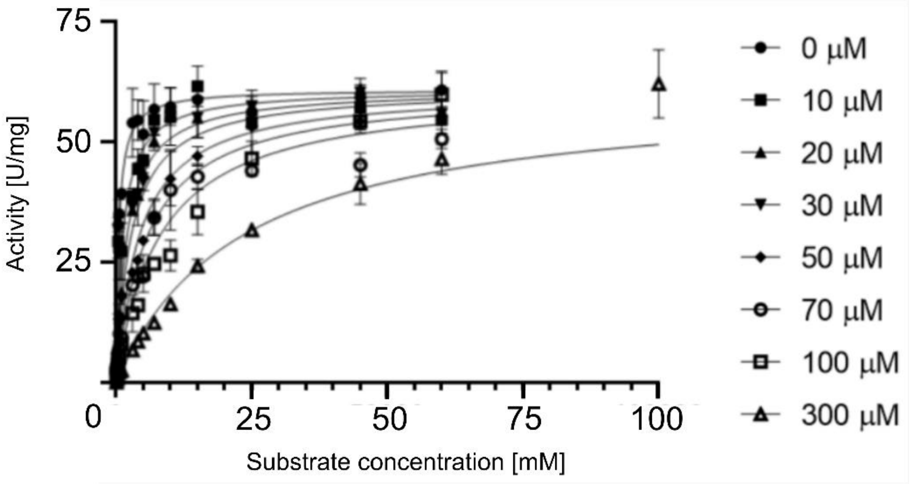


**Figure S4.** Measurement of BglC activity with substrate 4-nitrophenyl β-D-glucopyranoside in presence of various concentrations of D-glucono-δ-lactone. The data points are means ± SD calculated from at least three biological replicates.

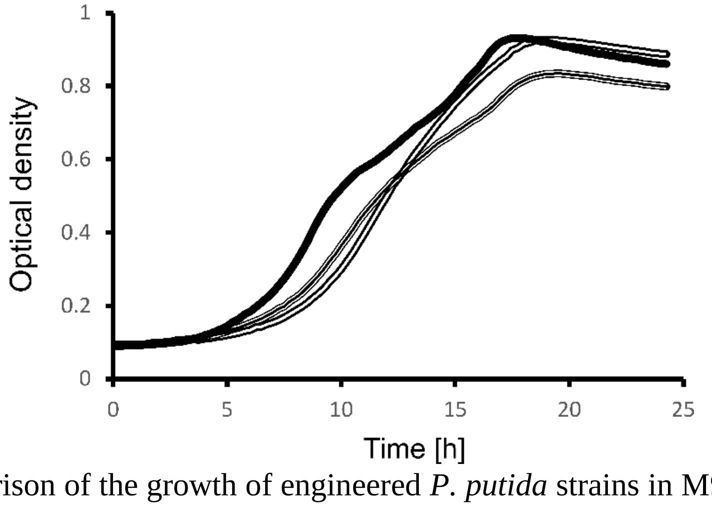


**Figure S5.** Comparison of the growth of engineered *P. putida* strains in M9 minimal medium with glucose (4 g/L) in 48-well plate format. GCDbglC_glf (*glf* inserted in chromosome), full line (▮); GCDbglC_*glf* (*glf* expressed from plasmid pSEVA2213_*bglC_glf*), half-full line (▯▮▯); GCDbglC, hollow line (▯).

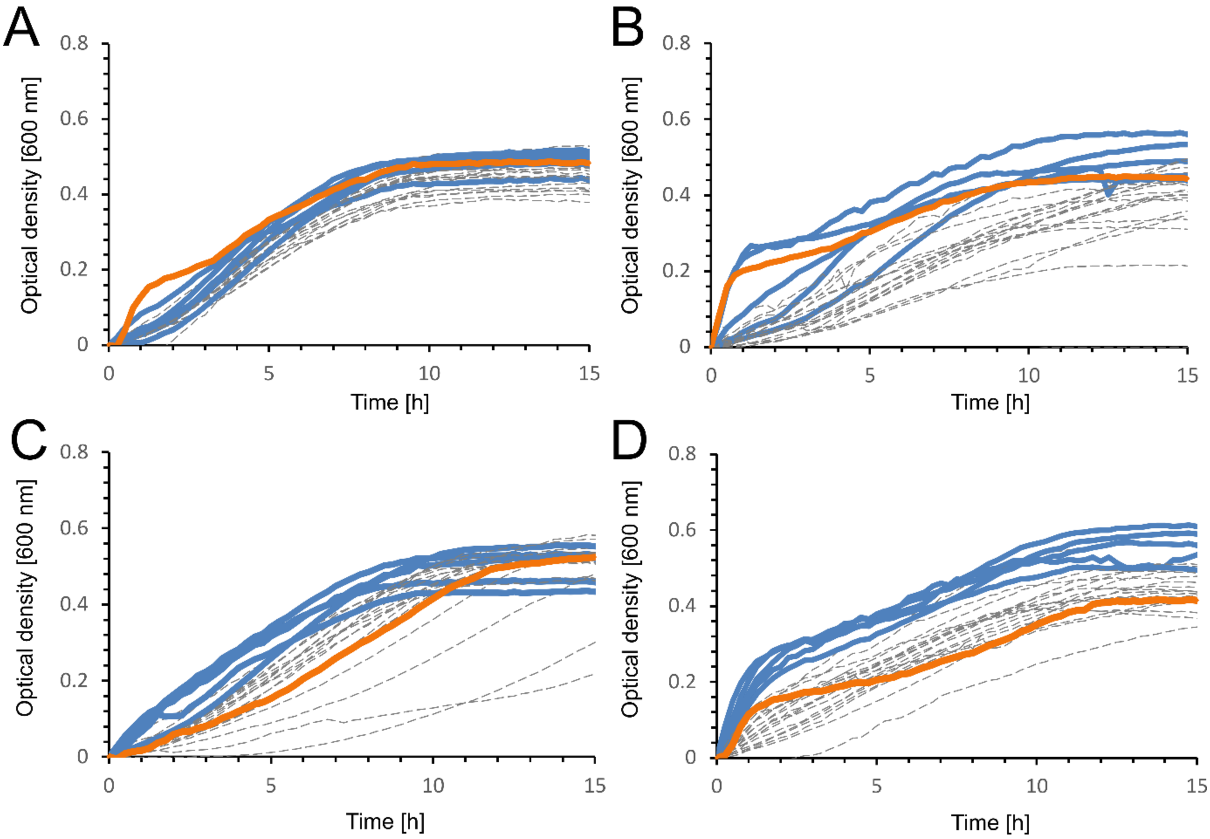


**Figure S6.** The first round of screening of *glf* mutants in 96-well plate format. The 23 new mutant strains for each parental strain were tested and then the five fastest-growing mutants were selected for the second round of screening (Figure S6). Parental strains (controls) were (**A**) WT, (**B**) hexWT, (**C**) GCD, (**D**) hexGCD. The initial optical density was subtracted from the growth profiles.

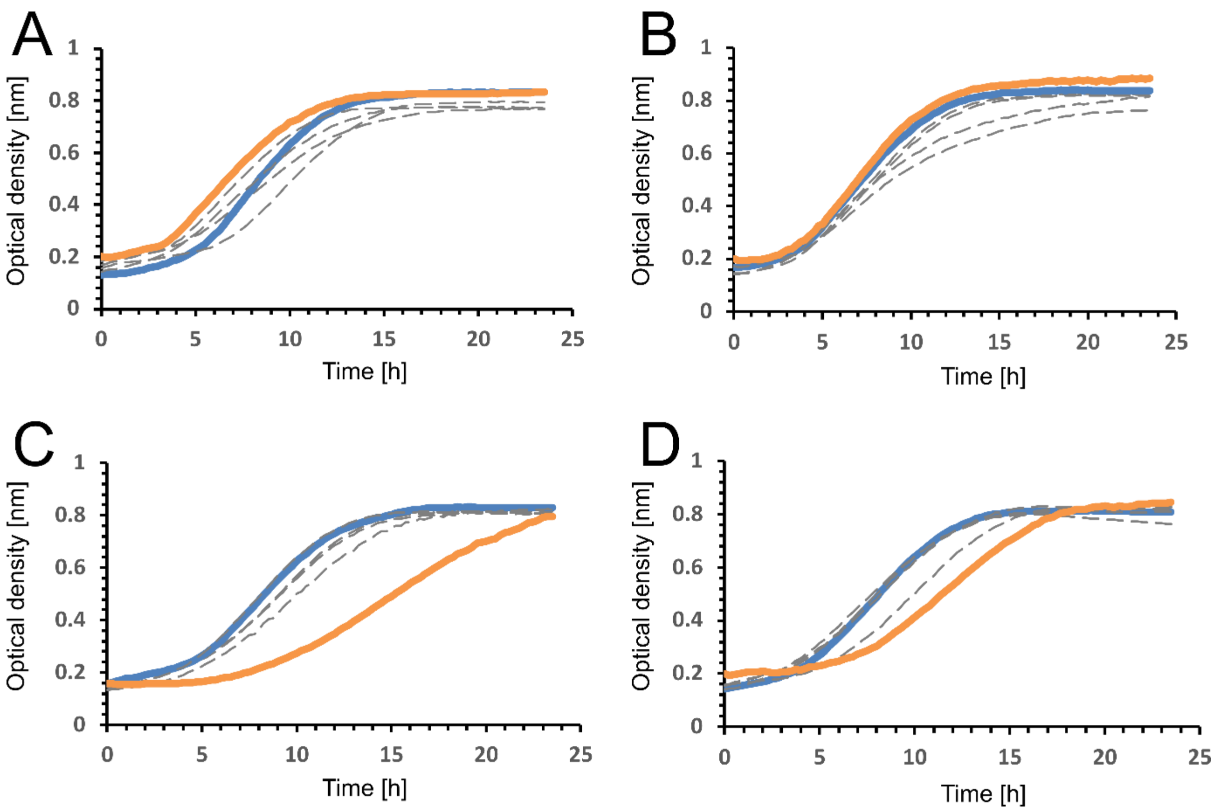


**Figure S7.** The second round of screening of *glf* mutants in 96-well plate format. Five best mutants from the first round (Figure S5) for each template strain were tested and the fastest-growing mutant was selected for further work. Parental strains (controls) were (**A**) WT, (**B**) hexWT, (**C**) GCD, (**D**) hexGCD.

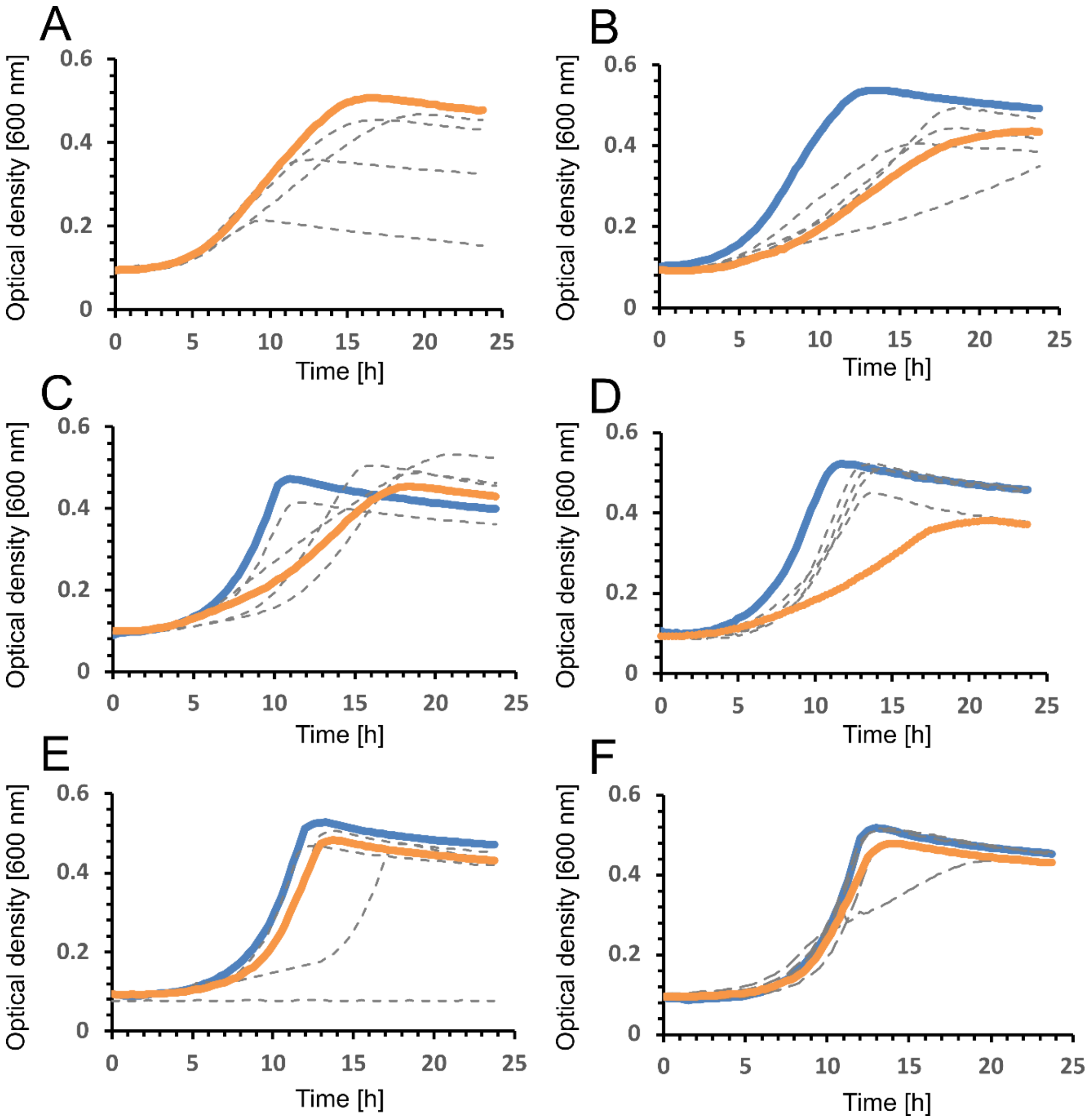


**Figure S8.** Screening of the library of *lacY* insertion mutants in 48 well-plate format. Strains were grown in M9 medium supplemented with cellobiose (2 g/L). Parental strains (controls) were (**A**) WTbglC, (**B**) WTbglC_glf, (**C**) hexWTbglC, (**D**) hexWTbglC_glf, (**E**) hexGCDbglC_glf, (**F**) GCDbglC_glf.

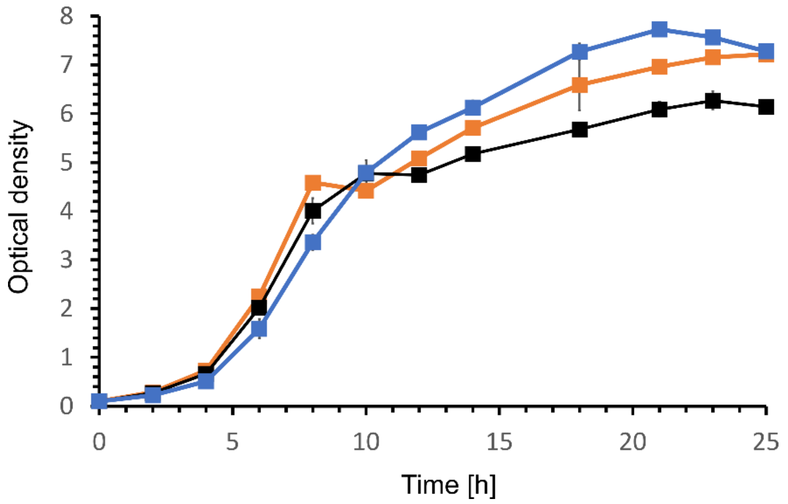


**Figure S9.** Comparison of the growth of three engineered *P. putida* strains in M9 minimal medium with the mixture of glucose (4 g/L) and cellobiose (4 g/L) in shake flasks. Strain hexWTbglC_glf_lacY (blue), strain hexWTbglC_lacY (orange), strain WTbglC_glf_lacY. The data points are means ± SD calculated from three biological replicates.

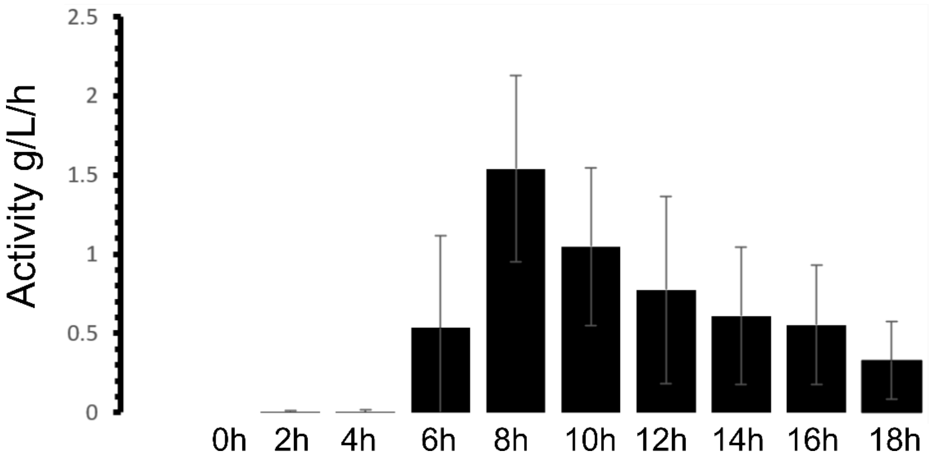


**Figure S10.** Cellobiohydrolytic activity in culture supernatants collected at multiple time points during shake flask culture of strain GCDbglC_glf_lacY on the mixture of glucose and cellobiose (4 g/L each). The columns show means ± SD from two biological replicates.

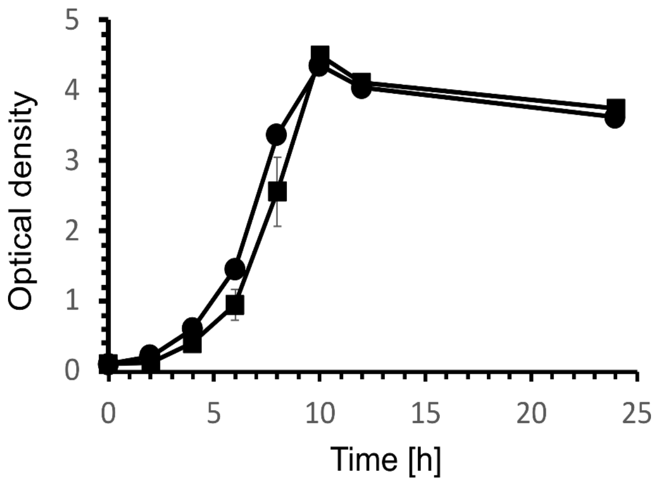


**Figure S11**. Comparison of the growth of HexGCDglf (filled circles ●) and hexGCDglf_colony2 (filled squares ■) in M9 minimal medium with glucose (4 g/L) in shake flasks. The data points are means ± SD calculated from three biological replicates.

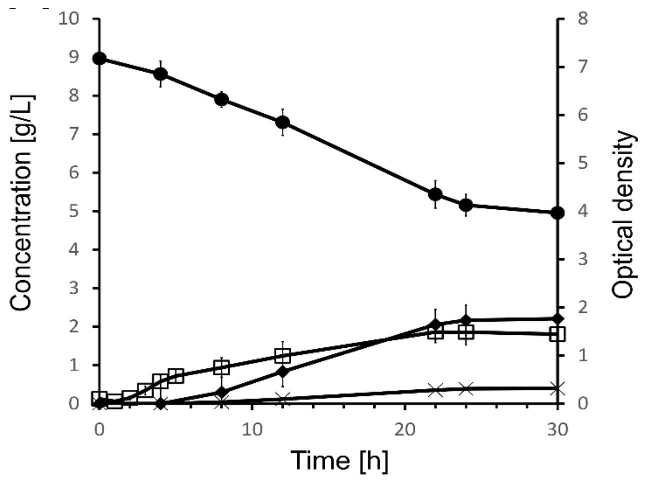


**Figure S12**. Shake flask culture of GCDglf colony 2 in M9 minimal medium with 8 g/L glucose. Cell growth, open squares (□); glucose, filled circles (●); pyruvate, filled diamonds (♦); acetate, stars (×). The data points are means ± SD calculated from three biological replicates.

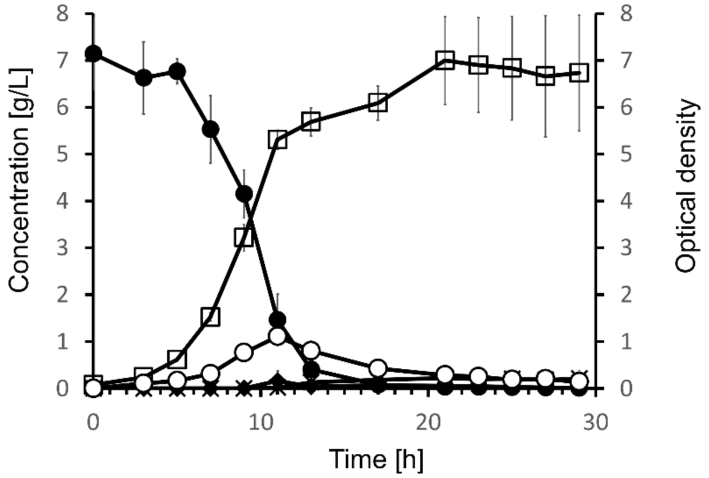


**Figure S13.** Shake flask culture of WTbglC in M9 minimal medium with 8 g/L glucose. Cell growth, open squares (□); glucose, filled circles (●); gluconate, open circles (○); pyruvate, filled diamonds (♦); acetate, stars (×). Only traces of pyruvate, acetate, and 2-ketogluconate were detected. The data points are means ± SD calculated from four biological replicates.

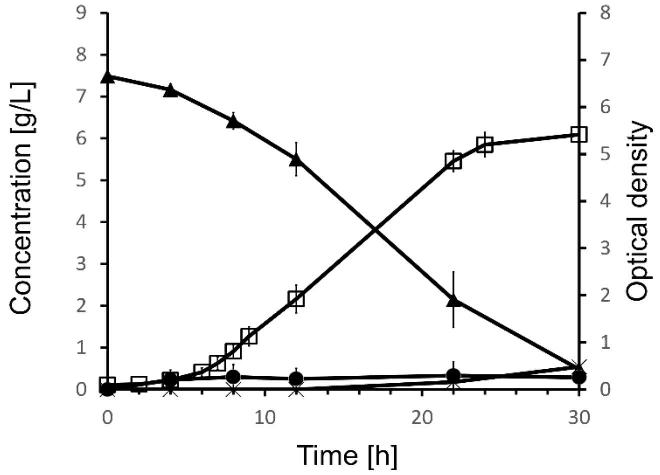


**Figure S14**. Shake flask culture of GCDbglC_glf_lacY in M9 minimal medium with 8 g/L cellobiose. Cell growth, open squares (□); cellobiose, filled triangles (▲); glucose, filled circles (●); pyruvate, filled diamonds (♦); acetate, stars (×). The data points are means ± SD calculated from three biological replicates.

#### Nucleotide sequences of genes used in this study

##### Gene *glf* from *Zymomonas mobilis*

ATGAGTTCTGAAAGTAGTCAGGGTCTAGTCACGCGACTAGCCCTAATCGCTGCTATAGGCGG
CTTGCTTTTCGGTTACGATTCAGCGGTTATCGCTGCAATCGGTACACCGGTTGATATCCATT
TTATTGCCCCTCGTCACCTGTCTGCTACGGCTGCGGCTTCCCTTTCTGGGATGGTCGTTGTT
GCTGTTTTGGTCGGTTGTGTTACCGGTTCTTTGCTGTCTGGCTGGATTGGTATTCGCTTCGG
TCGTCGCGGCGGATTGTTGATGAGTTCCATTTGTTTCGTCGCCGCCGGTTTTGGTGCTGCGT
TAACCGAAAAATTATTTGGAACCGGTGGTTCGGCTTTACAAATTTTTTGCTTTTTCCGGTTT
CTTGCCGGTTTAGGTATCGGTGTCGTTTCAACCTTGACCCCAACCTATATTGCTGAAATTGC
TCCGCCAGACAAACGTGGTCAGATGGTTTCTGGTCAGCAGATGGCCATTGTGACGGGTGCTT
TAACCGGTTATATCTTTACCTGGTTACTGGCTCATTTCGGTTCTATCGATTGGGTTAATGCC
AGTGGTTGGTGCTGGTCTCCGGCTTCAGAAGGCCTGATCGGTATTGCCTTCTTATTGCTGCT
GTTAACCGCACCGGATACGCCGCATTGGTTGGTGATGAAGGGACGTCATTCCGAGGCTAGCA
AAATCCTTGCTCGTCTGGAACCGCAAGCCGATCCTAATCTGACGATTCAAAAGATTAAAGCT
GGCTTTGATAAAGCCATGGACAAAAGCAGCGCAGGTTTGTTTGCTTTTGGTATCACCGTTGT
TTTTGCCGGGGTATCCGTTGCTGCCTTCCAGCAGTTGGTCGGTATTAACGCCGTGCTGTATT
ATGCACCGCAGATGTTCCAGAATTTAGGTTTTGGAGCTGATACGGCATTATTGCAGACCATC
TCTATCGGTGTTGTGAACTTCATCTTCACCATGATTGCTTCCCGTGTTGTTGACCGCTTCGG
CCGTAAACCTCTGCTTATTTGGGGTGCTCTCGGTATGGCTGCAATGATGGCTGTTTTAGGCT
GCTGTTTCTGGTTCAAAGTCGGTGGTGTTTTGCCTTTGGCTTCTGTGCTTCTTTATATTGCA
GTCTTTGGCATGTCATGGGGCCCTGTCTGCTGGGTTGTTCTGTCAGAAATGTTCCCGAGTTC
CATCAAGGGCGCAGCTATGCCTATCGCTGTTACCGGACAATGGTTAGCTAATATCTTGGTTA
ACTTCCTGTTTAAGGTTGCTGATGGTTCTCCAGCATTGAATCAGACTTTCAACCACGGTTTC
TCCTATCTCGTTTTCGCAGCATTAAGTATCTTAGGTGGCTTGATTGTTGCTCGCTTCGTGCC
GGAAACCAAAGGTCGGAGCCTGGATGAAATCGAGGAGATGTGGCGCTCCCAGAAGTAA

**Gene *lacY* from *Echerichia coli***

ATGTACTATTTAAAAAACACAAACTTTTGGATGTTCGGTTTATTCTTTTTCTTTTACTTTTT
TATCATGGGAGCCTACTTCCCGTTTTTCCCGATTTGGCTACATGACATCAACCATATCAGCA
AAAGTGATACGGGTATTATTTTTGCCGCTATTTCTCTGTTCTCGCTATTATTCCAACCGCTG
TTTGGTCTGCTTTCTGACAAACTCGGGCTGCGCAAATACCTGCTGTGGATTATTACCGGCAT
GTTAGTGATGTTTGCGCCGTTCTTTATTTTTATCTTCGGGCCACTGTTACAATACAACATTT
TAGTAGGATCGATTGTTGGTGGTATTTATCTAGGCTTTTGTTTTAACGCCGGTGCGCCAGCA
GTAGAGGCATTTATTGAGAAAGTCAGCCGTCGCAGTAATTTCGAATTTGGTCGCGCGCGGAT
GTTTGGCTGTGTTGGCTGGGCGCTGTGTGCCTCGATTGTCGGCATCATGTTCACCATCAATA
ATCAGTTTGTTTTCTGGCTGGGCTCTGGCTGTGCACTCATCCTCGCCGTTTTACTCTTTTTC
GCCAAAACGGATGCGCCCTCTTCTGCCACGGTTGCCAATGCGGTAGGTGCCAACCATTCGGC
ATTTAGCCTTAAGCTGGCACTGGAACTGTTCAGACAGCCAAAACTGTGGTTTTTGTCACTGT
ATGTTATTGGCGTTTCCTGCACCTACGATGTTTTTGACCAACAGTTTGCTAATTTCTTTACT
TCGTTCTTTGCTACCGGTGAACAGGGTACGCGGGTATTTGGCTACGTAACGACAATGGGCGA
ATTACTTAACGCCTCGATTATGTTCTTTGCGCCACTGATCATTAATCGCATCGGTGGGAAAA
ACGCCCTGCTGCTGGCTGGCACTATTATGTCTGTACGTATTATTGGCTCATCGTTCGCCACC
TCAGCGCTGGAAGTGGTTATTCTGAAAACGCTGCATATGTTTGAAGTACCGTTCCTGCTGGT
GGGCTGCTTTAAATATATTACCAGCCAGTTTGAAGTGCGTTTTTCAGCGACGATTTATCTGG
TCTGTTTCTGCTTCTTTAAGCAACTGGCGATGATTTTTATGTCTGTACTGGCGGGCAATATG
TATGAAAGCATCGGTTTCCAGGGCGCTTATCTGGTGCTGGGTCTGGTGGCGCTGGGCTTCAC
CTTAATTTCCGTGTTCACGCTTAGCGGCCCCGGTCCGCTTTCTCTACTGCGTCGTCAGGTGA
ATGAAGTCGCTTAA

**Synthetic *bglC_adhB_pdc* operon with genes encoding β-glucosidase BglC from *Thermobifida fusca* and alcohol dehydrogenase AdhB and pyruvate decarboxylase from *Zymomonas mobilis*, respectively. The latter two genes were codon optimized for expression in *P. putida* KT2440. Sequences of constitutive EM7 promoter, synthetic ribosom binding sites, and T0 terminator are included.**

TGTTGACAATTAATCATCGGCATAGTATATCGGCATAGTATAATACGACAAGGTGAGGAACT
AAACCCCTAGGCCGCGGCCGCGCGAATTCGAGCTCGGTACCCTTTAAGAAGGAGATATACAT
ATGCACCATCACCATCACCATACCTCGCAATCGACGACTCCTCTGGGCAATCTCGAGGAGAC
TCCCAAACCGGATATCCGCTTCCCGTCCGATTTCGTGTGGGGAGTGGCGACCGCTTCGTTCC

AGATCGAAGGCTCCACCACGGCCGACGGCCGCGGCCCCAGCATCTGGGACACCTTCTGCGCC
ACTCCGGGCAAGGTCGAGAACGGCGACACGGGCGACCCTGCCTGCGACCACTACAACCGGTA
CCGCGATGACGTGGCCTTGATGCGGGAGCTGGGCGTGGGCGCCTACCGCTTCTCCATCGCCT
GGCCGCGGATCCAGCCCGAGGGCAAGGGCACGCCCGTGGAGGCCGGGCTGGACTTCTACGAC
CGGCTTGTGGACTGCCTGCTGGAGGCCGGCATCGAGCCGTGGCCGACCCTCTACCACTGGGA
CCTGCCGCAGGCGCTGGAGGACGCGGGCGGCTGGCCCAACCGGGACACGGCCAAGCGGTTCG
CCGACTACGCGGAGATCGTCTACCGCCGGCTCGGCGACCGGATCACCAACTGGAACACGCTC
AACGAGCCGTGGTGCTCCGCGTTCCTGGGCTACGCCTCCGGCGTGCACGCCCCGGGCCGCCA
GGAGCCGGCTGCTGCGCTGGCCGCCGCCCACCACCTGATGCTGGGCCACGGGCTGGCCGCTG
CCGTGATGCGGGACTTGGCGGGCCAGGCCGGACGTTCCGTGCGGATCGGTGTCGCGCACAAC
CAGACCACGGTCCGTCCCTACACTGACAGTGAGGCCGACCGGGACGCTGCGCGCCGGATTGA
CGCCCTGCGGAACCGCATCTTCACCGAGCCGCTGGTGAAGGGCCGCTACCCGGAGGACCTGA
TCGAGGACGTCGCCGCGGTCACCGACTACAGCTTCGTCCAGGACGGCGACCTGAAGACCATC
TCCGCCAACCTGGACATGATGGGCGTCAACTTCTACAACCCGAGCTGGGTGTCAGGCAACCG
GGAGAACGGGGGCTCCGACCGGCTGCCCGACGAGGGCTACTCGCCGTCGGTCGGCAGCGAGC
ATGTCGTGGAGGTGGACCCCGGCCTGCCGGTGACCGCCATGGGCTGGCCGATCGACCCGACC
GGGCTGTACGACACGCTGACCCGGCTGGCCAACGACTACCCGGGCCTGCCGCTGTACATCAC
CGAGAACGGCGCCGCCTTCGAGGACAAGGTGGTCGACGGCGCGGTGCACGACACCGAGCGGA
TCGCCTACCTGGACTCGCACCTGCGGGCCGCGCACGCTGCCATTGAGGCGGGCGTGCCGCTC
AAGGGCTACTTCGCCTGGTCGTTCATGGACAACTTCGAGTGGGCCCTCGGGTACGGGAAGCG
GTTCGGCATCGTGCACGTGGACTACGAGAGCCAGACGCGCACGGTGAAGGACAGCGGCTGGT
GGTACTCCCGGGTGATGCGCAACGGGGGAATCTTCGGACAGGAATAGCTGCAGGGGCCGCGA
CTGCGGCAACCCCTACGAAATGGCGATCGCCGCCGGTTAACTCACACAGGAGGGTATCATAT
GGCCAGCAGCACCTTCTACATCCCGTTCGTGAACGAGATGGGCGAGGGCAGCCTGGAAAAGG
CCATCAAGGACCTGAACGGCAGCGGCTTCAAGAACGCCCTGATCGTGAGCGACGCCTTCATG
AACAAGAGCGGTGTGGTGAAGCAAGTGGCCGACCTGCTGAAGGCCCAGGGCATCAACAGCGC
CGTGTACGATGGCGTGATGCCGAACCCGACCGTGACCGCCGTGCTGGAAGGCCTGAAGATCC
TGAAGGACAACAACAGCGACTTCGTGATCAGCCTGGGTGGCGGTAGCCCACACGACTGCGCC
AAAGCGATCGCCCTGGTCGCCACCAACGGTGGCGAGGTGAAGGACTACGAGGGCATCGACAA
GAGCAAGAAGCCAGCGCTGCCGCTGATGAGCATCAACACCACCGCCGGCACCGCCAGCGAGA
TGACCCGCTTCTGCATCATCACCGACGAGGTGCGCCACGTGAAGATGGCGATCGTGGACCGC
CACGTCACCCCGATGGTGAGCGTGAACGACCCGCTGCTGATGGTGGGTATGCCGAAGGGTCT
GACCGCCGCCACCGGCATGGATGCGCTGACCCATGCCTTCGAGGCCTACAGCTCGACCGCGG
CGACCCCGATCACCGATGCCTGCGCGCTGAAAGCCGCCAGCATGATCGCCAAGAACCTGAAA

ACCGCCTGCGACAACGGCAAGGACATGCCAGCGCGTGAGGCGATGGCGTACGCGCAGTTCCT
GGCCGGTATGGCCTTCAACAACGCCAGCCTGGGCTACGTCCATGCCATGGCGCACCAGCTGG
GTGGCTACTACAACCTGCCGCACGGCGTGTGCAACGCGGTCCTGCTGCCGCATGTGCTGGCC
TACAACGCCTCGGTGGTGGCCGGTCGCCTGAAGGATGTGGGCGTCGCCATGGGCCTGGATAT
CGCCAACCTGGGCGACAAAGAGGGTGCCGAGGCCACCATCCAGGCCGTGCGCGACCTGGCCG
CCTCGATCGGCATCCCAGCCAACCTGACCGAGCTGGGTGCCAAGAAAGAGGACGTCCCACTG
CTGGCCGACCACGCGCTGAAGGACGCGTGCGCCCTGACCAACCCACGCCAGGGCGACCAGAA
AGAGGTGGAAGAACTGTTCCTGAGCGCCTTCTGACACCAACCCGAACGGCAGCAAGAACACT
GCTATCAGAGGAGACGAAACATATGAGCTACACCGTGGGCACCTACCTGGCCGAGCGCCTGG
TGCAGATCGGCCTGAAGCACCACTTCGCCGTGGCCGGTGACTACAACCTGGTGCTGCTGGAT
AACCTGCTGCTGAACAAGAACATGGAACAGGTGTACTGCTGCAACGAGCTGAACTGCGGCTT
CAGCGCCGAGGGCTATGCCCGTGCCAAGGGTGCCGCCGCCGCGGTGGTGACCTACAGCGTGG
GTGCCCTGAGCGCCTTCGACGCCATCGGTGGTGCGTACGCCGAGAACCTGCCGGTGATCCTG
ATCAGCGGTGCCCCAAACAACAACGACCATGCCGCCGGTCACGTGCTGCATCACGCCCTGGG
CAAGACCGACTACCACTACCAGCTGGAAATGGCCAAGAACATCACCGCCGCGGCCGAGGCCA
TCTACACCCCAGAAGAAGCCCCAGCCAAGATCGACCACGTGATCAAGACCGCGCTGCGCGAA
AAGAAGCCGGTCTACCTGGAAATCGCCTGCAACATCGCCTCGATGCCGTGCGCCGCGCCAGG
TCCAGCGAGCGCCCTGTTCAACGACGAGGCCAGCGACGAAGCCAGCCTGAACGCCGCCGTGG
AAGAAACCCTGAAGTTCATCGCCAACCGCGACAAGGTGGCCGTGCTGGTCGGCAGCAAGCTG
CGTGCCGCGGGTGCCGAAGAAGCCGCCGTCAAATTCGCCGACGCGCTGGGTGGTGCCGTGGC
CACCATGGCCGCCGCCAAGAGCTTCTTCCCGGAAGAGAACCCGCACTACATCGGCACCAGCT
GGGGTGAAGTGAGCTACCCAGGCGTGGAAAAGACCATGAAAGAGGCCGACGCCGTCATCGCG
CTGGCCCCAGTGTTCAACGATTACAGCACCACCGGCTGGACCGACATCCCCGATCCGAAGAA
ACTGGTGCTGGCCGAACCGCGTAGCGTGGTGGTGAACGGCATCCGCTTCCCGAGCGTGCACC
TGAAGGACTATCTGACCCGTCTGGCCCAGAAGGTGAGCAAGAAAACCGGTGCGCTGGACTTC
TTCAAGTCGCTGAACGCGGGTGAGCTGAAGAAAGCGGCCCCAGCCGATCCGAGCGCGCCACT
GGTGAACGCCGAAATCGCCCGTCAGGTGGAAGCCCTGCTGACCCCGAACACCACCGTGATCG
CCGAAACCGGTGACAGCTGGTTCAACGCCCAGCGCATGAAGCTGCCGAACGGTGCCCGTGTG
GAATACGAGATGCAGTGGGGTCACATCGGCTGGTCCGTGCCAGCCGCCTTCGGCTACGCCGT
GGGTGCGCCAGAACGCCGTAACATCCTGATGGTCGGCGACGGCTCGTTCCAGCTGACCGCGC
AAGAGGTGGCCCAGATGGTCCGCCTGAAACTGCCCGTGATCATCTTCCTGATCAACAACTAC
GGCTACACCATCGAGGTGATGATCCACGACGGTCCGTACAACAACATCAAGAACTGGGACTA
TGCCGGTCTGATGGAAGTGTTCAACGGCAACGGTGGCTACGACAGCGGTGCCGGTAAGGGCC
TGAAGGCCAAGACCGGTGGCGAACTGGCCGAAGCCATCAAGGTCGCCCTGGCGAACACCGAC

GGTCCCACCCTGATCGAGTGCTTCATCGGTCGCGAGGACTGCACCGAGGAACTGGTGAAGTG
GGGCAAGCGTGTGGCCGCCGCGAACAGCCGTAAGCCGGTGAACAAGCTGCTGTGAAAGCTTG
CGGCCGCGTCGTGACTGGGAAAACCCTGGCGACTAGTCTTGGACTCCTGTTGATAGATCCAG
TAATGACCTCAGAACTCCATCTGGATTTGTTCAGAACGCTCGGTTGCCGCCGGGCGTTTTTT
ATTGGTGAGAATCCAG

**Synthetic *LDHA_bglC* operon with genes encoding codon-optimized N109G mutant of L-lactate dehydrogenase from *Bos taurus* and β-glucosidase BglC from *Thermobifida fusca*. Sequences of constitutive EM7 promoter, synthetic ribosom binding sites, and T0 terminator are included.**

TGTTGACAATTAATCATCGGCATAGTATATCGGCATAGTATAATACGACAAGGTGAGGAACT
AAACCCCTAGGCCGCGGCCGCGCGAATTCAGGAGGACAGCTATGGCCACCCTGAAGGACCAG
CTGATCCAGAACCTGCTGAAGGAAGAACACGTCCCGCAGAACAAGATCACCATCGTGGGCGT
GGGCGCCGTGGGCATGGCCTGCGCCATCAGCATCCTGATGAAGGACCTGGCCGACGAGGTGG
CCCTGGTGGACGTGATGGAGGACAAGCTGAAGGGCGAGATGATGGACCTGCAGCACGGCAGC
CTGTTCCTGCGCACCCCGAAGATCGTGAGCGGCAAGGACTACAACGTGACCGCCAACAGCCG
CCTGGTGATCATCACCGCCGGCGCCCGCCAGCAGGAAGGTGAAAGCCGCCTGGGCCTGGTGC
AGCGCAACGTGAACATCTTCAAGTTCATCATCCCGAACATCGTGAAGTACAGCCCGAACTGC
AAGCTGCTGGTCGTGAGCAACCCGGTGGACATCCTGACCTACGTGGCCTGGAAGATCAGCGG
CTTCCCGAAGAACCGCGTGATCGGCAGCGGCTGCAACCTGGACAGCGCCCGCTTCCGCTACC
TGATGGGCGAACGCCTGGGCGTGCACCCGCTGAGCTGCCACGGCTGGATTCTGGGCGAACAC
GGCGACAGCAGCGTGCCGGTGTGGAGCGGCGTGAACGTGGCCGGCGTGAGCCTGAAGAACCT
GCACCCGGAACTGGGCACCGACGCCGACAAGGAACAGTGGAAGGCCGTGCACAAGCAGGTGG
TGGACAGCGCCTACGAGGTGATCAAGCTGAAGGGCTACACCAGCTGGGCCATCGGCCTGAGC
GTGGCGGACCTGGCCGAGAGCATCATGAAGAACCTGCGCCGCGTGCACCCGATCAGCACCAT
GATCAAGGGCCTGTACGGTATCAAGGAAGACGTGTTCCTGAGCGTGCCGTGCATCCTGGGCC
AGAACGGCATCAGCGACGTGGTGAAGGTGACCCTGACCCACGAGGAAGAGGCCTGCCTGAAG
AAGAGCGCCGACACCCTGTGGGGCATCCAGAAGGAACTGCAGTTCTGAGAGCTCGGTACCCT
TTAAGAAGGAGATATACATATGCACCATCACCATCACCATACCTCGCAATCGACGACTCCTC
TGGGCAATCTCGAGGAGACTCCCAAACCGGATATCCGCTTCCCGTCCGATTTCGTGTGGGGA
GTGGCGACCGCTTCGTTCCAGATCGAAGGCTCCACCACGGCCGACGGCCGCGGCCCCAGCAT
CTGGGACACCTTCTGCGCCACTCCGGGCAAGGTCGAGAACGGCGACACGGGCGACCCTGCCT
GCGACCACTACAACCGGTACCGCGATGACGTGGCCTTGATGCGGGAGCTGGGCGTGGGCGCC

TACCGCTTCTCCATCGCCTGGCCGCGGATCCAGCCCGAGGGCAAGGGCACGCCCGTGGAGGC
CGGGCTGGACTTCTACGACCGGCTTGTGGACTGCCTGCTGGAGGCCGGCATCGAGCCGTGGC
CGACCCTCTACCACTGGGACCTGCCGCAGGCGCTGGAGGACGCGGGCGGCTGGCCCAACCGG
GACACGGCCAAGCGGTTCGCCGACTACGCGGAGATCGTCTACCGCCGGCTCGGCGACCGGAT
CACCAACTGGAACACGCTCAACGAGCCGTGGTGCTCCGCGTTCCTGGGCTACGCCTCCGGCG
TGCACGCCCCGGGCCGCCAGGAGCCGGCTGCTGCGCTGGCCGCCGCCCACCACCTGATGCTG
GGCCACGGGCTGGCCGCTGCCGTGATGCGGGACTTGGCGGGCCAGGCCGGACGTTCCGTGCG
GATCGGTGTCGCGCACAACCAGACCACGGTCCGTCCCTACACTGACAGTGAGGCCGACCGGG
ACGCTGCGCGCCGGATTGACGCCCTGCGGAACCGCATCTTCACCGAGCCGCTGGTGAAGGGC
CGCTACCCGGAGGACCTGATCGAGGACGTCGCCGCGGTCACCGACTACAGCTTCGTCCAGGA
CGGCGACCTGAAGACCATCTCCGCCAACCTGGACATGATGGGCGTCAACTTCTACAACCCGA
GCTGGGTGTCAGGCAACCGGGAGAACGGGGGCTCCGACCGGCTGCCCGACGAGGGCTACTCG
CCGTCGGTCGGCAGCGAGCATGTCGTGGAGGTGGACCCCGGCCTGCCGGTGACCGCCATGGG
CTGGCCGATCGACCCGACCGGGCTGTACGACACGCTGACCCGGCTGGCCAACGACTACCCGG
GCCTGCCGCTGTACATCACCGAGAACGGCGCCGCCTTCGAGGACAAGGTGGTCGACGGCGCG
GTGCACGACACCGAGCGGATCGCCTACCTGGACTCGCACCTGCGGGCCGCGCACGCTGCCAT
TGAGGCGGGCGTGCCGCTCAAGGGCTACTTCGCCTGGTCGTTCATGGACAACTTCGAGTGGG
CCCTCGGGTACGGGAAGCGGTTCGGCATCGTGCACGTGGACTACGAGAGCCAGACGCGCACG
GTGAAGGACAGCGGCTGGTGGTACTCCCGGGTGATGCGCAACGGGGGAATCTTCGGACAGGA
ATAGCTGCAGACATGCAAGCTTGCGGCCGCGTCGTGACTGGGAAAACCCTGGCGACTAGTCT
TGGACTCCTGTTGATAGATCCAGTAATGACCTCAGAACTCCATCTGGATTTGTTCAGAACGC
TCGGTTGCCGCCGGGCGTTTTTTATTGGTGAGAATCCAG
